# Transition Intermediates Encode Ligand Efficacy at the μ-Opioid Receptor-Gi Protein Complex

**DOI:** 10.64898/2026.09.02.748988

**Authors:** Kirill Konovalov, Sabina Novack, Davide Provasi, Vishal Maingi, Georgios Skiniotis, Marta Filizola

## Abstract

How ligands with different efficacies regulate G protein-coupled receptor signaling remains incompletely understood. Here, we combine Markov state modeling of nearly one millisecond of aggregate all-atom molecular dynamics simulations with time-resolved cryogenic-electron microscopy (cryo-EM) to reconstruct the most probable transition pathways connecting recently resolved structures of guanosine triphosphate (GTP)-bound μ-opioid receptor (MOR)–G_i1_ complexes with the full agonist lofentanil (LFT) or partial agonist mitragynine pseudoindoxyl (MP). The models predict four previously unresolved intermediate conformations. Guided by these predictions, reanalysis of the cryo-EM particle ensemble identifies conformational populations consistent with all four intermediates. Remarkably, the largest ligand-dependent structural differences emerge within these intermediates rather than in the previously resolved states. LFT favors a predominantly sequential and kinetically efficient transition pathway, whereas MP stabilizes a broader intermediate ensemble along this pathway and slows progression toward later states. Together, these complementary approaches provide an enriched thermodynamic and kinetic description of MOR-mediated G protein activation, revealing that opioid efficacy is encoded, at least in part, in the differential stabilization and kinetics of transition intermediates extending beyond GTP binding but well before complete G protein dissociation.

## Main

Opioid analgesics acting at the μ-opioid receptor (MOR), a member of the G protein-coupled receptor (GPCR) superfamily, remain the most effective treatments for acute, perioperative, cancer-related, and palliative pain, but their clinical use is limited by respiratory depression, tolerance, physical dependence, and abuse liability. These challenges have motivated diverse strategies to develop safer opioids, including G protein–biased agonists, low-efficacy (partial) agonists, peripherally restricted compounds, and orthosteric, allosteric, and bitopic ligands designed to exploit Gα subtype selectivity^1^. Despite the identification of several compounds with improved pharmacological profiles, no clinically approved MOR agonist fully dissociates potent analgesia from acute toxicity and abuse liability.

Understanding how chemically distinct opioids produce different signaling efficacies that give rise to beneficial or adverse effects is therefore central to the development of safer analgesics. However, the molecular determinants that enable such functional tuning remain incompletely understood. Upon agonist binding, MOR undergoes conformational changes that promote activation of its cognate G proteins, primarily of the Gi/o family. Activation proceeds through the transition of the inactive guanosine diphosphate (GDP)-bound Gαβγ heterotrimer to an active state initiated by receptor-catalyzed separation of the Gα α-helical domain (AHD) from the Ras homology domain (RHD), followed by GDP release from Gα, exchange with guanosine triphosphate (GTP), and a cascade of structural rearrangements that culminates in dissociation of the GTP-bound Gα and Gβγ subunits from MOR. The liberated G protein subunits subsequently engage downstream effectors to propagate intracellular signaling.

Although numerous high-resolution structures have defined the architecture of active MOR-G protein complexes^2–14^, their static nature offers limited insight into the thermodynamics and kinetics that govern ligand efficacy. Recent single particle cryogenic electron microscopy (cryo-EM) studies^10^ and time-resolved cryo-EM experiments^11^ have captured distinct intermediate conformations of the MOR-G_i1_ complex with GDP or GTP bound along the activation pathway, providing important structural and mechanistic insights into G protein activation and ligand efficacy. Together with single-molecule experiments and microsecond-scale conventional molecular dynamics (MD) simulations,^11^ these studies suggest that agonist efficacy is determined not only by the conformations of individual intermediates but also by the kinetics of the transitions between them. However, they do not provide a quantitative thermodynamic and kinetic description of the activation landscape. Such a description requires sufficient sampling of the conformational ensemble to identify intermediate and transition states, characterize the pathways connecting them, and quantify free-energy differences between conformational states, free-energy barriers, equilibrium populations, transition rates, mean first-passage times, and rate-limiting steps.

Here, we perform millisecond-scale adaptive-sampling all-atom MD simulations initiated from the GTP-bound snapshots of MOR–G_i1_ activation^11^ identified by time-resolved cryo-EM (termed GTP-primed, G-ACT-1, G-ACT-2, and G-ACT-3) and build Markov state models (MSMs) of the initial steps of the activation landscape of GTP-bound G_i1_ in complex with MOR bound to either the full agonist lofentanil (LFT) or the weak partial agonist mitragynine pseudoindoxyl (MP).

## Results

### Simulations Reveal Hidden Intermediate States Connecting GTP-Bound Conformations Along the MOR–G_i1_ Activation Pathway

To elucidate the transition pathways connecting the time-resolved cryo-EM structures of the GTP-bound LFT–MOR–G_i1_ and the MP–MOR–G_i1_ complexes resolved to date (termed GTP-primed, G-ACT-1, and G-ACT-2/3),^11^ we set up an adaptive sampling strategy that iteratively expanded the conformational exploration between the known states (see Methods for details). Briefly, explorative MSMs were used to identify disconnected or poorly sampled regions of conformational space, which were bridged using biased non-equilibrium steering simulations to generate new seed structures for subsequent rounds of unbiased sampling. This approach yielded ∼1 millisecond of aggregate unbiased simulation data (∼546 µs for LFT–MOR–G_i1_ and ∼410 µs MP–MOR–G_i1_; Supplementary Table 1).

To characterize the transitions between the aforementioned experimental states of GTP-bound LFT–MOR–G_i1_ and MP–MOR–G_i1_ complexes, production MSMs were constructed from the complete unbiased simulation datasets using a shared featurization that included selected Cα-Cα distances and fractions of native contacts (FNCs) relative to each cryo-EM structure (see Methods and Supplementary Table 2). Trajectories were projected onto a shared 7-dimensional time-lagged Independent Component Analysis (tICA) space, and clustered into microstates (see details in Methods). To regularize kinetic estimates and assess uncertainty, we constructed 100 maximum-likelihood MSMs from bootstrap samples of the ligand-bound MOR–G_i1_ trajectories and retained only models containing complete transition paths between the GTP-primed-like and G-ACT-2/3-like states. Based on implied-timescale analysis (see Methods and Supplementary Figure 1), a lag-time of 20 ns was selected for final model construction.

The free energy landscapes resulting from averaging over the bootstrap samples are shown in Figure 1. Microstates resembling the GTP-primed, G-ACT-1, G-ACT-2, and G-ACT-3 cryo-EM conformations (filled black circles in Figure 1) were identified using the FNCs and grouped into the corresponding cryo-EM-like macrostates (crosses in Figure 1). The remaining microstates were partitioned into 4 intermediate macrostates (I1-I4; see Methods), yielding a final 7-state kinetic model. The consistency of this model with the simulated dynamics was confirmed by a Chapman-Kolmogorov Markovianity test (Supplementary Figure 2).

**Figure 1.**
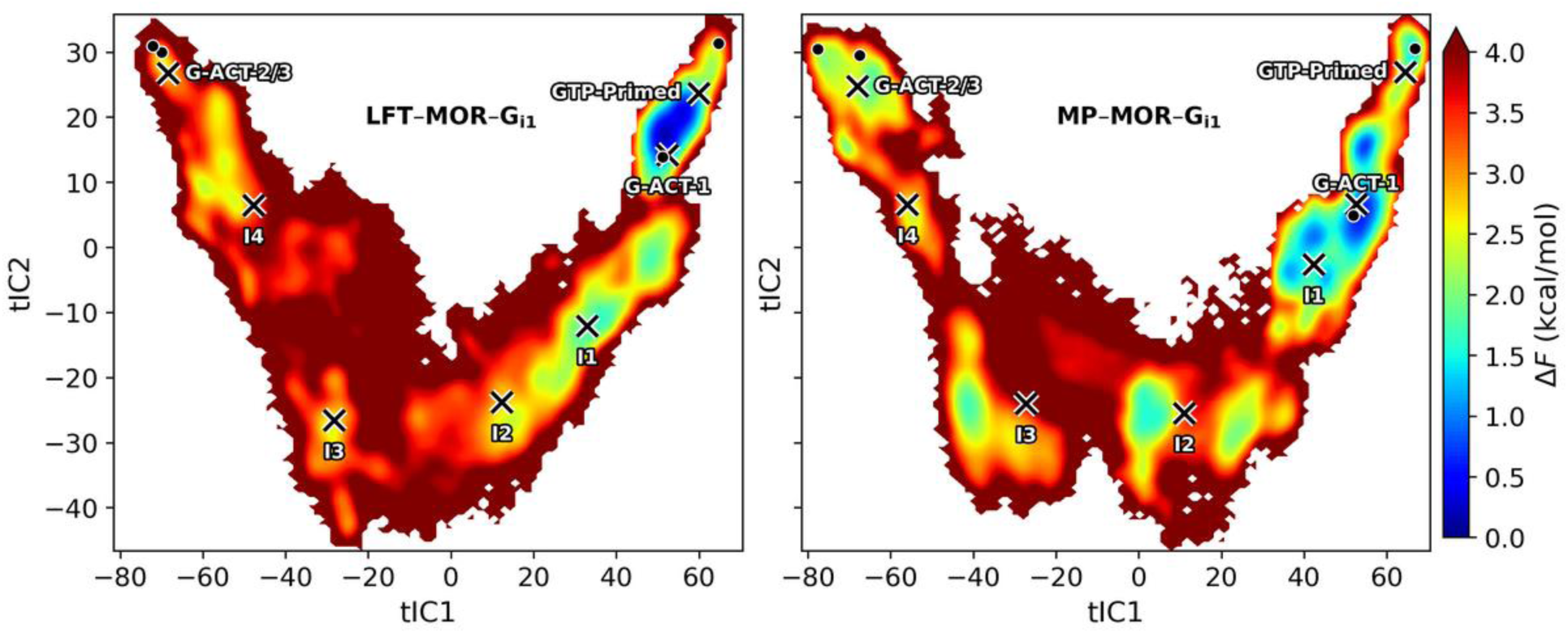
Free-energy landscapes of MOR-mediated G_i1_ activation. Two-dimensional average free-energy surfaces projected onto the two slowest reaction coordinates, tIC1 and tIC2, for the LFT-bound (left) and MP-bound (right) MOR-G_i1_ complexes. The color scale represents the average relative free energy (ΔF) in kcal/mol, calculated from the equilibrium probability distribution, with dark blue denoting the lowest free-energy (most populated) conformational basins and red indicating higher free-energy, less populated regions. Filled black circles indicate states corresponding to the cryo-EM GTP-primed, G-ACT-1, G-ACT-2, and G-ACT-3 structures. The corresponding macrostates identified by MSM analysis are indicated by labels and crosses. Four previously uncharacterized intermediate macrostates are also denoted by crosses and labeled I1-I4.

### LFT and MP Differentially Stabilize GTP-Bound MOR-G_i1_ Macrostates

Although GTP-induced MOR–G_i1_ conformational transitions are intrinsically non-equilibrium processes, the steady-state populations obtained from MSM analysis of the LFT- and MP-bound MOR–G_i1_ simulations provide estimates of the relative thermodynamic stability of macrostates along the transition pathway from the GTP-primed state to G-ACT-2/3 states in the two ligand-bound systems. These thermodynamic preferences are in agreement with the relative ligand-dependent populations of the G-ACT-1 and G-ACT-2/3 conformational states inferred from time-resolved cryo-EM.^11^ The stationary populations across bootstrap samples for all identified macrostates are shown in Figure 2, with the corresponding median values and 95% confidence intervals reported in Supplementary Table 3.

**Figure 2.**
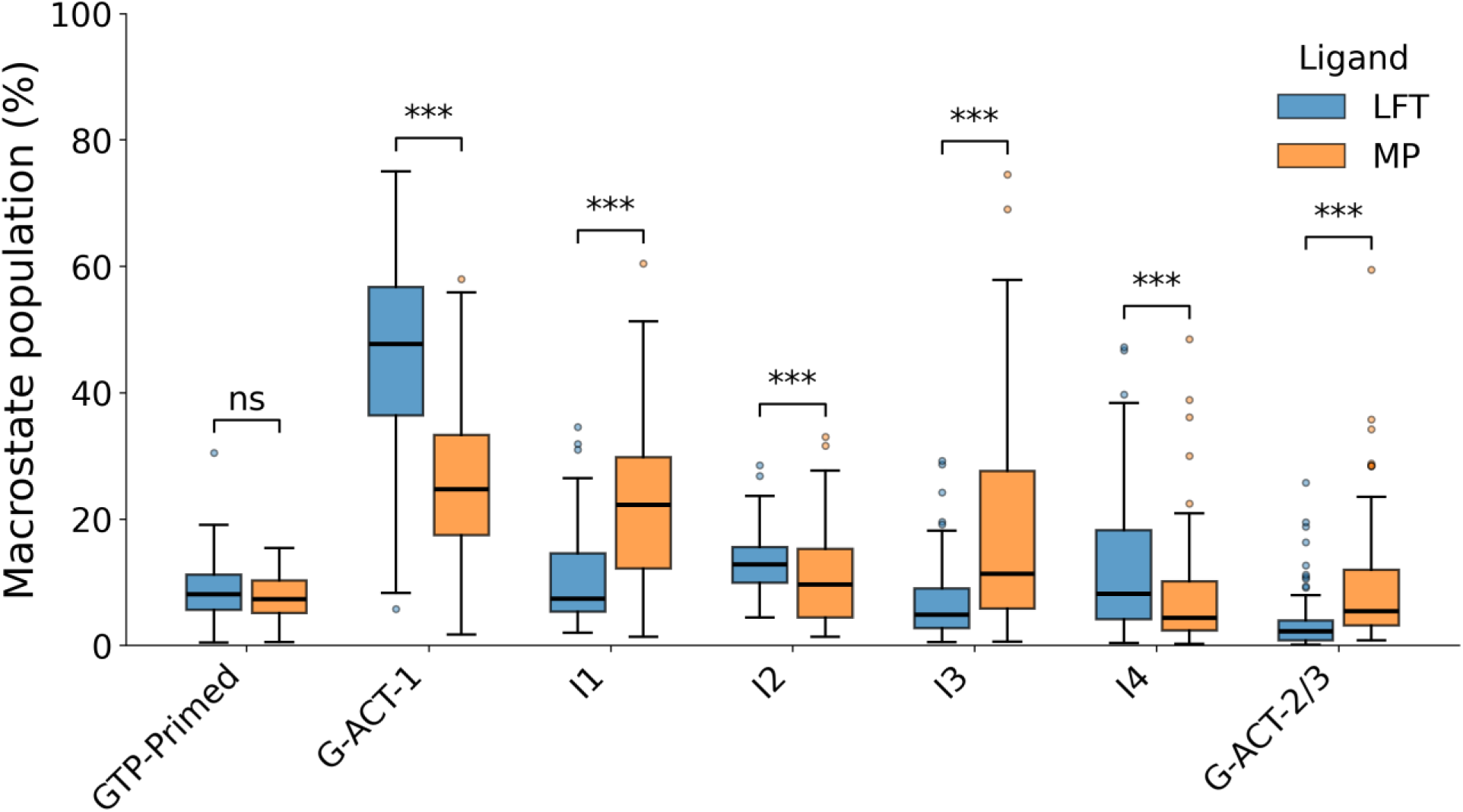
MSM-derived equilibrium populations of the metastable macrostates along the transition pathway from the GTP-primed state to the G-ACT-2/3 states. Box plots represent the steady-state populations of the seven identified macrostates for LFT-bound (blue) or MP-bound (orange) MOR–G_i1_ complexes across 93 and 82 bootstrapped models, respectively. The center line indicates the median, the box spans the interquartile range (IQR), the whiskers extend to 1.5×IQR, and outliers are shown as open circles. Macrostates corresponding to experimentally resolved conformations are labeled GTP-primed, G-ACT-1, and G-ACT-2/3, whereas additional intermediate states identified by the MSM are labeled I1–I4. Median differences between the two systems were assessed using the Mann-Whitney U test. Statistical significance is indicated as *** for *p* < 0.001; *ns* indicates no statistically significant difference.

For both ligand-bound MOR–G_i1_ systems, GTP-primed-like states are only sparsely populated (<10%), indicating that neither agonist effectively stabilizes receptor–G protein complexes with an open AHD. Similarly, G-ACT-2/3-like states exhibit low equilibrium populations, particularly for LFT–MOR–G_i1_ (∼2%), with MP–MOR–G_i1_ being slightly more populated (∼5%), in line with the ligand-dependent conformational preferences observed in cryo-EM structures.^11^

Although previous cryo-EM analyses emphasized the GTP-primed state as a prominent non-equilibrium activated conformation following GTP addition,^11^ our analysis identifies the G-ACT-1-like macrostate as the most populated state in both ligand-bound complexes. In the LFT– MOR–G_i1_ ensemble, G-ACT-1 accounts for approximately 48% of the equilibrium population. In contrast, MP shifts the conformational distribution away from this state, reducing its population to ∼25% while increasing the occupancy of intermediate states along the transition pathway from GTP-primed to G-ACT-2/3 states, most notably I1 (∼22%) and I3 (∼11%). By comparison, each of these intermediate states accounts for less than ∼13% of the population in the LFT-bound system, indicating that MP stabilizes a broader distribution of partially activated receptor–G protein conformations relative to LFT.

The identification of four distinct intermediate states (I1-I4) in the conformational landscapes of both the LFT–MOR–G_i1_ and MP–MOR–G_i1_ systems prompted us to reanalyze the previously acquired time-resolved cryo-EM datasets used to reconstruct the G-ACT1–3 states.^11^ We sought to determine whether these computationally identified intermediate states are also detectable in the experimental conformational ensemble identified by 3D variability analysis (3DVA) (see Methods). Because the G protein was resolved at substantially higher resolution than the receptor in the original cryo-EM reconstructions for the G-ACT1–2 states,^11^ we generated density maps for the Gα and Gβ subunits from the all-atom microstates assigned to states I1–I4 by MSM analysis of the LFT–MOR–G_i1_ system, which served as a reference example to illustrate the analysis (see Methods). These G protein subunits were chosen because their conformational changes are captured by the shared featurization used in our MSM. The resulting model-derived maps were independently fitted into the corresponding Gαβ densities of 3DVA-derived cluster maps generated from the LFT–MOR–G_i1_ cryo-EM data by clustering particles along a single 3DVA component. For each model-derived map, the best-matching cluster was identified by both visual inspection and cross-correlation scoring between the density maps. All four intermediate states (I1-I4) showed global agreement with their corresponding cryo-EM densities, supporting the existence of the computationally predicted intermediates (Figure 3; see also Supplementary Figure 3).

**Figure 3.**
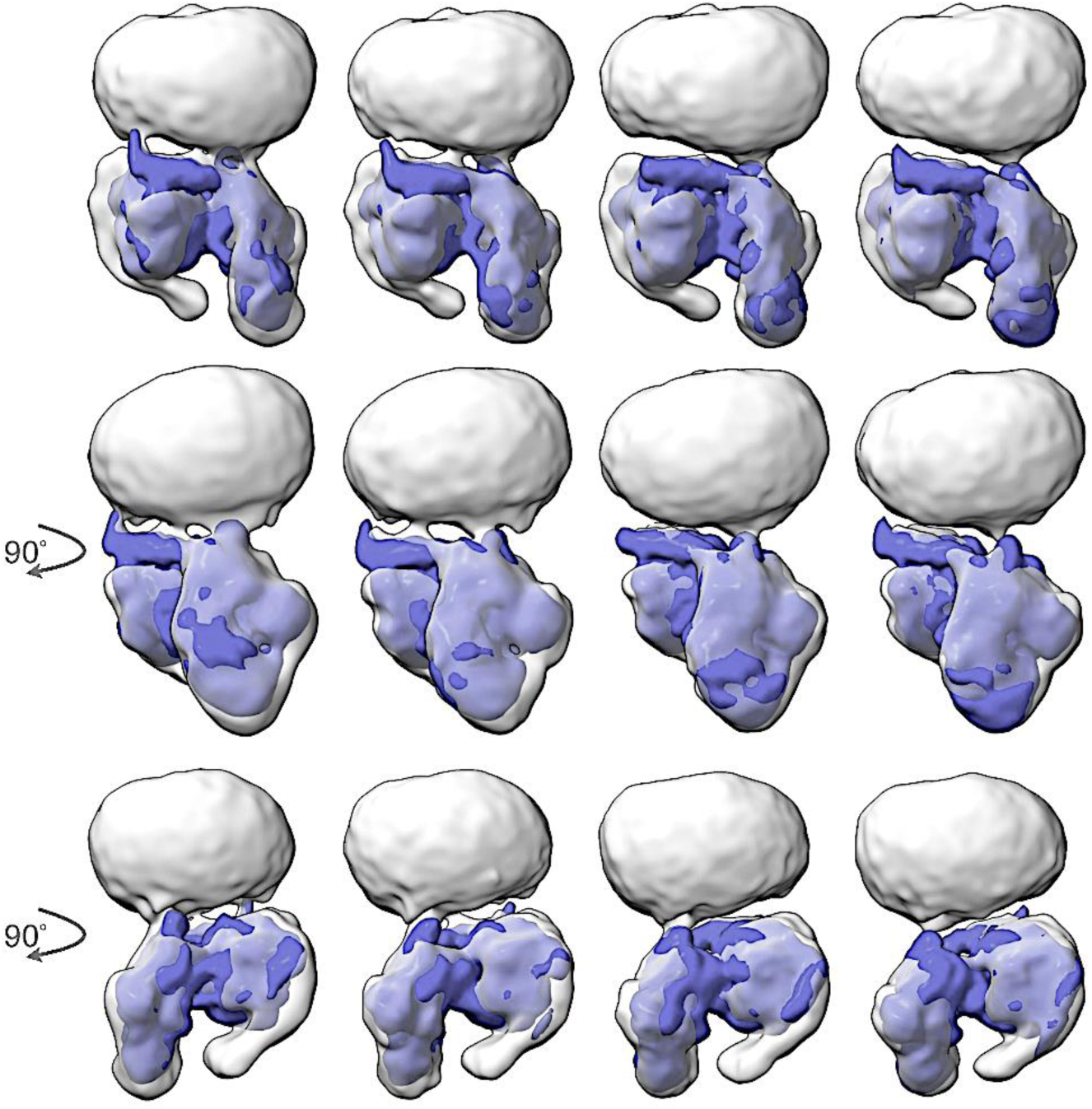
Experimental validation of MD-derived intermediate states by 3DVA. Gα-Gβ density maps (blue), generated from the all-atom microstates corresponding to the I1-I4 states identified by MSM analysis of MD simulations of the LFT-MOR-G_i1_ complex, were fitted into the corresponding low-resolution Gα-Gβ density regions of the 3DVA-derived cryo-EM cluster maps (transparent grey). The top row shows side views, while the middle and bottom rows show successive 90° rotations. The fitted models reveal a progressive conformational transition characterized by stepwise disengagement of the Gα α5 helix from the receptor, accompanied by increasing bending of the AHD as the system progresses from I1 to I4. Fitting was performed using ChimeraX v1.10 (see Methods), yielding map correlation coefficients of 0.9191, 0.9227, 0.9252 and 0.9281 for I1, I2, I3, and I4, respectively. The corresponding 3DVA cluster maps have estimated resolutions of 7.2 Å, 6.0 Å, 5.7 Å, and 5.8 Å, with occupancies of 2.87%, 6.75%, 7.67%, and 8.96% of the analyzed particle dataset, respectively (see Methods).

### Newly Identified Intermediate States Along the GTP-Bound MOR-G_i1_ Transition Pathway Display Ligand-Specific Structural Features

Inspection of the mean maximum FNC for each macrostate relative to the available cryo-EM structures (see Methods and Supplementary Figure 4) reveals a clear separation between the intermediate and cryo-EM-like states. The intermediate states I1-I4 exhibit substantially lower similarity to the published cryo-EM structures (mean maximum FNC<45%) than the GTP-primed, G-ACT-1, and G-ACT-2/3 states, all of which are cryo-EM-like and exhibit mean maximum FNC values ≥ 60%.

To further characterize these newly identified intermediates (see representative structures in Supplementary Figure 5), we quantified a series of MOR and G protein structural descriptors previously shown to distinguish the experimentally characterized GTP-primed, G-ACT-1, and G-ACT-2/3 states^11^ (Supplementary Figure 6; see Methods). These descriptors included the Gα AHD opening angle, the number of MOR-Gα contacts, the helicity of the Gα αN and α5 helices, and the helicity of the receptor intracellular loops 2 and 3 (ICL2 and ICL3). We also monitored the R182-R273 distance between transmembrane (TM) helix 4 (TM4) and TM6 (R182^4×40^–R273^6×28^ in generic numbering; see Methods), together with a set of MOR–G protein contacts involving receptor residues in ICL3, TM5, and TM6 and either Gα or Gβ subunits (see Supplementary Figure 6 caption for details).

Remarkably, the largest structural differences between the MP- and LFT-bound MOR–G_i1_ systems are not observed in the experimentally resolved GTP-primed, G-ACT-1, or G-ACT-2/3 states, but instead in the newly identified intermediates I1–I3. These differences primarily involve Gα AHD closure and MOR–Gα interactions, although large ligand-dependent changes are also evident in the helicity and interactions of MOR ICL2 and ICL3, as well as the helicity of Gα αN and α5 helices, and interactions between the Gα α5 helix and MOR (Supplementary Figures 5 and 6). For example, although progression from the GTP-primed state to G-ACT-1 is characterized by AHD closure with AHD fluctuations becoming increasingly stabilized through the I4 state in both systems, the AHD remains consistently more open in the MP-bound than the LFT-bound MOR-G_i1_ system throughout I1-I3. Likewise, Gα disengagement begins in I2, where the α5 helix loses approximately 20% of its helicity. This loss is more pronounced in the MP-bound MOR-G_i1_ system, whereas the largest reduction in receptor–G protein contacts occurs in the LFT-MOR-G_i1_ complex. Consequently, the G protein remains associated with the receptor for longer in the MP-bound system, only undergoing substantial disengagement upon reaching I3.

Receptor descriptors that distinguish active-like (GTP-primed and G-ACT-1) from inactive-like (G-ACT-2/3) conformations further indicate that I3 represents the first receptor intermediate adopting an inactive-like architecture. This transition, monitored by the R182^4×40^– R273^6×28^ distance, occurs more abruptly in the LFT-bound than in the MP-bound MOR-G_i1_ system.

### LFT and MP Elicit Distinct Most Probable Transition Pathways and Kinetics

We used transition path theory (TPT)^15,16^ to identify the most probable transition pathways and their reactive flux connecting the GTP-primed and G-ACT-2/3 states in the LFT–MOR–G_i1_ and MP–MOR–G_i1_ complexes (Figure 4 and Supplementary Table 4). We also calculated dwell times (Supplementary Figure 7 and Supplementary Table 5) and mean first-passage times (MFPTs) between pairs of macrostates from the MSMs (Figure 5, Supplementary Figures 8 and 9, and Supplementary Table 6).

**Figure 4.**
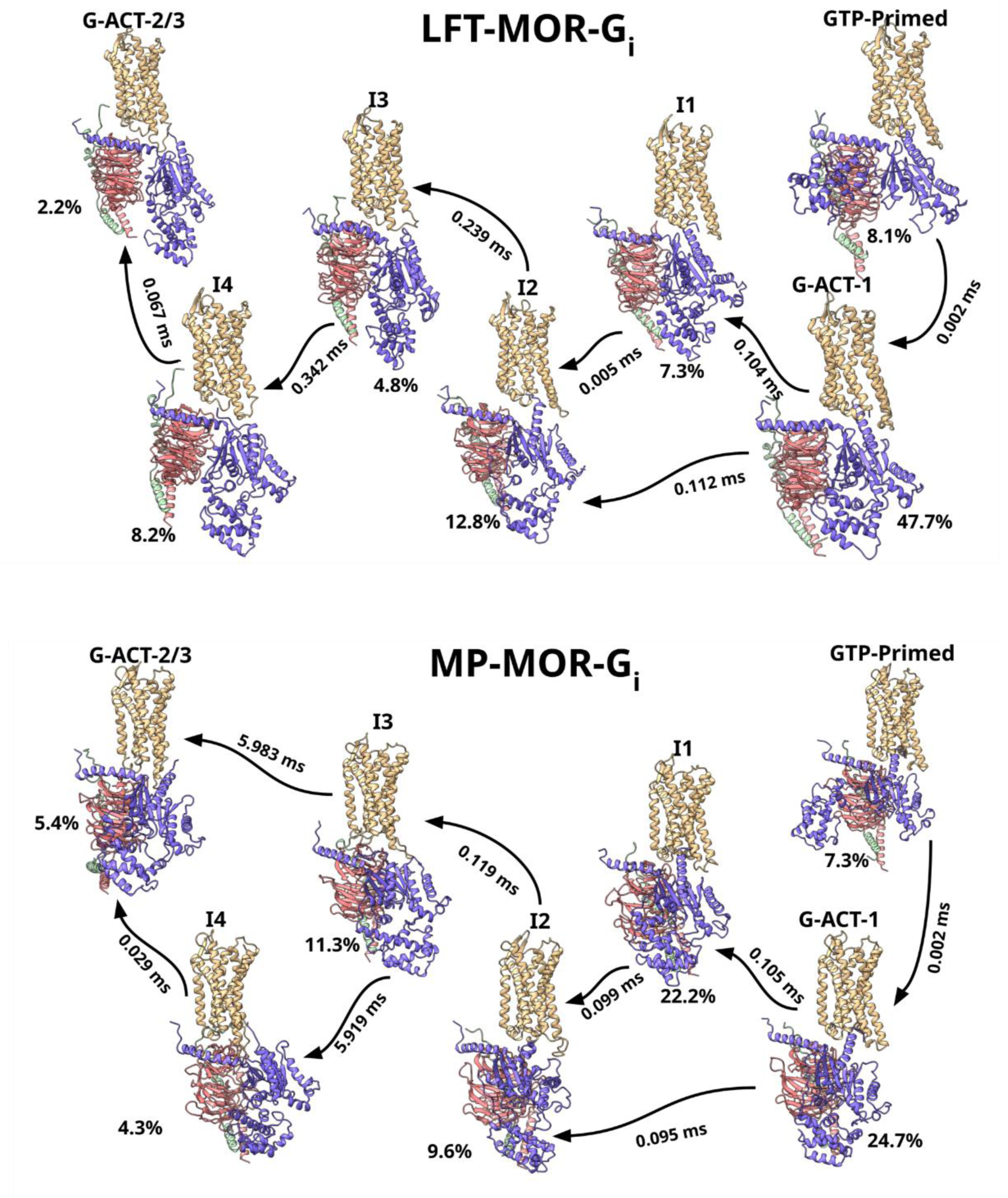
Ligand-Specific Kinetic Networks of MOR-G_i1_ Transitions from GTP-primed to G-ACT-2/3 States. Kinetic network representations of transition pathways obtained from filtered bootstrapped Markov state models (see Methods) for the LFT-MOR-G_i1_ (top) and MP-MOR-G_i1_ (bottom) complexes. Structures are macrostate representatives labeled by state name and positioned approximately according to their locations in tICA space (see Figure 1); macrostate populations are reported as median percentages (see Supplementary Table 3). MOR is shown in tan, Gα in blue, Gβ in red, and Gγ in green. Black arrows indicate the most probable transition pathways connecting the GTP-primed and G-ACT-2/3 states, as identified by transition path theory (see Supplementary Table 4), and labeled by their MFPTs (see Supplementary Table 6).

**Figure 5.**
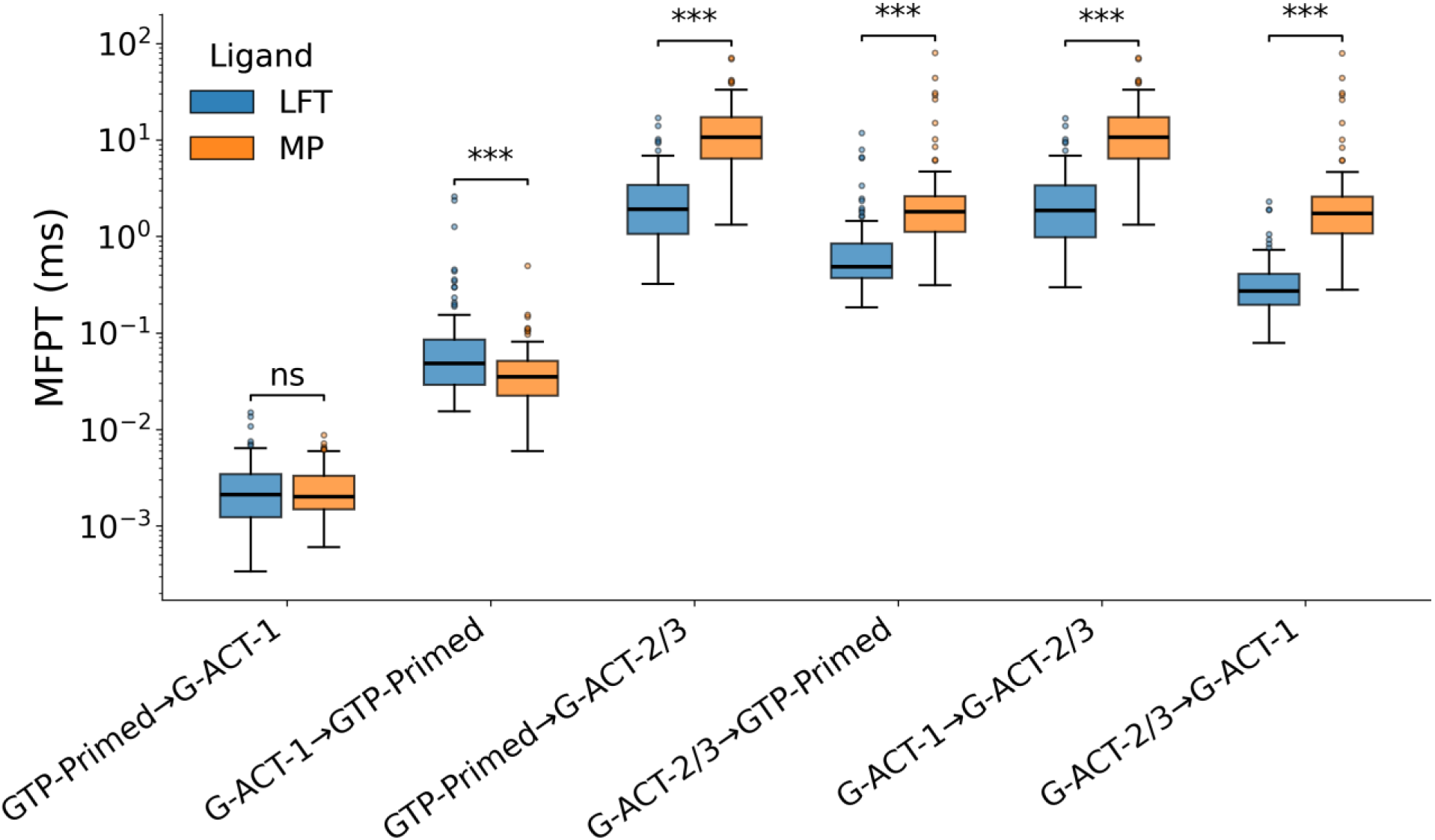
Mean first-passage times between experimental-like conformations. Box plots compare the distributions of bidirectional mean first-passage times (MFPTs; ms) for transitions between the cryo-EM-like GTP-primed, G-ACT-1, and G-ACT-2/3macrostates in LFT-MOR-G_i1_ and MP-MOR-G_i1_. MFPT distributions were obtained by bootstrap sampling. The center line indicates the median, the box spans the interquartile range (IQR), the whiskers extend to 1.5×IQR, and outliers are shown as open circles. Median value difference statistical significance was assessed using the Mann-Whitney U test, with *** denoting p < 0.001. *ns* denotes no statistically significant difference.

As summarized in Supplementary Table 4 and illustrated in Figure 4, the two systems exhibit distinct highest-flux transition pathways from the GTP-primed state to the G-ACT-2/3 state.

In the LFT–MOR–G_i1_ system, the GTP-primed → G-ACT-1 → I1 → I2 → I3 → I4 → G-ACT-2/3 pathway dominates the transition network, accounting for ∼99% of the total coarse-grained flux across the corresponding ligand-specific models (total reactive flux over all MSM bootstrapped models). In contrast, this pathway contributes only ∼13% of the total reactive flux over all MSM bootstrapped models in MP–MOR–G_i1_, which instead exhibits a more heterogeneous transition network. The dominant transition pathway in MP–MOR–G_i1_ from the GTP-primed and G-ACT-2/3 states bypasses I4, transitioning directly from I3 to G-ACT-2/3 and accounting for an average of ∼68% of the total reactive flux over all models. Two additional pathways unique to MP–MOR–G_i1_ bypass I1 or both I1 and I4, contributing ∼11% and ∼9% of the total reactive flux over all MSM bootstrapped models, respectively. The pathway bypassing I1 is also observed in LFT–MOR–G_i1_ but contributes only ∼1% of the total model flux. Despite sharing similarities in the pathway topology, the two systems differ substantially in their transition kinetics. Along the common GTP-primed → G-ACT-1 → I1 → I2 → I3 → I4 → G-ACT-2/3 pathway (see Supplementary Table 4), the MFPT is markedly longer for MP–MOR–G_i1_ (54.89 ms; 95% confidence interval (CI): 14.27–411.49 ms) than for LFT–MOR–G_i1_ (2.61 ms; 95% CI: 0.97–12.95 ms). The fastest MP–MOR–G_i1_ pathway, which bypasses I4, also exhibits a substantially longer MFPT (18.22 ms; 95% CI: 4.86–69.79 ms) than the dominant LFT–MOR–G_i1_ pathway. Notably, in LFT–MOR–Gi1, the low-flux pathway that bypasses I1 dramatically increases the MFPT from 2.61 ms (95% CI: 0.97–12.95 ms) to 144.04 ms (95% CI: 19.23–886.19 ms), highlighting the critical role of I1 in promoting efficient GTP-primed-G-ACT-2/3 transition.

As shown in Figure 5 (see Supplementary Table 6 for numerical values), the initial transition from the GTP-primed state to G-ACT-1 is rapid for both ligands (∼2 µs), while reverse transitions are ∼20-fold slower for both ligands (49 µs vs. 2.1 µs for LFT-MOR-G_i1_ and 35 µs vs. 2.0 µs for MP-MOR-G_i1_), confirming strong thermodynamic stabilization of the G-ACT-1 state in both systems.

Beyond this initial transition, however, the kinetics diverge markedly for the two different ligand-bound systems. The total mean first passage times from the GTP-primed or G-ACT-1 states to G-ACT-2/3 (Supplementary Table 6) are substantially slower in MP-MOR-G_i1_ (MFPT ≈ 11 ms) than in LFT-MOR-G_i1_ (MFPT ≈ 1.9 ms), while the reverse transitions occur 4-6-fold faster (∼490 μs vs. 1.9 ms for LFT-MOR–G_i1_ and ∼1.8 ms vs. 11 ms for MP-MOR-G_i1_), consistent with the lower stability of the G-ACT-2/3 states. These kinetic differences are consistent with the broader distribution of intermediate states between GTP-primed/G-ACT-1 and G-ACT2/3 in MP-MOR-G_i1_ relative to LFT-MOR-G_i1_. In particular, the pronounced stabilization of the I1 and I3 states in the MP-bound MOR-G_i1_ free energy landscape (Figure 1) appears to hinder progression toward the G-ACT-2/3 states. Consistent with this interpretation, the I1 → I4, I2 → I4, and I3 → I4 transitions are substantially slower for the MP-bound MOR-G_i1_ than the LFT-bound MOR-G_i1_ system (11 ms versus 1.3 ms, 11 ms versus 1.4 ms, and 5.9 versus 0.34 ms, respectively; see Supplementary Figure 8 and Supplementary Table 6). As noted in the previous section, structurally, this slowdown is associated with a more compact and conformationally restricted intermediate ensemble in the MP-bound MOR-G_i1_ system, characterized by a greater number of MOR-G_i1_ contacts, including tighter coupling between the receptor intracellular core and the G protein α5 helix, as well as a more open AHD conformation (Supplementary Figure 6). Together, these interactions appear to stabilize intermediate states and hinder the large-scale conformational rearrangements required to reach the G-ACT-2/3 states, and their possible progression towards G protein dissociation and downstream signaling.

Dwell-time analysis (Supplementary Figure 7 and Supplementary Table 5) reveals that the intermediate macrostates I1-I4 have residence times comparable to those of cryo-EM-like states in the two ligand-bound MOR-G_i1_ systems. Nevertheless, marked differences in residence times were observed between the LFT- and MP-bound MOR-G_i1_ systems, with the largest differences occurring for G-ACT-1 (3.91 µs [3.50, 4.56] vs. 1.14 µs [0.98, 1.30]) and I4 (3.28 µs [2.93, 3.59] vs. 0.89 µs [0.79, 0.99]).

## Discussion

A major finding of this work is that the transition between the previously identified GTP-primed and G-ACT-2/3 conformations along the MOR-mediated G protein activation pathway is considerably more complex than suggested by the experimentally resolved structures alone. Rather than proceeding solely through the previously described G-ACT-1 state, both the full agonist (LFT)- and partial agonist (MP)-bound MOR–Gi1 complexes traverse a series of previously uncharacterized metastable intermediates with distinct structural and kinetic properties. The identification of these intermediates, together with their correlation to populations in the experimental time-resolved cryo-EM data, suggests that they are not artifacts of molecular dynamics simulations, but bona fide conformational states populated during G protein activation.

By integrating time-resolved cryo-EM with Markov state modeling of millisecond-scale MD simulations, our study provides a substantially more complete description of the thermodynamic and kinetic landscape of MOR-mediated G protein activation than either approach alone.

Our analysis further supports the hypothesis that ligand efficacy is encoded by differences in the stability and interconversion kinetics of intermediate conformations. The largest structural differences between the full agonist LFT and the partial agonist MP occur within the newly identified intermediate states rather than in the GTP-primed, G-ACT-1, or G-ACT-2/3 conformations themselves. This observation is consistent with an emerging view of GPCR activation in which efficacy is determined by redistribution of conformational ensembles and modulation of transition kinetics rather than stabilization of a single active state. In the present system, the partial agonist broadens the free-energy landscape, stabilizes multiple intermediate conformations, and substantially slows progression toward the later G-ACT-2/3 states, whereas the full agonist favors a more directed and kinetically efficient transition pathway.

Transition path analysis also provides mechanistic insight into the sequence of structural events leading toward G protein dissociation. The temporal separation of conformational changes across the kinetic network supports a stepwise, rather than concerted, mechanism of G protein disengagement. Specifically, displacement of the Gα α5 helix from the receptor’s intracellular cavity precedes receptor deactivation: MOR retains an active-like conformation despite substantial weakening of receptor–G protein interactions and transitions toward an inactive-like conformation only after extensive α5 disengagement. Consistent with this sequence, the direct I2→I3 transition constitutes an obligatory step along the dominant pathways connecting the GTP-primed and G-ACT-2/3 states for both agonists.

Although the present simulations did not capture complete dissociation of G_i1_ from MOR, the relative populations and kinetics of the intermediate states reveal clear agonist-dependent differences in the progression toward this endpoint. In the LFT-MOR-G_i1_ system, the I2 state is more highly populated than in MP-MOR-G_i1_, suggesting that the full agonist preferentially stabilizes an active-like receptor conformation even after receptor–G protein interactions have been substantially weakened. In contrast, the higher population of I3 in MP-MOR-G_i1_ indicates that the partial agonist preferentially stabilizes an inactive-like intermediate that is incompatible with productive G protein binding. Consistent with this interpretation, kinetic analysis shows that the I2→I3 transition occurs more rapidly in MP-MOR-G_i1_, whereas the reverse I3→I2 transition is faster in LFT-MOR-G_i1_. Together, these observations indicate that the principal energetic distinction between the two agonists lies not in the initiation of G protein activation characterized by AHD closure upon GTP binding, which proceeds rapidly for both ligands, but rather in the stability of intermediate states that determine whether the receptor remains competent to support G protein engagement or progresses toward receptor deactivation.

The ligand-dependent transition pathways identified here further suggest that efficacy is governed by the organization of the kinetic network itself. Lofentanil predominantly follows a single dominant pathway that sequentially traverses all intermediate states before reaching G-ACT-2/3, whereas mitragynine pseudoindoxyl distributes reactive flux across several alternative pathways and frequently bypasses I4. This increased pathway heterogeneity, together with the slower transition rates between intermediates, is consistent with a more rugged free-energy landscape in the partial agonist-bound complex. Such kinetic heterogeneity may divert part of the signaling flux toward less productive pathways or promote trapping in intermediate states, thereby contributing to the reduced signaling efficacy characteristic of partial agonists.

More broadly, these findings reinforce the concept that GPCR efficacy cannot be understood solely from static structural comparisons. Instead, efficacy emerges from the interplay between thermodynamics and kinetics across an ensemble of transient conformational states. By explicitly quantifying state populations, transition pathways, and timescales, the present work extends the structural information provided by time-resolved cryo-EM into a quantitative mechanistic model of MOR-mediated G protein activation. Because all GPCRs are expected to undergo nucleotide-dependent transitions during G protein activation, the computational framework presented here should be broadly applicable to dissecting efficacy mechanisms throughout the GPCR superfamily.

A few important limitations should nevertheless be acknowledged. First, although the aggregate unbiased simulation time approached 1 millisecond, complete dissociation of G_i1_ from MOR was not observed, suggesting that this process either occurs on longer timescales than those accessed here or requires enhanced sampling along collective variables different from those employed in the present study. Consequently, the present MSMs describe the early stages of G protein activation following GTP binding rather than the complete signaling cycle. Second, the inferred thermodynamic quantities should be interpreted within the context of the modeled state space. GTP-driven G-protein activation is intrinsically a nonequilibrium process, whereas the stationary distributions of the reversible MSMs describe equilibrium-like populations of the discrete conformational states represented in the models. Third, characterization of the intermediate states and their kinetics depends on the choice of model parameters, including structural featurization, dimensionality reduction, clustering, lag time, and macrostate definitions used to construct the MSMs. Nevertheless, the agreement between the predicted intermediate states and independent cryo-EM conformational ensembles, together with the internal consistency of the MSM-derived thermodynamic and kinetic observables, supports the biological relevance of the identified intermediates and transition pathways. Finally, the simulations used mouse MOR in complex with human G_i1_, a specific membrane composition, and a particular set of force-field and protonation-state assumptions. Although these conditions were selected to reproduce the available structural systems, differences in receptor species, G-protein subtype, membrane composition, or molecular interactions not represented in the simulations could alter the relative populations and kinetics of the identified states.

In summary, this work provides a quantitative mechanistic framework linking ligand efficacy to the kinetics of early G protein activation by MOR. Rather than altering a single activation step, full and partial agonists reshape the free-energy landscape by differentially stabilizing intermediate conformations and modulating the rates of transitions between them. These results suggest that transient kinetic intermediates, rather than only the experimentally resolved endpoint conformations, represent promising targets for the rational design of opioid ligands capable of selectively tuning signaling efficacy while potentially minimizing adverse effects.

## Methods

### Generic Residue Numbering for Mouse MOR and Human G_i1_

To facilitate broad interpretation and generalization of our results, we use the structure-based generic GPCRdb numbering scheme^17^ for OPRM1_Mouse and the generic numbering scheme for Gαi^18^ (GNAI1_Human) throughout this study.

### System Setup

Eight all-atom systems of LFT- or MP-bound MOR-Gα_i1_β_1_γ_2_ complexes containing GTP and Mg^2+^ in the Gα_i1_ nucleotide-binding site were prepared for MD simulations. Initial coordinates were based on previously refined cryo-EM structures^11^ representing four conformational states of ligand-bound mouse MOR in complex with human G_i1_: GTP-primed (deposited as PDB IDs 9ODF and 9ODJ for LFT- and MP-bound MOR-G_i1_ systems, respectively); G-ACT-1 (deposited as PDB ID 9ODG for the LFT-bound MOR-G_i1_ system, with the corresponding MP-bound MOR-G_i1_ model generated by ligand replacement); G-ACT-2 (for the LFT-bound MOR–G_i1_ complex, only the G_i1_ subunit was deposited as PDB ID 9ODH, whereas the complete MP-bound MOR–Gi1 complex was deposited as PDB ID 9ODK); and G-ACT-3 (deposited as PDB ID 9ODL for the MP-bound MOR–G_i1_ system, with the corresponding LFT-bound MOR–G_i1_ model generated by ligand replacement). All mutations introduced for structure determination were reverted to the corresponding wild-type sequences of mouse MOR and human G_i1_. Missing side chains and short unresolved segments were modeled by homology or *ab initio*. These included: MOR helix 8 (residues C356^8×53^-I352^8×59^), the Gα_i1_ N terminus (residues G2^G.HN.02^ - T4^G.HN.10^), and Gγ segments A7-S8 and K64-C68. Homology modeling was guided by high-resolution structures of MOR (PDB ID 4DKL^19^) and Gα_i1_ (PDB ID 1GP2^20^), while *ab initio* building was performed with MODELLER 10.5^21^.

Systems were parameterized using the CHARMM General Force Field^22^ for ligands and the CHARMM36m force field^23^ for proteins and lipids. Membrane systems were constructed with the CHARMM-GUI Bilayer Builder^24,25^. MOR residue D114^2×50^ was modeled in its protonated form, while all other titratable residues were assigned protonation states corresponding to pH 7.0.

To reproduce the physiological membrane anchoring of the Gα_i1_β_1_γ_2_ heterotrimer, Gα_i1_ was modeled with N-terminal myristoylation at G2^G.HN.02^ and palmitoylation at C3^G.HN.03^, while Gγ was modeled with geranylgeranylation at C68, consistent with prior studies^26^. Each complex was embedded in a lipid bilayer approximating mammalian plasma membrane composition, comprising 1-palmitoyl-2-oleoyl-sn-glycero-3-phosphocholine (POPC), 1-palmitoyl-2-oleoyl-sn-glycero-3-phosphoethanolamine (POPE), 1-palmitoyl-2-oleoyl-sn-glycero-3-phosphoserine (POPS), palmitoyl sphingomyelin (PSM), monosialodihexosylganglioside (GM3), cholesterol, and 1,2-diacyl-sn-glycero-3-phospho-1-D-myo-inositol 4,5-bisphosphate (PIP2) at concentrations previously reported in the literature^27^. Systems were solvated with TIP3P water^28^ and neutralized with 0.15 M NaCl. Hydrogen mass repartitioning^29^ was applied by increasing hydrogen masses to 4 atomic mass units and correspondingly reducing the mass of bonded heavy atoms, enabling a 4-fs integration timestep. System compositions are summarized in Supplementary Table 7.

### Molecular Dynamics Simulations

All systems were equilibrated in GROMACS 2025.1^30^ using a stepwise restraint-release protocol. Following 5,000 steps of steepest-descent energy minimization, restrained NPT equilibration was performed as detailed in Supplementary Table 8. Temperature was maintained at 303.15 K using the stochastic velocity-rescaling thermostat^31^, and pressure was maintained at 1 bar with a semi-isotropic C-rescale barostat^32^. Short-range electrostatic and van der Waals interactions were computed with a 12 Å cutoff, while long-range electrostatic interactions were computed via Particle Mesh Ewald (PME) summation^33^. Positional restraints on non-solvent heavy atoms were gradually released over the first 2 ns (see Supplementary Table 8 for details). To further equilibrate the lipid environment, each system was subsequently simulated for 200 ns with weak restraints applied to protein backbone and ligand heavy atoms. Membrane relaxation was assessed by monitoring lipid headgroup radial distributions in the membrane plane (Supplementary Figure 10).

Production simulations followed an adaptive sampling strategy (see details below) and consisted of multiple 0.5-2 μs runs performed in GROMACS using in-house computational resources^34^, as well as several longer unbiased simulations of 5-10 μs each carried out on Anton 3.^35^ For the Anton simulations, equilibrated systems were converted to Desmond format using InterMol^36^, and simulation parameters, including non-bonded cutoffs, temperature, and pressure were matched to those used in GROMACS. Temperature and pressure in Anton simulations were controlled using the Antithetic thermostat and semi-isotropic Monte Carlo barostat within the Multigrator integration framework^37^. Electrostatic interactions in these simulations were computed using the u-series/SinhGS methods^38,39^, and trajectories were propagated using an r-RESPA^40^ multiple time-step integration scheme with a 2.5 fs outer timestep, while long-range non-bonded interactions were evaluated every three steps. Coordinates were saved every 100 ps for subsequent analysis in both GROMACS and Anton simulations. In total, ∼546 µs and ∼410 µs of production simulation time were accumulated for the LFT- and MP-bound MOR-G_i1_ complexes, respectively (see Supplementary Table 1 for details).

### Sampling Strategy

Four cycles of an adaptive sampling strategy were employed to enhance conformational space exploration and accelerate transitions between cryo-EM states. This approach involved iterative rounds of multiple unbiased MD simulations, exploratory MSM construction to identify undersampled regions of the conformational space, targeted non-equilibrium simulations between metastable states, and selection of new conformations for subsequent rounds of unbiased simulations to improve sampling.

Initial unbiased simulations were initiated as multiple independent replicas from each of the eight equilibrated cryo-EM structures, with randomized initial velocities and trajectory lengths ranging from 0.5 to 10 µs. Following an aggregate of 350 µs of sampling across both ligand-bound MOR-G_i1_ systems, an exploratory MSM constructed from intra-(limited to MOR and Gα) and intermolecular (MOR-Gα, MOR-Gβγ and Gα-Gβ) Cα–Cα distances corresponding to contacts that varied across the starting structures revealed no spontaneous transitions between the G-ACT-1 and G-ACT-2/3 states, unlike between GTP-primed and G-ACT-1. After an initial screening of residue pairs with minimum heavy-atom distance below 4.5 Å in at least one of the experimental structures, pairs were retained if their Cα-Cα distances differed by at least 0.4 Å between any two structures. After pruning redundant contacts by removing pairs in which both residues formed more than three contacts with other residues while preserving network connectivity, the final feature set comprised 685 Cα-Cα distances spanning key structural elements of Gα and MOR (Supplementary Figure 11 and Supplementary Table 9).

Features were computed with MDAnalysis 2.9.0^41^, projected onto the six slowest time-lagged independent component analysis (tICA)^42,43^ components using commute-map scaling^44^, and discretized into 50 microstates by k-means clustering with k-means++ initialization^45^. tICA and clustering models were fit using the combined LFT-bound and MP-bound MOR-G_i1_ simulation datasets, while all explorative MSMs^46,47^ were built using Deeptime 0.4.5^48^ maximum-likelihood estimator, separately for each system, with transition counts obtained via a sliding window at a 30 ns lag time and enforcing detailed balance.

Under reversibility constraints, exploratory models identified the GTP-primed, G-ACT-1, and G-ACT-2/3 states as being kinetically disconnected in higher-dimensional components during sampling. To obtain a fully connected MSM, representative conformations from each pair of kinetically disconnected conformational ensembles were selected based on the minimum Euclidean distance in the first six tICA dimensions^42,43^. These conformations were then used to initiate constant-velocity non-equilibrium steering simulations in PLUMED 2.9^49^, employing moving harmonic restraints along the six tICA coordinates with a force constant of 5 kJ/(mol·[CV]^2^) per component. The restraint velocities spanned 0.6–48.6 CV/ns (5th–95th percentile range) across all pulling simulations, reflecting differences in the tICA-space distance between the source and target conformations. Biased trajectories were excluded from MSM construction and used exclusively to generate seed conformations for subsequent unbiased simulations after filtering out structures displaying local unfolding or other structural artifacts. For simulations initiated from steering-derived conformations, velocities were reassigned at 200 K and linearly increased to 303.15 K over 1 ns; this annealing phase was also excluded from subsequent MSM construction. A summary of the total number of trajectories and simulation times across all adaptive sampling rounds is provided in Supplementary Table 1.

### Production Markov State Models for Activation Kinetics

Production MSMs describing ligand-bound MOR-G_i1_ activation kinetics were constructed from the final unbiased simulation datasets of ∼545 µs and ∼410 µs collected for LFT-MOR-G_i1_ and MP-MOR-G_i1_, respectively (see Supplementary Table 1). A shared featurization was generated using MDTraj 1.11.1^50^ to capture the largest conformational changes associated with the transition between the cryo-EM-resolved GTP-primed state, G-ACT-1, G-ACT-2, and G-ACT-3 states. Specifically, the feature set comprised Cα-Cα distances corresponding to inter- and intra-chain heavy-atom contacts between MOR, Gα, and Gβ subunits and within each protein that differed by >5 Å across any pair of the cryo-EM structures (GTP-primed, G-ACT-1, G-ACT-2, G-ACT-3). The 4-dimensional vector of FNCs relative to each experimental structure^51^ was also included in the features.

All subsequent MSM analyses were performed on trajectories with 1 ns stride between consecutive frames using Deeptime 0.4.5^48^. A system-shared tICA decomposition with commute map scaling^52^ was applied to the featurized trajectories using a lag time of 5 ns after discarding the first 10 ns of each trajectory for equilibration. Dimensionality continuity analysis was then performed by constructing clustering models and maximum-likelihood MSMs^53^ across increasing tICA dimensionalities to determine the highest dimensionality yielding continuous coverage of tIC1, which led to the retention of 7 tICA dimensions for final model construction.

In the resulting 7-dimensional tICA space, system-shared minibatch-kmeans clustering^54^ was optimized using VAMP-2 scoring^55^. Although higher cluster counts improved VAMP-2 scores, finer-grained models exhibiting greater kinetic variance confined the MSM to incomplete conformational sub-spaces. MSMs spanning the full tICA space could only be constructed with up to 50 cluster centers, which were therefore retained. An additional 4 microstates were introduced for frames exhibiting FNC≥0.8 to the cryo-EM structures, yielding 54 microstates in total.

Final MSM construction employed trajectory bootstrapping^56^, in which trajectories were randomly resampled with replacement to generate 100 independent models for each ligand-specific dataset. For each bootstrap sample, reversible maximum-likelihood MSMs were built using sliding-window transition counting at lag times 5, 10, 15, 20, and 30 ns. Transition path theory (TPT)^16^ filtering was then applied to retain only models containing complete paths between GTP-primed-like and G-ACT-2/3-like microstates. Implied timescales analysis indicated that the slowest dynamical process was, on average, stable up to a lag time of 20 ns^57,58^ (Supplementary Figure 1). This lag time was therefore selected for the final analysis, as it falls within the locally converged regime while retaining sufficient transition statistics. At this lag time, 93 models for LFT-MOR-G_i1_ and 82 for MP-MOR-G_i1_ were retained. In contrast, increasing the lag time to 30 ns substantially reduced the number of MP-MOR-G_i1_ models that remained connected following transition-path filtering, indicating that estimates at longer lag times were increasingly affected by limited sampling.

Microstates with mean maximum FNC>0.5 relative to a cryo-EM reference structure were assigned to the corresponding cryo-EM-like macrostate. Intermediate states were partitioned into 4 macrostates, labeled I1, I2, I3, and I4, using hierarchical Ward clustering^59^ of FNCs. Markovianity of the discrete dynamics on the resulting 7-state decomposition was validated using a Chapman-Kolmogorov test^47^ (Supplementary Figure 2). Mean first passage times (MFPTs) from macrostates (A) to (B) were calculated using Deeptime, as the self-consistent expectation value (E) of the time to reach a state in (B) from a state in (A):

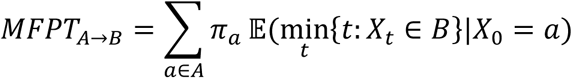

where (π_a_) are the equilibrium populations of state (A). Dwell times were calculated from the transition matrix as:

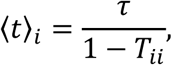

where (*τ*) is the MSM lag time and (*T_ii_*) is the self-transition probability of macrostate (i).

For each transition MFPT and dwell time, ligand-dependent differences were assessed using a two-sided Mann–Whitney U test.^60^ Statistical significance was reported as * (p < 0.05), ** (p < 0.01), and *** (p < 0.001); ns denotes no statistically significant difference.

Reactive flux analysis, as implemented in the Deeptime package, was applied to each TPT-filtered MSM. Reactive fluxes were coarse-grained over the defined macrostates (*f_AB_* = Σ*_a_*_∈*A*,*b*∈*B*_*f_ab_*) to identify the pathways with the highest net flux from the GTP-primed to G-ACT-2/3 states. Specifically, the forward committor:

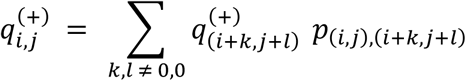

describes the probability that a path initiated in state (i,j) will progress through neighboring states (i+k,j+l) with transition probability *p*_(*i*+*k*,*j*+*l*)_. The forward committor can be obtained explicitly for transitions between target states (A and B) by solving the boundary value problem;

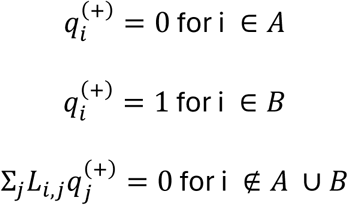

where *L* = *P* − *I* is the generator matrix and P is the transition probability matrix. From the committor, the gross reactive flux between two states can be calculated as:

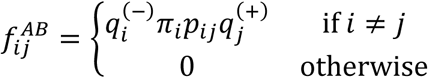

where 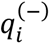 is the backward committor and *π*_i_ is the equilibrium probability of state (i). The corresponding net reactive flux can be calculated as:

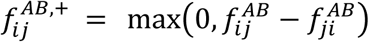

Dominant ligand-dependent pathways from GTP-primed to G-ACT-2/3 were obtained by decomposing the net reactive flux into individual pathways.

For a pathway *P* = (*s*_1_, *s*_2_,…, *s_k_*), with *s*_1_ ∈ *A* and *s_k_* ∈ *B* over all MSMs, the pathway flux is defined as:

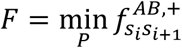

corresponding to the bottleneck flux along the pathway. This analysis identified the dominant ligand-specific reaction pathways across the MSM ensemble. Supplementary Table 4 summarizes the ligand-dependent pathways, the fraction of models in which each pathway was observed, the path flux fraction over the total flux of all ligand-specific models, and the mean MFPT (with associated uncertainties). These ligand-dependent pathways were then used to interpret the kinetic networks governing the GTP-primed to G-ACT-2/3 transition (Figure 4).

### Geometric Analysis of MOR–G_i1_ Structural Ensembles

Geometric analyses were performed to quantify a set of MOR and G-protein structural descriptors previously shown to distinguish the experimentally characterized GTP-primed, G-ACT-1, and G-ACT-2/3 states^11^. Analyses were conducted on receptor-aligned structural ensembles representing each macrostate, consisting of 100 randomly selected structures assigned to the corresponding state for both LFT- and MP-bound MOR–Gα_i1_ complexes. For each ensemble, all structures were first superposed onto the ligand-specific reference structure (the corresponding GTP-primed cryo-EM structure) using MOR backbone atoms (N, Cα, C, and O).

Receptor conformational changes were quantified using MOR TM helices and loop regions defined according to the Uniprot OPRM1_MOUSE residue numbering scheme.^17^ TM helices were defined as TM1: M65^1×29^-Y96^1×60^, TM2: T101^2×37^-G131^2×67^, TM3:G136^3×21^-H171^3×56^, TM4: T180^4×38^-A206^4×63^, TM5: P224^5×31^-V262^5×68^, TM6: S268^6×23^-I306^6×61^, TM7: T311^7×29^-L339^7×56^, and H8: D340^8×47^-I352^8×59^. The intracellular TM4-TM6 distance variation was measured as the Cα–Cα distance between residues R182^4×40^ and R273^6×28^.

Gα_i1_ structural descriptors were calculated, with residue numbering defined according to the Uniprot GNAI1_HUMAN sequence.^18^ Regions were defined as the αN helix: G2^G.HN.2^– K29^G.HN.53^, AHD: E63 ^H.HA.01^–R176 ^H.HF.06^, RHD: E33^G.S1.01^–H57^G.H1.12^ and I184^H.S2.01–^ D328^G.s6h5.05^, and α5 helix: T329^G.H5.01–^354^G.H5.26^. RMSDs of the RHD and α5 helix were calculated relative to the reference structure after receptor-based alignment. AHD opening was calculated by first aligning the RHD of each structure to the reference RHD, and then measuring the angle formed by the reference AHD center, the reference RHD center, and the AHD center of the current structure.

Receptor–G-protein coupling was quantified by counting residue-residue contacts between G_αi1_ and MOR using a closest-heavy-atom distance cutoff of 4.5 Å. Contact frequencies were calculated as the fraction of structures in which a given residue pair satisfied the distance criterion.

Specific MOR–G protein interface distances were also measured as closest-heavy-atom distances between selected residue pairs. These included: (i) MOR ICL3 interactions with Gα or Gβ subunits: MOR L265–Gα R32^G.hns1.03^, MOR L265–Gα L37 ^G.S1.05^, MOR L265–Gα T219^G.h2s4.05^, MOR L265 –Gα N347^G.H5.19^, MOR S266 –Gα A31^G.hns1.02^, MOR S266–Gα G217^G.h2s4.03^, MOR G267 –Gα A31^G.hns1.02^, MOR G267–Gα G217 ^G.h2s4.03^, MOR S268^6×23^–Gα E216^G.h2s4.02^, MOR S268^6×23^–Gα G217^G.h2s4.03^, and MOR S268^6×23^–Gβ A56; and (ii) MOR TM5/TM6 interactions with the C-terminal region of the Gα α5 helix: MOR I256^5×62^–Gα L353^G.H5.25^, MOR I256^5×62^–Gα F354^G.H5.26^), MOR K269^6×24^–Gα C351^G.H5.23^, MOR R273^6×28^–Gα C351^G.H5.23^, MOR R276^6×31^–Gα D350^G.H5.22^, and MOR R280^6×35^–Gα F354^G.H5.26^.

Helicity was calculated from backbone dihedral angles. Residues were classified as helical when their φ angle was between −120° and 0° and ψ angle was between −90° and 15°. Helicity was reported for MOR ICL2 and ICL3, as well as the Gα_i1_ αN and α5 helices, as the fraction of residues satisfying these criteria. Changes were interpreted relative to the corresponding reference state.

Within each ligand–macrostate group, the distribution of each geometric descriptor was visualized using violin plots, with ligand-specific mean values overlaid.

### Generation of Gαβ density maps from simulated I1–I4 states of the LFT-MOR–G_i1_ complex

Ligand-specific Gαβ density maps were generated for the LFT–MOR–G_i1_ ensemble using equal-count bootstrap MSMs constructed at a lag time of 20 ns. For each ligand and microstate, 100 assigned MD frames were sampled uniformly using a fixed random seed (13), with replacement when fewer than 100 frames were available. Each sampled structure was superposed on its ligand-specific G-ACT-1 reference using matched Gαβ backbone atoms (N, Cα, C, and O).

A density map was computed for each microstate using the aligned coordinates of non-hydrogen Gα_i1_ atoms. Atoms were weighted by atomic number and projected onto a common (280 × 280 × 280) Cartesian grid with a voxel spacing of 0.96 Å. The resulting densities were Gaussian-smoothed using `scipy.ndimage.gaussian_filter`,^61^ to a nominal resolution of 4.0 Å, and each microstate map was normalized to unit integrated density.

Only bootstrap MSMs supporting a transition-path-theory pathway from the GTP-Primed state to either G-ACT-2 or G-ACT-3 were retained. G-ACT-2 and G-ACT-3 were subsequently merged into a single G-ACT-2/3 macrostate. For each ligand (ℓ) and macrostate (*M*), the normalized microstate density maps were averaged according to their stationary populations across the retained bootstrap MSMs:

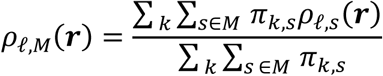

where (*ρ*_l,*s*_) is the normalized density map of microstate *s* for ligand ℓ, and *π_k_*_,*s*_ is its stationary population of microstate *s* in bootstrap MSM *k*. Final macrostate density maps were written in MRC format on a common grid.

### 3D variability analysis

CryoSPARC v4.7.1^62^ was used for all cryo-EM analyses. For mask generation, the LFT-MOR-G_i1_ G-ACT-1 state was used as the consensus starting map. The receptor and micelle densities were excluded so that the mask encompassed only the G-protein complex (Gα, Gβ, and Gγ). To minimize model bias and better capture conformational motions during 3D variability analysis (3DVA), the mask was intentionally expanded using a dilation radius of 8 pixels and a soft padding width of 12 pixels, resulting in a total outward extension of 20 pixels (19.2 Å at a pixel size of 0.96 Å).

To characterize conformational variability within the G-ACT-1–3-like population, 3DVA was first performed using the simple option with default parameters, generating six variability components, each represented by 20 frames for visualization. Approximately 1.6 million particles from the LFT-MOR-G_i1_ dataset that had previously been assigned to G-ACT-1–3-like states^11^ were analyzed. Visual inspection of the resulting trajectories indicated that the sixth component captured the most prominent conformational changes, particularly the large-scale motions of the Gα AHD and the α5 helix that distinguish the G-ACT-1–3 states (Supplementary Movie 1). To resolve discrete conformational substates along this trajectory, 3DVA was subsequently performed along the same sixth component using the cluster option with default parameters, yielding 18 discrete particle clusters. Each cluster was then subjected to non-uniform refinement, and representative maps placed along the trajectory are shown in Supplementary Figure 3. The corresponding map resolutions ranged from 5.6 to 8.2 Å. Cluster occupancies were calculated as the fraction of particles assigned to each cluster relative to the total number of particles included in the 3DVA.

To compare the Gαβ density maps from the simulated LFT–MOR–G_i1_ microstates with the 3DVA-derived cryo-EM cluster maps, contour levels were first selected by visual inspection. The maps were then manually positioned in ChimeraX, and the default Fit option was used to optimize the alignment and obtain cross-correlation values.

## Acknowledgments

This work was supported by NIH grant R01DA063209 (to M.F.) and the St. Jude Children’s Research Hospital Collaborative Research Consortium on GPCRs (G.S.). We thank Michael J. Robertson for sharing the cryo-EM data prior to publication. Computational work was supported in part through the computational resources and staff expertise provided by Scientific Computing at the Icahn School of Medicine at Mount Sinai and supported by the Clinical and Translational Science Awards (CTSA) grant UL1TR004419 from the National Center for Advancing Translational Sciences. Additionally, Anton 3 computer time was provided by the Pittsburgh Supercomputing Center (PSC) through Grant 1R24GM154042 from the National Institutes of Health. The Anton 3 machine at PSC is made available by D. E. Shaw Research. Research reported in this paper was also supported by the Office of Research Infrastructure of the National Institutes of Health under award number S10OD026880 and S10OD030463. The content is solely the responsibility of the authors and does not necessarily represent the official views of the National Institutes of Health. For cryo-EM data analysis, we acknowledge the St. Jude High-Performance Computing Facility (HPCF) research cluster and its staff for providing the computational resources and infrastructure that supported this work.

