## Supplementary Material for "Transition Intermediates Encode Ligand Efficacy at the μ-Opioid Receptor-Gi Protein Complex"

Supplementary Figure 1……………………………………………………………pg. 2
Supplementary Figure 2……………………………………………………………pg. 3
Supplementary Figure 3……………………………………………………………pg. 4
Supplementary Figure 4……………………………………………………………pg. 5
Supplementary Figure 5……………………………………………………………pg. 6
Supplementary Figure 6……………………………………………………………pg. 7
Supplementary Figure 7……………………………………………………………pg. 8
Supplementary Figure 8……………………………………………………………pg. 9
Supplementary Figure 9……………………………………………………………pg. 10
Supplementary Figure 10…………………………………………………………...pg. 11
Supplementary Figure 11…………………………………………………………...pg. 12
Supplementary Table 1……………………………………………………………...pg. 13
Supplementary Table 2……………………………………………………………...pg. 14
Supplementary Table 3……………………………………………………………...pg. 20
Supplementary Table 4……………………………………………………………...pg. 21
Supplementary Table 5……………………………………………………………...pg. 22
Supplementary Table 6……………………………………………………………...pg. 23
Supplementary Table 7……………………………………………………………...pg. 24
Supplementary Table 8……………………………………………………………...pg. 25
Supplementary Table 9……………………………………………………………...pg. 26

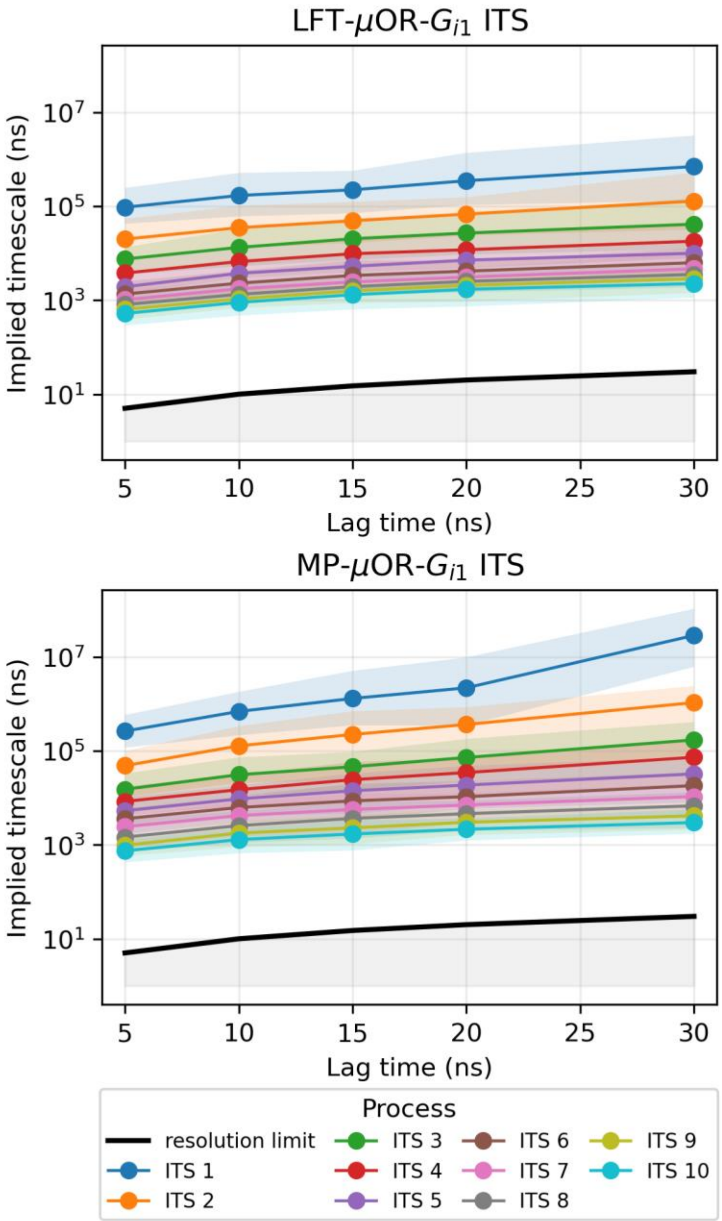


**Supplementary Figure 1.** Implied timescales (ITS) for the ten slowest processes as a function of lag time for LFT-MOR-$G_{i1}$ (top) and MP-MOR-$G_{i1}$ (bottom) complexes. Shaded regions indicate the 95% confidence intervals computed across bootstrap samples. The region below the lag-time, which represents the resolution limit of the model, is indicated in gray.

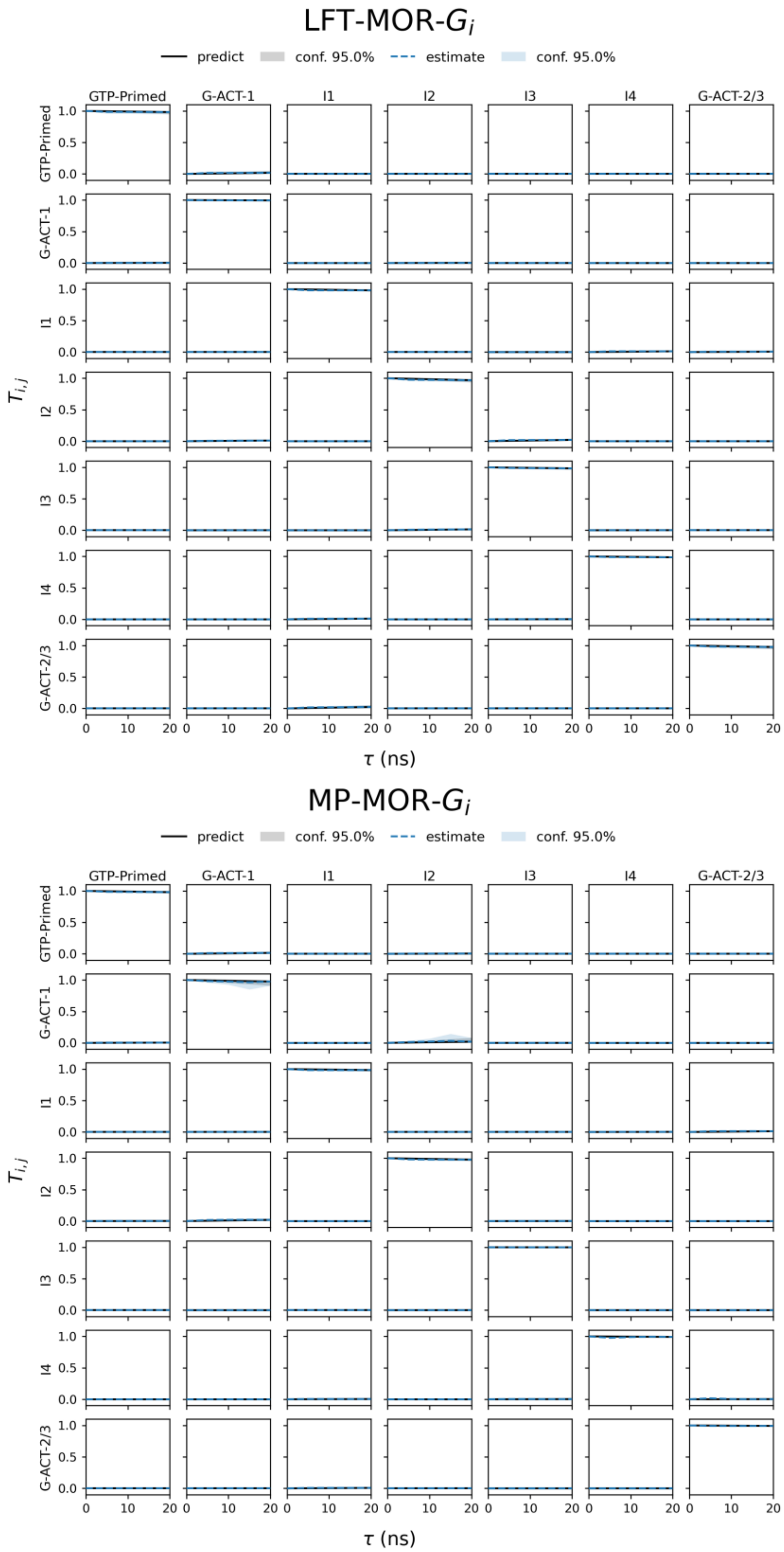


**Supplementary Figure 2.** Chapman-Kolmogorov tests for the final 20 ns-lag 7-macrostate kinetic activation models for the (top) LFT-MOR-$G_{i1}$ and (bottom) MP-MOR-$G_{i1}$ complexes.

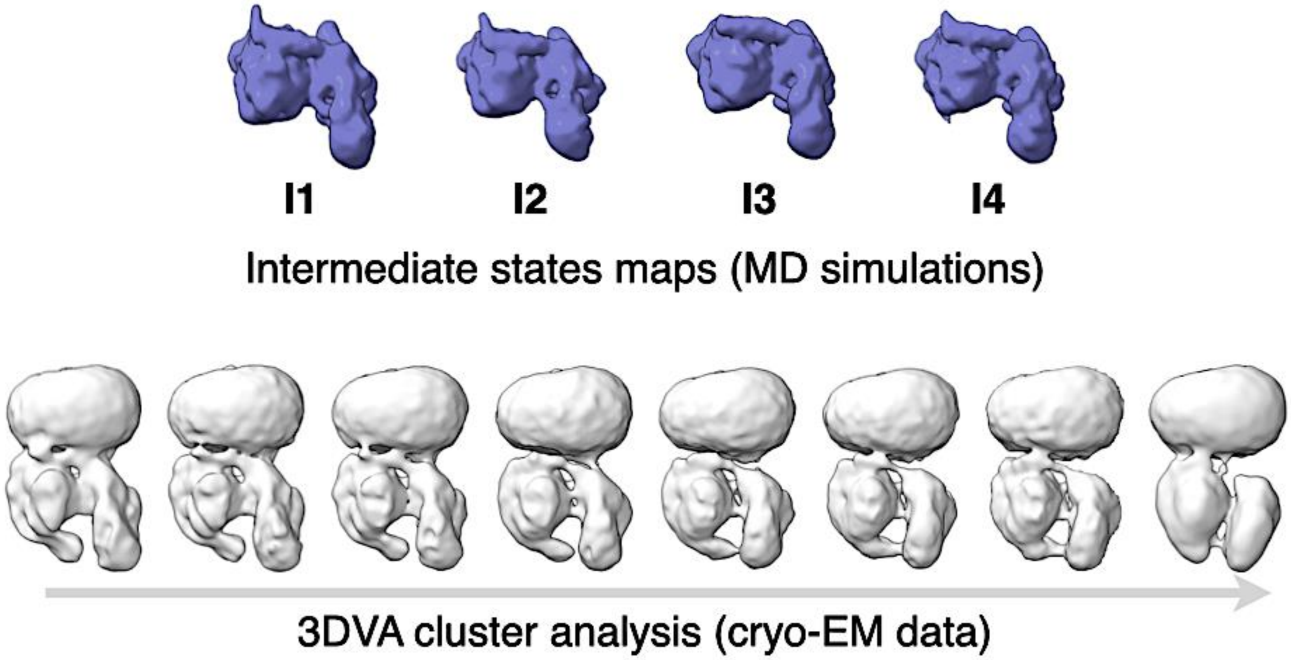


**Supplementary Figure 3. Representative density maps of intermediate states derived from MSM analysis of MD simulations and 3DVA-derived cryo-EM cluster maps.** Top row shows representative Gα-Gβ density maps generated from the all-atom microstates corresponding to the I1-I4 states identified by MSM analysis of MD simulations of the LFT-MOR-$G_{i1}$ complex. Bottom row shows density maps obtained after non-uniform refinement of cryo-EM cluster maps from 3DVA analysis (see Methods). Out of the 18 discrete cluster maps, only eight representative maps are shown here to illustrate the transition (see the Supporting Movie 1 showing all 18 clusters). The grey arrow indicates the transition from G-ACT-1-like to G-ACT-2/3-like states, with intermediate states I1-I4 in between.

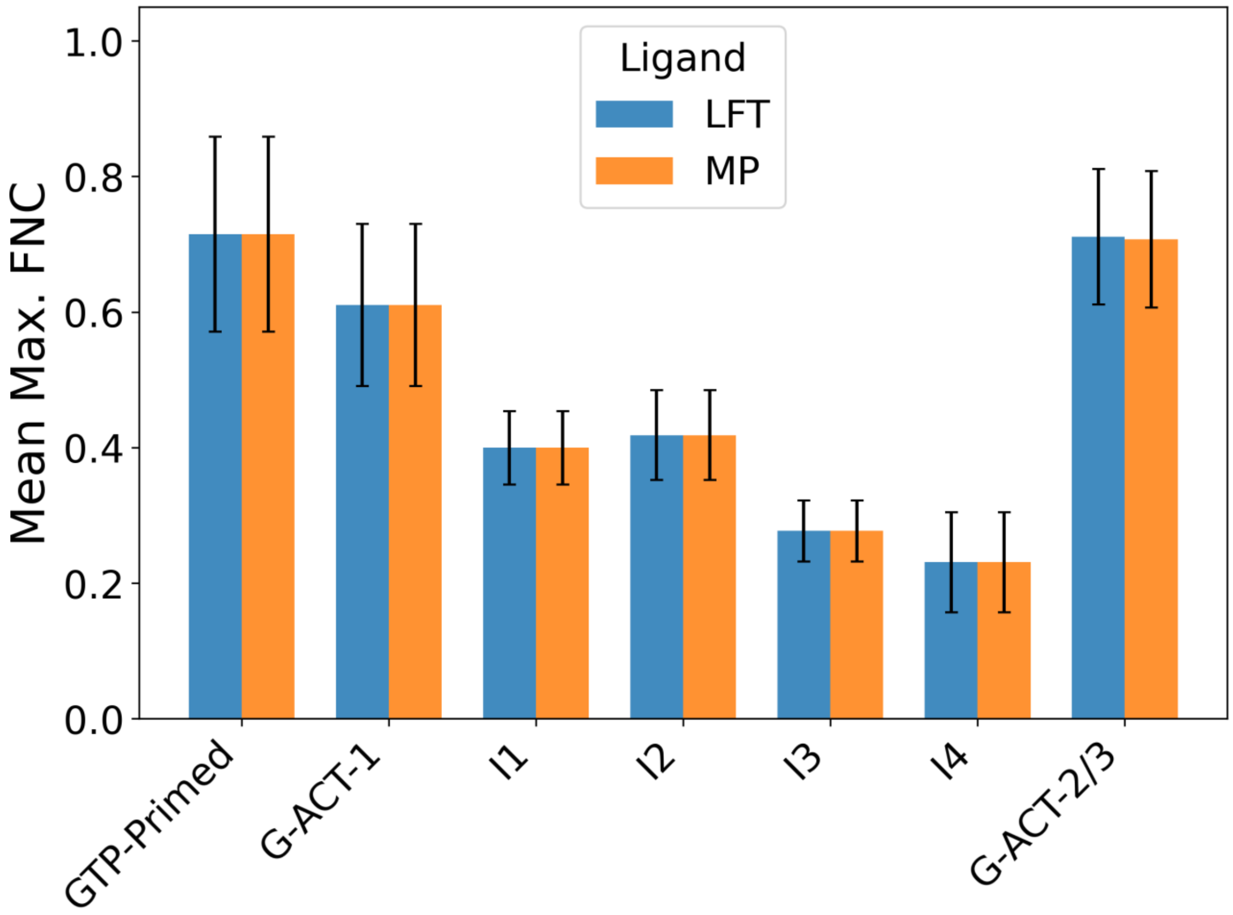


**Supplementary Figure 4. Mean maximum FNC per macrostate.** Mean maximum FNC values with respect to available cryo-EM structures across microstate frames assigned to each macrostate assignment (see Methods) are shown for LFT-MOR-$G_{i1}$ and MP-MOR-$G_{i1}$ systems (blue and orange, respectively). Error bars indicate the standard deviation across macrostate-assigned microstates. Cryo-EM-like (GTP-primed, G-ACT-1, and G-ACT-2/3) macrostates display mean maximum FNC values above 60%, whereas intermediate states (I1-I4) remain below 45%, supporting their divergence from experimentally characterized conformations.

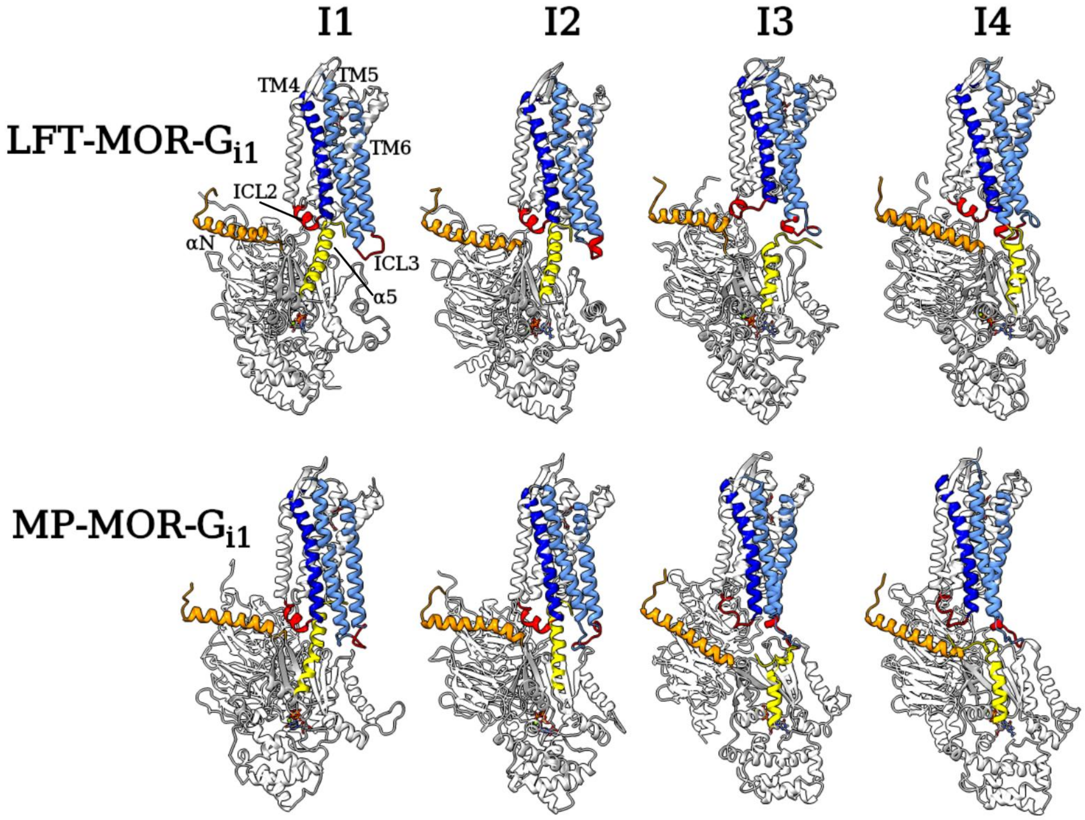


**Supplementary Figure 5. Representative structures of the intermediate states I1-I4 identified along the transition from GTP-primed to G-ACT-2/3.** I1-I4 representative structures aligned by MOR for LFT-MOR-$G_{i1}$ (top) and MP-MOR-$G_{i1}$ (bottom)**.** Structural elements including motifs that show significant difference between the two ligand-bound systems in at least one intermediate macrostate (see Supplementary Figure 6) are colored; MOR: TM5 and 6 (light blue), TM4 (blue), ICL2 and ICL3 (red). $G\alpha_{i1}$: αN (orange) and α5 (yellow).

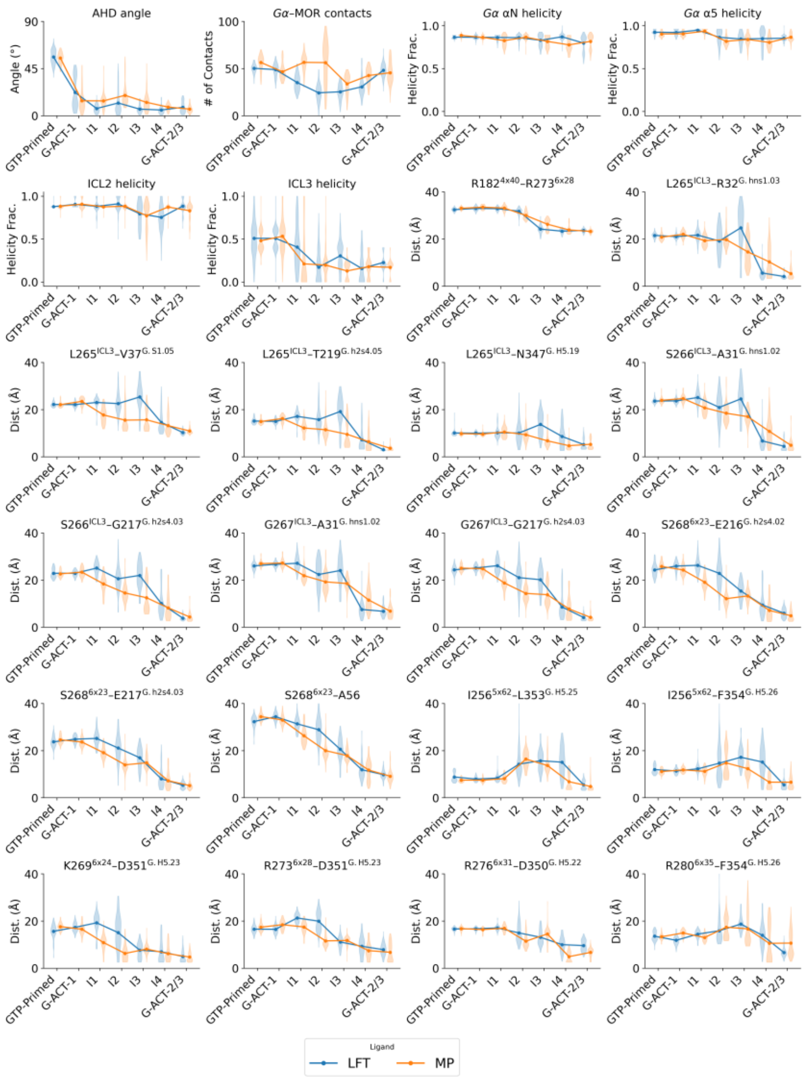


**Supplementary Figure 6: Selected MOR and $G\alpha_{i1}$ Structural Descriptors Calculated for each Macrostate Along the Transition Pathway from GTP-primed to G-ACT-2/3.** Violin plots with the line showing the trend of the mean of structural features previously shown to distinguish the experimentally characterized GTP-primed, G-ACT-1, and G-ACT-2/3 states (Robertson, MJ et al. Nature 652(8110):794-802) in MP- (orange) or LFT-bound (blue) MOR-$G_{i1}$ for each macrostate along the transition pathway from GTP-primed to G-ACT-2/3. Panels are in the order of their reference in the text. From top left to bottom right: Gα-AHD opening angle; number of MOR-Gα contacts; helicity of Gα αN, Gα α5, MOR ICL2, and MOR ICL3; TM4-TM6 interaction: MOR R182$^{4\times40}$–MOR R273$^{6\times28}$; MOR ICL3 interactions with Gα or Gβ subunits: MOR L265–Gα R32$^{G.hns1.03}$, MOR L265–Gα L37$^{G.S1.05}$, MOR L265–Gα T219$^{G.h2s4.05}$, MOR L265–Gα N347$^{G.H5.19}$, MOR S266–Gα A31$^{G.hns1.02}$, MOR S266–Gα G217$^{G.h2s4.03}$, MOR G267–Gα A31$^{G.hns1.02}$, MOR G267–Gα G217$^{G.h2s4.03}$, MOR S268$^{6\times23}$–Gα E216$^{G.h2s4.02}$, MOR S268$^{6\times23}$–Gα E217$^{G.h2s4.03}$, and MOR S268$^{6\times23}$–Gβ A56; MOR TM5/TM6 interactions with the C-terminal region of the Gα α5 helix: MOR I256$^{5\times62}$–Gα L353$^{G.H5.25}$, MOR I256$^{5\times62}$–Gα F354$^{G.H5.26}$, MOR K269$^{6\times24}$–Gα D351$^{G.H5.23}$, MOR R273$^{6\times28}$–Gα D351$^{G.H5.23}$, MOR R276$^{6\times31}$–Gα D350$^{G.H5.22}$, and MOR R280$^{6\times35}$–Gα F354$^{G.H5.26}$.

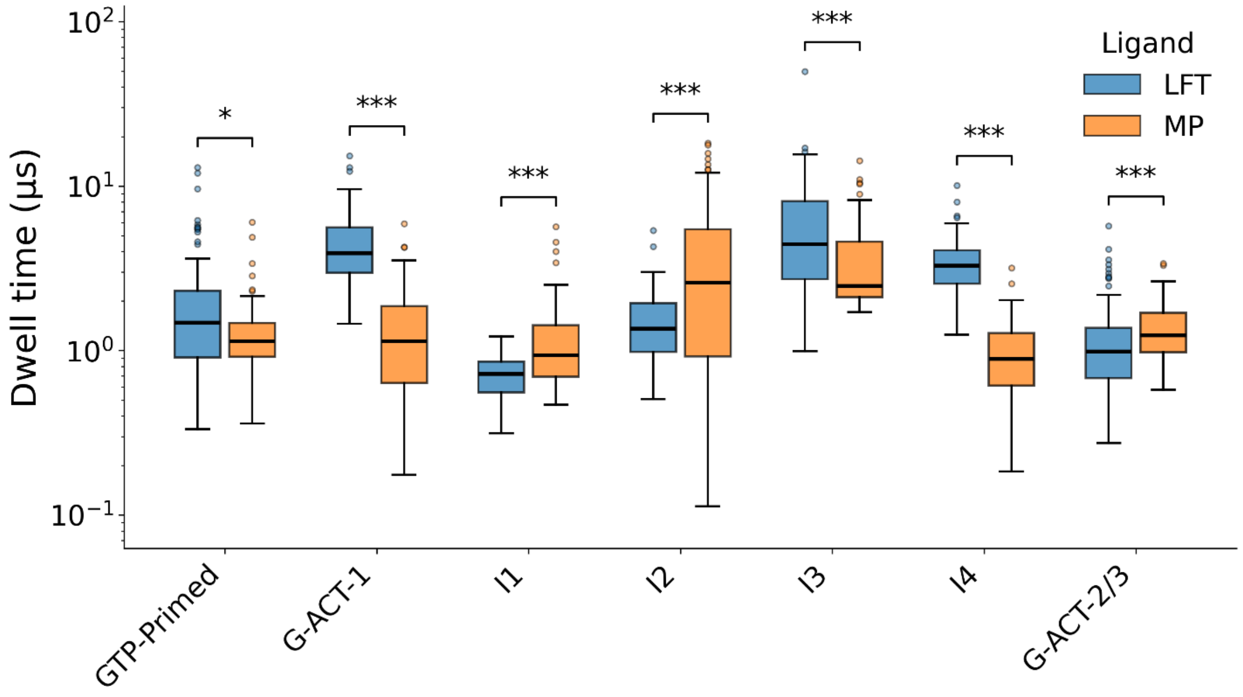


**Supplementary Figure 7: Ligand-Specific Median Dwell Times for MOR-G$_{i1}$ Macrostates.** Median dwell times across bootstrap samples for each macrostate are shown for LFT-MOR-G$_{i1}$ and MP-MOR-G$_{i1}$ systems (blue and orange, respectively). Median value difference statistical significance was assessed using the Mann-Whitney U test, with *, and *** denoting $p < 0.05$ and $p < 0.001$, respectively.

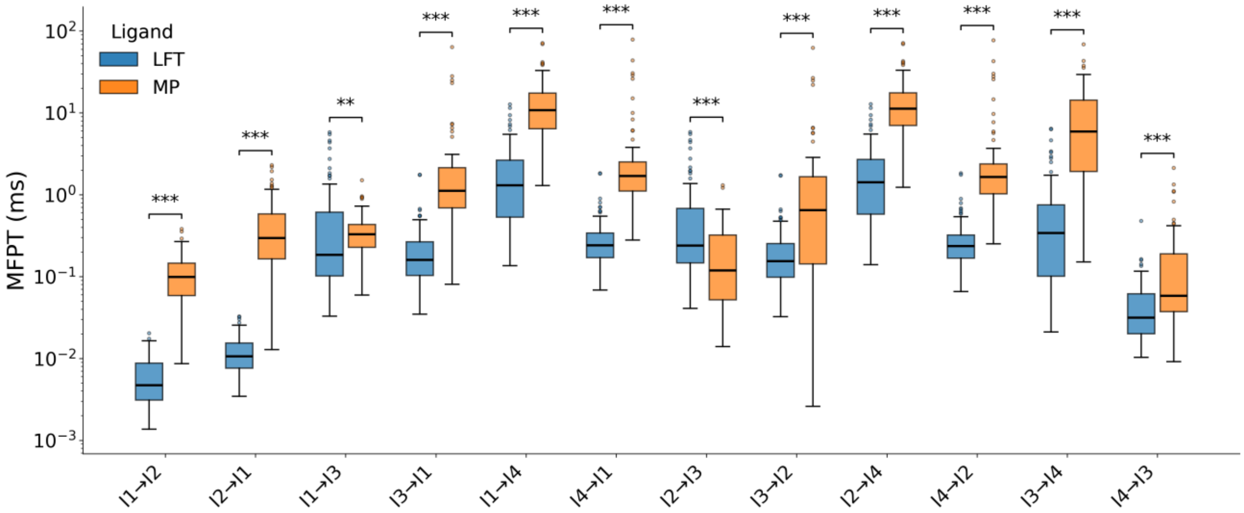


**Supplementary Figure 8. Mean first-passage times between intermediate macrostates.** MFPTs (in ms) for all pairwise transitions among the four intermediate macrostates (I1-I4) are shown for LFT-MOR-$G_{i1}$ and MP-MOR-$G_{i1}$ (blue and red, respectively). Box plots indicate the distribution of MFPTs from bootstrap sampling. The center line indicates the median, the box spans the interquartile range (IQR), the whiskers extend to 1.5×IQR, and outliers are shown as open circles. Median value difference statistical significance was assessed using the Mann-Whitney U test, with ** and *** denoting $p < 0.01$ and $p < 0.001$, respectively.

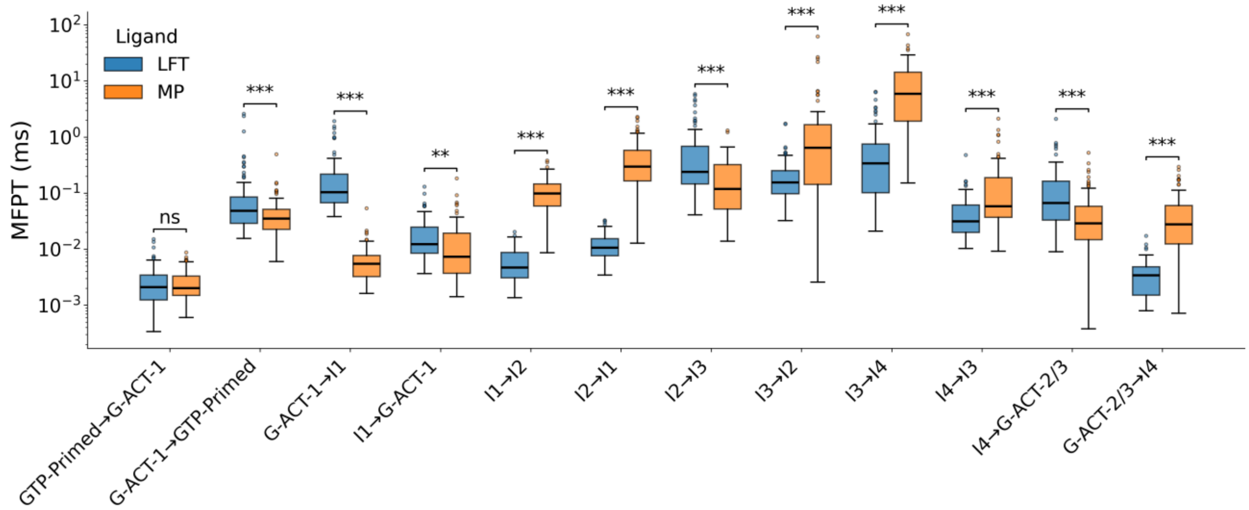


**Supplementary Figure 9. Mean first-passage times between pathway-relevant macrostates.** MFPTs (in ms) for all pathway-relevant transitions among the macrostates GTP-primed, G-ACT-1, I1-I4, G-ACT-2/3) are shown for LFT-MOR-$G_{i1}$ and MP-MOR-$G_{i1}$ (blue and orange, respectively). Box plots indicate the distribution of MFPTs from bootstrap sampling. The center line indicates the median, the box span the interquartile range (IQR), the whiskers extend to 1.5×IQR, and outliers are shown as open circles. Median value difference statistical significance was assessed using the Mann-Whitney U test, with ** and *** denoting $p < 0.01$ and $p < 0.001$, respectively. ns denotes no statistically significant difference.

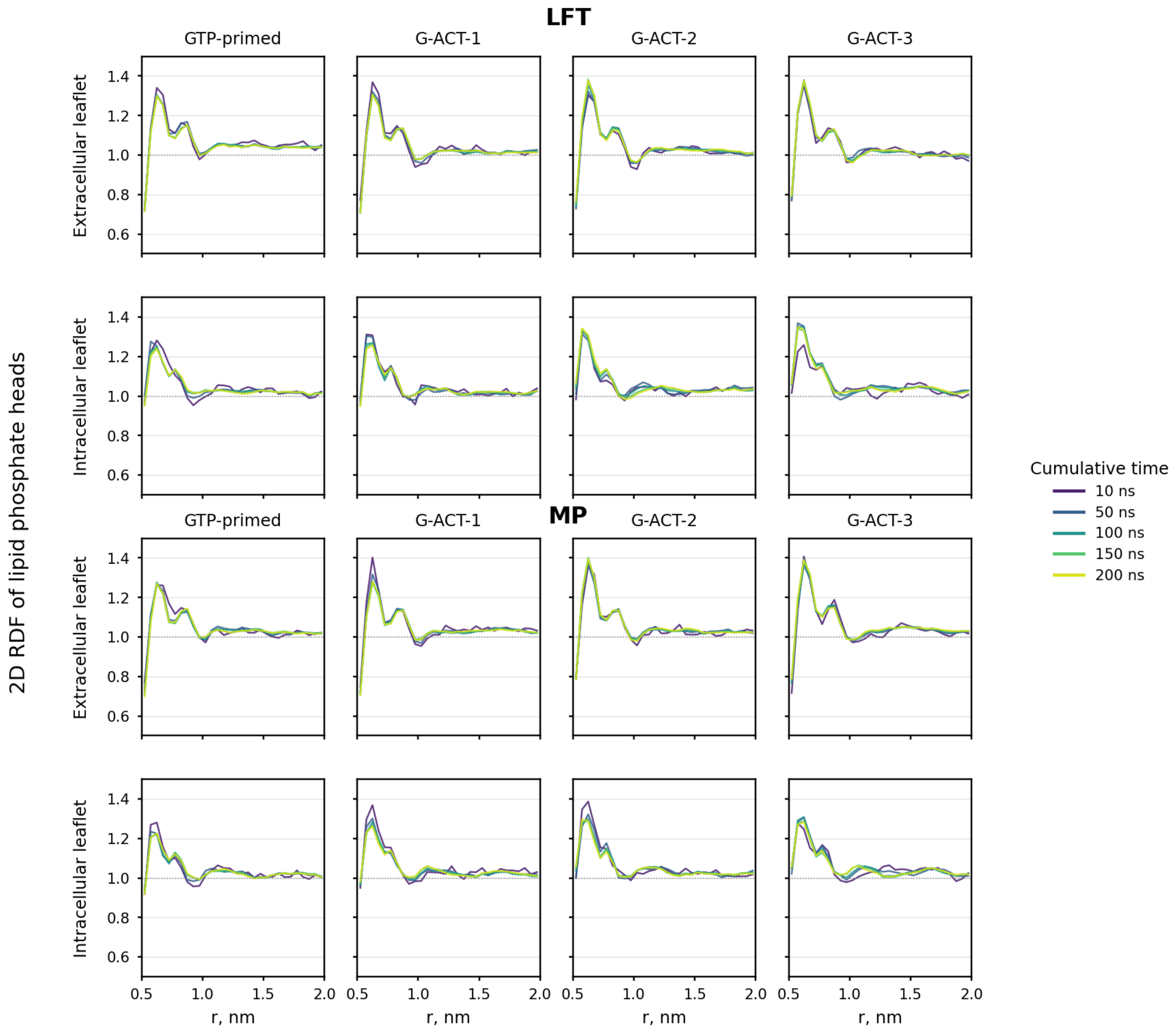


**Supplementary Figure 10. Equilibration of the membrane.** Radial distribution functions of lipid head groups in the XY-plane are shown for all starting conformations during equilibration for LFT-MOR-$G_{i1}$ (row 1 and 2) and MP MOR-$G_{i1}$ (row 3 and 4).

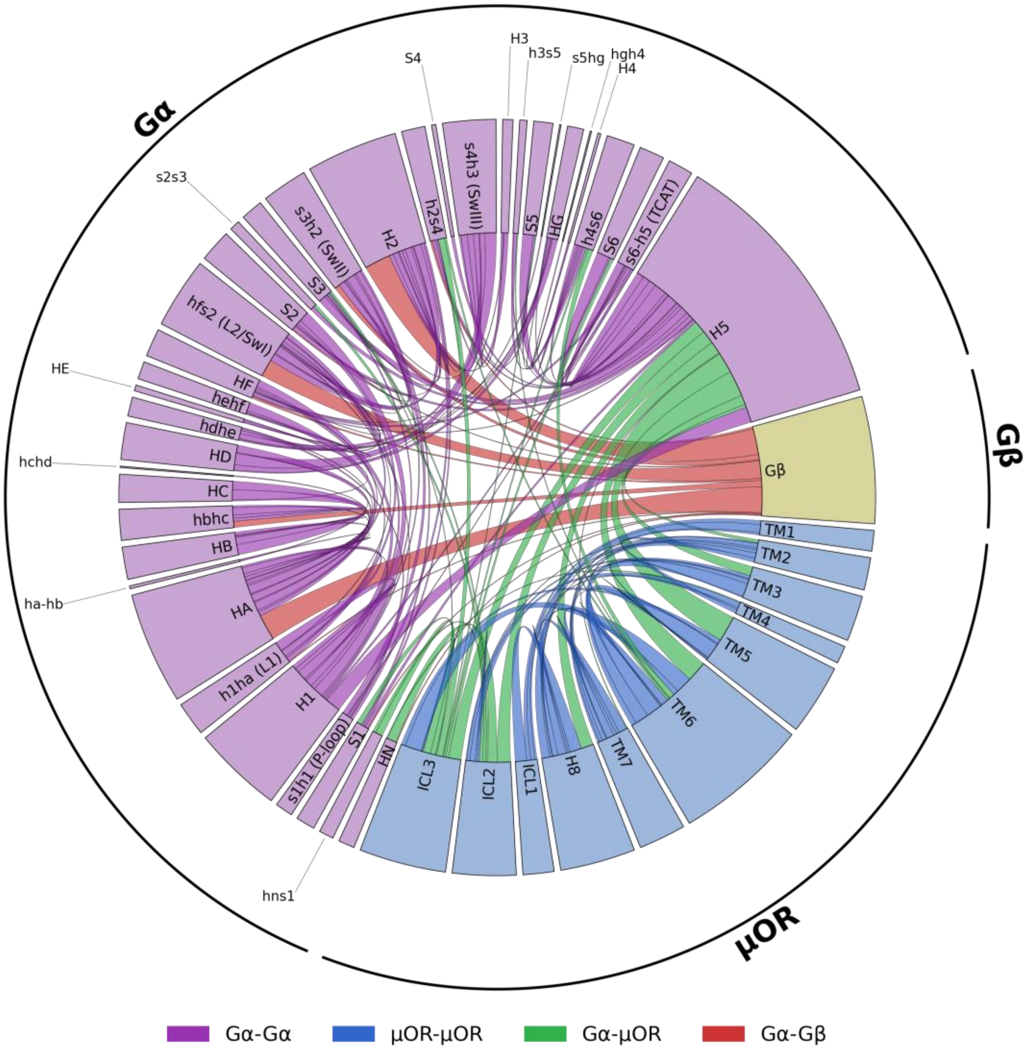


**Supplementary Figure 11. Map of features used to estimate MSMs during adaptive sampling.** The 685 Cα-Cα distance features used to construct the MSMs are depicted as connections between structural elements of each protein chain within the complex. Line thickness is proportional to the number of features associated with each connection.

**Supplementary Table 1. Overview of adaptive sampling rounds.**

| | **LFT-MOR-$G_{i1}$** | | | | | **MP-MOR-$G_{i1}$** | | | | |
|---|---|---|---|---|---|---|---|---|---|---|
| | **GROMACS** | | **Anton** | | | **GROMACS** | | **Anton** | | |
| **Round** | # Traj. | Time [µs] | # Traj. | Time [µs] | **Total nb. Time [µs]** | # Traj. | Time [µs] | # Traj. | Time [µs] | **Total nb. Time [µs]** |
| Initial nb. | 155 | 148 | 4 | 40 | **188** | 77 | 94.2 | 4 | 40 | **134.2** |
| #1$^{pul.}$ | 10 | 0.62 | - | - | **-** | 9 | 0.88 | - | - | **-** |
| #1$^{nb.}$ | 198 | 171 | - | - | **171** | 291 | 77.3 | 4 | 20 | **97.3** |
| #2$^{pul.}$ | 16 | 0.31 | - | - | **-** | 11 | 0.07 | - | - | **-** |
| #2$^{nb.}$ | 270 | 75 | 18 | 90 | **165** | 164 | 22.5 | 4 | 17 | **39.5** |
| #3$^{pul.}$ | 14 | 0.09 | - | - | **-** | 7 | 0.02 | - | - | **-** |
| #3$^{nb.}$ | 180 | 18.6 | - | - | **18.6** | 226 | 63.5 | 5 | 60 | **123.5** |
| #4$^{pul.}$ | 3 | 0.01 | - | - | **-** | 33 | 0.04 | - | - | **-** |
| #4$^{nb.}$ | 21 | 3.21 | - | - | **3.21** | 213 | 15.6 | - | - | **15.6** |
| **Total nb.** | 824 | 415.8 | 22 | 130 | **545.8** | 971 | 273.1 | 17 | 137 | **410.1** |
| Total pul. | 43 | 1.03 | 0 | 0 | 1.03 | 60 | 1.01 | 0 | 0 | 1.01 |

*nb.: non-biased simulations*
*pul.: pulling simulations*

**Supplementary Table 2. Shared featurization used to build production MSMs.**

| MSM Contact Features | |
|---|---|
| **Residue i** | **Residue j** |
| Gα-E145$^{\text{H.HD.12}}$ | Gα-V233$^{\text{G.s4h3.07}}$ |
| Gα-E145$^{\text{H.HD.12}}$ | Gα-L232$^{\text{G.s4h3.06}}$ |
| Gα-R144$^{\text{H.HD.11}}$ | Gα-D231$^{\text{G.s4h3.05}}$ |
| Gα-R144$^{\text{H.HD.11}}$ | Gα-L232$^{\text{G.s4h3.06}}$ |
| Gα-E145$^{\text{H.HD.12}}$ | Gα-L234$^{\text{G.s4h3.08}}$ |
| Gα-R144$^{\text{H.HD.11}}$ | Gα-V233$^{\text{G.s4h3.07}}$ |
| Gα-Q147$^{\text{H.hdhe.02}}$ | Gα-A235$^{\text{G.s4h3.09}}$ |
| Gα-E145$^{\text{H.HD.12}}$ | Gα-A235$^{\text{G.s4h3.09}}$ |
| Gα-K70$^{\text{H.HA.08}}$ | Gβ-R134 |
| Gα-Q68$^{\text{H.HA.06}}$ | Gβ-V133 |
| Gα-Q68$^{\text{H.HA.06}}$ | Gβ-N132 |
| Gα-A71$^{\text{H.HA.09}}$ | Gβ-R134 |
| Gα-A71$^{\text{H.HA.09}}$ | Gβ-S136 |
| Gα-Q68$^{\text{H.HA.06}}$ | Gβ-R134 |
| Gα-Y69$^{\text{H.HA.07}}$ | Gβ-R134 |
| Gα-K67$^{\text{H.HA.05}}$ | Gβ-R134 |
| Gα-V72$^{\text{H.HA.10}}$ | Gβ-S136 |
| Gα-A71$^{\text{H.HA.09}}$ | Gβ-V135 |
| Gα-E65$^{\text{H.HA.03}}$ | Gβ-P94 |
| Gα-V72$^{\text{H.HA.10}}$ | Gβ-V135 |
| Gα-Y69$^{\text{H.HA.07}}$ | Gβ-V135 |
| Gα-A30$^{\text{G.hns1.01}}$ | MOR-G267 |
| Gα-V72$^{\text{H.HA.10}}$ | Gβ-R137 |
| Gα-Q68$^{\text{H.HA.06}}$ | Gβ-V135 |
| Gα-A31$^{\text{G.hns1.02}}$ | MOR-G267 |
| Gα-R176$^{\text{H.HF.06}}$ | Gα-T327$^{\text{G.s6h5.04}}$ |
| Gα-T340$^{\text{G.H5.12}}$ | MOR-V173$^{\text{34x51}}$ |
| Gα-L175$^{\text{H.HF.05}}$ | Gα-T327$^{\text{G.s6h5.04}}$ |
| Gα-Q172$^{\text{H.HF.02}}$ | Gα-T327$^{\text{G.s6h5.04}}$ |
| Gα-A30$^{\text{G.hns1.01}}$ | MOR-S266 |
| Gα-T340$^{\text{G.H5.12}}$ | MOR-P172$^{\text{34x50}}$ |
| Gα-A31$^{\text{G.hns1.02}}$ | MOR-S266 |
| Gα-E33$^{\text{G.S1.01}}$ | MOR-S266 |
| Gα-G217$^{\text{G.h2s4.03}}$ | MOR-G267 |
| Gα-K349$^{\text{G.H5.21}}$ | MOR-N342$^{\text{8x49}}$ |
| Gα-G217$^{\text{G.h2s4.03}}$ | MOR-S268$^{\text{6x23}}$ |
| Gα-R32$^{\text{G.hns1.03}}$ | MOR-S266 |
| Gα-I343$^{\text{G.H5.15}}$ | MOR-V173$^{\text{34x51}}$ |
| Gα-I344$^{\text{G.H5.16}}$ | MOR-A168$^{\text{3x53}}$ |
| Gα-E216$^{\text{G.h2s4.02}}$ | MOR-S268$^{\text{6x23}}$ |
| Gα-E216$^{\text{G.h2s4.02}}$ | MOR-G267 |
| Gα-I343$^{\text{G.H5.15}}$ | MOR-P172$^{\text{34x50}}$ |
| Gα-D350$^{\text{G.H5.22}}$ | MOR-T101$^{\text{2x37}}$ |

| MSM Contact Features | |
|---|---|
| **Residue i** | **Residue j** |
| Gα-V179$^{G.hfs2.03}$ | Gβ-E138 |
| Gα-K35$^{G.S1.03}$ | MOR-S266 |
| Gα-G217$^{G.h2s4.03}$ | MOR-S266 |
| Gα-D315$^{G.h4s6.09}$ | MOR-K271$^{6x26}$ |
| Gα-V34$^{G.S1.02}$ | MOR-S266 |
| Gα-I344$^{G.H5.16}$ | MOR-P172$^{34x50}$ |
| Gα-D193$^{G.s2s3.02}$ | MOR-V173$^{34x51}$ |
| Gα-V218$^{G.h2s4.04}$ | MOR-G267 |
| Gα-C351$^{G.H5.23}$ | MOR-T103$^{2x39}$ |
| Gα-V218$^{G.h2s4.04}$ | MOR-S266 |
| Gα-I344$^{G.H5.16}$ | MOR-V169$^{3x54}$ |
| Gα-E33$^{G.S1.01}$ | MOR-L265 |
| Gα-A31$^{G.hns1.02}$ | MOR-L265 |
| Gα-R32$^{G.hns1.03}$ | MOR-L265 |
| Gα-L194$^{G.S3.01}$ | MOR-V173$^{34x51}$ |
| Gα-D350$^{G.H5.22}$ | MOR-N342$^{8x49}$ |
| Gα-R32$^{G.hns1.03}$ | MOR-D177$^{34x55}$ |
| Gα-G352$^{G.H5.24}$ | MOR-N342$^{8x49}$ |
| Gα-N347$^{G.H5.19}$ | MOR-A168$^{3x53}$ |
| Gα-T48$^{G.H1.03}$ | Gα-R176$^{H.HF.06}$ |
| Gα-T48$^{G.H1.03}$ | Gα-L175$^{H.HF.05}$ |
| Gα-V34$^{G.S1.02}$ | MOR-L265 |
| Gα-G352$^{G.H5.24}$ | MOR-F343$^{8x50}$ |
| Gα-L348$^{G.H5.20}$ | MOR-R165$^{3x50}$ |
| Gα-A31$^{G.hns1.02}$ | MOR-M264 |
| Gα-R32$^{G.hns1.03}$ | MOR-M264 |
| Gβ-R48 | MOR-T97$^{12x48}$ |
| Gβ-R46 | MOR-T97$^{12x48}$ |
| Gα-D315$^{G.h4s6.09}$ | MOR-E270$^{6x25}$ |
| Gα-L348$^{G.H5.20}$ | MOR-A168$^{3x53}$ |
| Gα-T219$^{G.h2s4.05}$ | MOR-S266 |
| Gα-V179$^{G.hfs2.03}$ | Gβ-A140 |
| Gα-G352$^{G.H5.24}$ | MOR-E341$^{8x48}$ |
| Gα-D315$^{G.h4s6.09}$ | MOR-S268$^{6x23}$ |
| Gα-R32$^{G.hns1.03}$ | MOR-L176$^{34x54}$ |
| Gα-Q52$^{G.H1.07}$ | Gα-L175$^{H.HF.05}$ |
| Gβ-T47 | MOR-T97$^{12x48}$ |
| Gα-G352$^{G.H5.24}$ | MOR-D340$^{8x47}$ |
| Gα-N347$^{G.H5.19}$ | MOR-P172$^{34x50}$ |
| Gα-K54$^{G.H1.09}$ | Gα-Y61$^{G.h1ha.04}$ |
| Gα-G352$^{G.H5.24}$ | MOR-S261$^{5x67}$ |
| Gα-L353$^{G.H5.25}$ | MOR-N342$^{8x49}$ |
| Gα-C351$^{G.H5.23}$ | MOR-R263 |
| Gα-G352$^{G.H5.24}$ | MOR-K260$^{5x66}$ |
| Gα-R24$^{G.HN.48}$ | MOR-V173$^{34x51}$ |

| **MSM Contact Features** | |
|---|---|
| **Residue i** | **Residue j** |
| Gα-L353$^{G.H5.25}$ | MOR-E341$^{8x48}$ |
| Gα-G352$^{G.H5.24}$ | MOR-T103$^{2x39}$ |
| Gα-C351$^{G.H5.23}$ | MOR-S261$^{5x67}$ |
| Gα-T219$^{G.h2s4.05}$ | MOR-L265 |
| Gα-T316$^{G.h4s6.10}$ | MOR-M264 |
| Gα-K51$^{G.H1.06}$ | Gα-L175$^{H.HF.05}$ |
| Gα-G352$^{G.H5.24}$ | MOR-R263 |
| Gα-E318$^{G.h4s6.12}$ | MOR-M264 |
| Gα-C351$^{G.H5.23}$ | MOR-V262$^{5x68}$ |
| Gα-N347$^{G.H5.19}$ | MOR-V169$^{3x54}$ |
| Gα-L348$^{G.H5.20}$ | MOR-V169$^{3x54}$ |
| Gα-I55$^{G.H1.10}$ | Gα-Y61$^{G.h1ha.04}$ |
| Gα-C351$^{G.H5.23}$ | MOR-K260$^{5x66}$ |
| Gα-F354$^{G.H5.26}$ | MOR-E341$^{8x48}$ |
| Gα-C351$^{G.H5.23}$ | MOR-R165$^{3x50}$ |
| Gα-G352$^{G.H5.24}$ | MOR-V262$^{5x68}$ |
| Gα-L353$^{G.H5.25}$ | MOR-D340$^{8x47}$ |
| Gα-K54$^{G.H1.09}$ | Gα-G60$^{G.h1ha.03}$ |
| Gα-D350$^{G.H5.22}$ | MOR-R263 |
| MOR-L265 | MOR-L275$^{6x30}$ |
| Gα-K345$^{G.H5.17}$ | Gα-F354$^{G.H5.26}$ |
| MOR-K260$^{5x66}$ | MOR-S266 |
| Gα-D350$^{G.H5.22}$ | MOR-V262$^{5x68}$ |
| MOR-S64 | MOR-T312$^{7x28}$ |
| Gα-Q204$^{G.s3h2.03}$ | Gβ-T143 |
| Gα-Q204$^{G.s3h2.03}$ | Gβ-G144 |
| MOR-L265 | MOR-N274$^{6x29}$ |
| Gα-I55$^{G.H1.10}$ | Gα-G60$^{G.h1ha.03}$ |
| Gα-L353$^{G.H5.25}$ | MOR-R263 |
| Gα-L353$^{G.H5.25}$ | MOR-S261$^{5x67}$ |
| Gα-L353$^{G.H5.25}$ | MOR-K260$^{5x66}$ |
| Gα-F354$^{G.H5.26}$ | MOR-K271$^{6x26}$ |
| Gα-G352$^{G.H5.24}$ | MOR-R273$^{6x28}$ |
| Gα-T181$^{G.hfs2.05}$ | Gβ-I120 |
| Gα-Q204$^{G.s3h2.03}$ | Gβ-Y145 |
| Gα-G352$^{G.H5.24}$ | MOR-L259$^{5x65}$ |
| Gα-K54$^{G.H1.09}$ | Gα-A59$^{G.h1ha.02}$ |
| MOR-S266 | MOR-L275$^{6x30}$ |
| Gα-L348$^{G.H5.20}$ | Gα-F354$^{G.H5.26}$ |
| Gα-D341$^{G.H5.13}$ | MOR-V262$^{5x68}$ |
| MOR-L265 | MOR-K271$^{6x26}$ |
| Gα-N347$^{G.H5.19}$ | MOR-L265 |
| Gα-D341$^{G.H5.13}$ | MOR-R263 |
| Gα-C351$^{G.H5.23}$ | MOR-M264 |
| Gα-T182$^{G.hfs2.06}$ | Gα-E207$^{G.H2.03}$ |

| MSM Contact Features | |
|---|---|
| **Residue i** | **Residue j** |
| Gα-R205$^{G.H2.01}$ | Gβ-G144 |
| MOR-T103$^{2x39}$ | MOR-I278$^{6x33}$ |
| Gα-K349$^{G.H5.21}$ | Gα-F354$^{G.H5.26}$ |
| MOR-S261$^{5x67}$ | MOR-S266 |
| MOR-A168$^{3x53}$ | MOR-L275$^{6x30}$ |
| Gα-F354$^{G.H5.26}$ | MOR-K260$^{5x66}$ |
| MOR-S64 | MOR-V316$^{7x32}$ |
| MOR-L259$^{5x65}$ | MOR-L265 |
| Gα-I56$^{G.H1.11}$ | Gα-Q333$^{G.H5.05}$ |
| MOR-K260$^{5x66}$ | MOR-L265 |
| Gα-K54$^{G.H1.09}$ | Gα-E58$^{G.h1ha.01}$ |
| Gα-I55$^{G.H1.10}$ | Gα-A59$^{G.h1ha.02}$ |
| Gα-I56$^{G.H1.11}$ | Gα-T329$^{G.H5.01}$ |
| MOR-Y96$^{1x60}$ | MOR-C351$^{8x58}$ |
| Gα-G352$^{G.H5.24}$ | MOR-R276$^{6x31}$ |
| Gα-R205$^{G.H2.01}$ | Gβ-D163 |
| Gα-K180$^{G.hfs2.04}$ | Gα-G202$^{G.s3h2.01}$ |
| Gα-F354$^{G.H5.26}$ | MOR-N274$^{6x29}$ |
| MOR-V92$^{1x56}$ | MOR-C351$^{8x58}$ |
| Gα-G183$^{G.hfs2.07}$ | Gα-K210$^{G.H2.06}$ |
| Gα-L348$^{G.H5.20}$ | Gα-L353$^{G.H5.25}$ |
| Gα-L353$^{G.H5.25}$ | MOR-V262$^{5x68}$ |
| Gα-N347$^{G.H5.19}$ | MOR-R263 |
| Gα-T181$^{G.hfs2.05}$ | Gβ-N119 |
| MOR-T97$^{12x48}$ | MOR-F347$^{8x54}$ |
| Gα-T181$^{G.hfs2.05}$ | Gβ-D118 |
| MOR-R165$^{3x50}$ | MOR-T279$^{6x34}$ |
| MOR-S266 | MOR-K271$^{6x26}$ |
| Gα-Y61$^{G.h1ha.04}$ | Gα-C66$^{H.HA.04}$ |
| Gα-A326$^{G.s6h5.03}$ | Gα-V332$^{G.H5.04}$ |
| Gα-G42$^{G.s1h1.03}$ | Gα-Q204$^{G.s3h2.03}$ |
| Gα-L353$^{G.H5.25}$ | MOR-D272$^{6x27}$ |
| Gα-G183$^{G.hfs2.07}$ | Gα-E207$^{G.H2.03}$ |
| Gα-A41$^{G.s1h1.02}$ | Gα-G203$^{G.s3h2.02}$ |
| Gα-K349$^{G.H5.21}$ | MOR-V262$^{5x68}$ |
| MOR-M99$^{12x50}$ | MOR-N342$^{8x49}$ |
| Gα-A41$^{G.s1h1.02}$ | Gα-Q204$^{G.s3h2.03}$ |
| Gα-T181$^{G.hfs2.05}$ | Gα-G202$^{G.s3h2.01}$ |
| MOR-K98$^{12x49}$ | MOR-N342$^{8x49}$ |
| MOR-V89$^{1x53}$ | MOR-C346$^{8x53}$ |
| MOR-M99$^{12x50}$ | MOR-C346$^{8x53}$ |
| Gα-I56$^{G.H1.11}$ | Gα-H188$^{G.S2.05}$ |
| MOR-S266 | MOR-D272$^{6x27}$ |
| MOR-L88$^{1x52}$ | MOR-F350$^{8x57}$ |
| Gα-L353$^{G.H5.25}$ | MOR-L259$^{5x65}$ |

| MSM Contact Features | |
|---|---|
| **Residue i** | **Residue j** |
| Gα-G202$^{G.s3h2.01}$ | Gα-E207$^{G.H2.03}$ |
| Gα-L348$^{G.H5.20}$ | MOR-R263 |
| Gα-L348$^{G.H5.20}$ | MOR-V262$^{5x68}$ |
| MOR-L265 | MOR-D272$^{6x27}$ |
| MOR-V262$^{5x68}$ | MOR-S266 |
| Gα-L353$^{G.H5.25}$ | MOR-R273$^{6x28}$ |
| Gα-I55$^{G.H1.10}$ | Gα-T190$^{G.S2.07}$ |
| Gα-F354$^{G.H5.26}$ | MOR-I256$^{5x62}$ |
| Gα-G203$^{G.s3h2.02}$ | Gα-E207$^{G.H2.03}$ |
| Gα-A71$^{H.HA.09}$ | Gα-A114$^{H.hbhc.02}$ |
| Gα-F354$^{G.H5.26}$ | MOR-R280$^{6x35}$ |
| Gα-G40$^{G.s1h1.01}$ | Gα-G203$^{G.s3h2.02}$ |
| Gα-C325$^{G.s6h5.02}$ | Gα-V332$^{G.H5.04}$ |
| Gα-S206$^{G.H2.02}$ | Gβ-G162 |
| Gα-G183$^{G.hfs2.07}$ | Gα-Q204$^{G.s3h2.03}$ |
| Gα-M53$^{G.H1.08}$ | Gα-E58$^{G.h1ha.01}$ |
| Gα-N347$^{G.H5.19}$ | MOR-M264 |
| MOR-T67$^{1x31}$ | MOR-T312$^{7x28}$ |
| MOR-M264 | MOR-K271$^{6x26}$ |
| Gα-K349$^{G.H5.21}$ | Gα-L353$^{G.H5.25}$ |
| Gα-I56$^{G.H1.11}$ | Gα-T187$^{G.S2.04}$ |
| Gα-L353$^{G.H5.25}$ | MOR-I256$^{5x62}$ |
| Gα-T4$^{G.HN.10}$ | Gα-E8$^{G.HN.32}$ |
| MOR-I107$^{2x43}$ | MOR-A337$^{7x54}$ |
| Gα-T4$^{G.HN.10}$ | Gα-D9$^{G.HN.33}$ |
| Gα-L353$^{G.H5.25}$ | MOR-I278$^{6x33}$ |
| Gα-K345$^{G.H5.17}$ | MOR-M264 |
| MOR-M99$^{12x50}$ | MOR-F343$^{8x50}$ |
| Gα-G42$^{G.s1h1.03}$ | Gα-G203$^{G.s3h2.02}$ |
| Gα-A114$^{H.hbhc.02}$ | Gα-M119$^{H.hbhc.14}$ |
| Gα-V201$^{G.S3.08}$ | Gα-K210$^{G.H2.06}$ |
| Gα-N347$^{G.H5.19}$ | MOR-V262$^{5x68}$ |
| Gα-A59$^{G.h1ha.02}$ | Gα-H188$^{G.S2.05}$ |
| Gα-F354$^{G.H5.26}$ | MOR-L259$^{5x65}$ |
| Gα-I184$^{G.S2.01}$ | Gα-Q204$^{G.s3h2.03}$ |
| Gα-F354$^{G.H5.26}$ | MOR-R276$^{6x31}$ |
| Gα-S206$^{G.H2.02}$ | Gβ-G144 |
| MOR-K98$^{12x49}$ | MOR-C346$^{8x53}$ |
| Gα-A114$^{H.hbhc.02}$ | Gα-T120$^{H.hbhc.15}$ |
| Gα-L353$^{G.H5.25}$ | MOR-R276$^{6x31}$ |
| Gα-F354$^{G.H5.26}$ | MOR-I278$^{6x33}$ |
| Gα-L110$^{H.HB.12}$ | Gα-A114$^{H.hbhc.02}$ |
| Gα-E207$^{G.H2.03}$ | Gβ-D186 |
| Gα-I56$^{G.H1.11}$ | Gα-F189$^{G.S2.06}$ |
| Gα-K180$^{G.hfs2.04}$ | Gα-G203$^{G.s3h2.02}$ |

| **MSM Contact Features** | |
|---|---|
| **Residue i** | **Residue j** |
| MOR-K260$^{5x66}$ | MOR-L275$^{6x30}$ |
| Gα-L353$^{G.H5.25}$ | MOR-N274$^{6x29}$ |
| Gα-S206$^{G.H2.02}$ | Gβ-D163 |
| MOR-T97$^{12x48}$ | MOR-F343$^{8x50}$ |
| Gα-Y74$^{H.HA.12}$ | Gα-A114$^{H.hbhc.02}$ |
| MOR-G131$^{2x67}$ | MOR-G213 |
| MOR-L88$^{1x52}$ | MOR-F347$^{8x54}$ |
| Gα-I55$^{G.H1.10}$ | Gα-T329$^{G.H5.01}$ |
| Gα-T324$^{G.s6h5.01}$ | Gα-V335$^{G.H5.07}$ |
| Gα-I55$^{G.H1.10}$ | Gα-F191$^{G.S2.08}$ |
| Gα-V201$^{G.S3.08}$ | Gα-E207$^{G.H2.03}$ |
| MOR-I93$^{1x57}$ | MOR-K100$^{12x51}$ |
| MOR-I69$^{1x33}$ | MOR-L129$^{2x65}$ |
| MOR-V89$^{1x53}$ | MOR-M99$^{12x50}$ |
| Gα-E43$^{G.s1h1.04}$ | Gα-G203$^{G.s3h2.02}$ |
| MOR-S64 | MOR-L129$^{2x65}$ |
| Gα-G112$^{H.HB.14}$ | Gα-F118$^{H.hbhc.13}$ |
| Gα-S206$^{G.H2.02}$ | Gβ-Y145 |
| MOR-V92$^{1x56}$ | MOR-C346$^{8x53}$ |
| Gα-G203$^{G.s3h2.02}$ | Gα-R208$^{G.H2.04}$ |
| Gα-T182$^{G.hfs2.06}$ | Gα-Q204$^{G.s3h2.03}$ |
| Gα-Y61$^{G.h1ha.04}$ | Gα-E65$^{H.HA.03}$ |
| Gα-S206$^{G.H2.02}$ | Gβ-D186 |
| MOR-S261$^{5x67}$ | MOR-L265 |
| Gα-F354$^{G.H5.26}$ | MOR-L275$^{6x30}$ |
| Gα-H213$^{G.H2.09}$ | Gα-W258$^{G.H3.17}$ |
| Gα-I55$^{G.H1.10}$ | Gα-F189$^{G.S2.06}$ |

**Supplementary Table 3. Median stationary populations with 95% confidence intervals.**

| System | GTP-primed | G-ACT-1 | I1 | I2 | I3 | I4 | G-ACT-2/3 |
|---|---|---|---|---|---|---|---|
| LFT-MOR-$G_{i1}$ | 8.05% [1.65, 17.23] | 47.65% [10.68, 67.72] | 7.33% [2.57, 29.60] | 12.79% [5.70, 23.36] | 4.82% [0.89, 22.82] | 8.16% [1.39, 39.28] | 2.24% [0.15, 18.03] |
| MP-MOR-$G_{i1}$ | 7.32% [2.08, 14.45] | 24.70% [6.52, 44.64] | 22.22% [3.08, 48.57] | 9.60% [2.36, 27.65] | 11.28% [1.10, 57.77] | 4.34% [0.46, 35.94] | 5.41% [1.11, 34.04] |

**Supplementary Table 4. TPT pathways and statistics across bootstrapped MSMs.** TPT pathways and associated statistics include the fraction of bootstrapped models in which each pathway is observed (Model Frac.), the fraction of total reactive flux accounted for by each pathway across all models (Flux Frac.), and the median pathway mean first-passage time with 95% confidence intervals (Med. MFPT [95% CI]).

| **System** | **Path** | **Model frac.** | **Flux frac.** | **Med. MFPT [95% CI] (ms)** |
|---|---|---|---|---|
| LFT-MOR-$G_{i1}$ | GTP-primed → G-ACT-1 → I1 → I2 → I3 → I4 → G-ACT-2/3 | 1.00 | 0.986 | 2.61 [0.97,12.95] |
| LFT-MOR-$G_{i1}$ | GTP-primed → G-ACT-1 → I2 → I3 → I4 → G-ACT-2/3 | 0.48 | 0.014 | 144.04 [19.23,886.19] |
| MP-MOR-$G_{i1}$ | GTP-primed → G-ACT-1 → I1 → I2 → I3 → G-ACT-2/3 | 0.88 | 0.681 | 18.22 [4.86, 69.79] |
| MP-MOR-$G_{i1}$ | GTP-primed → G-ACT-1 → I1 → I2 → I3 → I4 → G-ACT-2/3 | 0.44 | 0.127 | 54.89 [14.27, 411.49] |
| MP-MOR-$G_{i1}$ | GTP-primed → G-ACT-1 → I2 → I3 → I4 → G-ACT-2/3 | 0.49 | 0.105 | 61.24 [19.08, 424.68] |
| MP-MOR-$G_{i1}$ | GTP-primed → G-ACT-1 → I2 → I3 → G-ACT-2/3 | 0.67 | 0.087 | 102.52 [28.20, 1254.87] |

**Supplementary Table 5. Median dwell times with 95% confidence intervals.**

| System | GTP-primed | G-ACT-1 | I1 | I2 | I3 | I4 | G-ACT-2/3 |
|---|---|---|---|---|---|---|---|
| LFT-MOR-$G_{i1}$ | 1.48 µs [1.17, 1.77] | 3.91 µs [3.50, 4.56] | 0.72 µs [0.66, 0.77] | 1.36 µs [1.25, 1.48] | 4.43 µs [3.68, 5.41] | 3.28 µs [2.93, 3.59] | 0.99 µs [0.86, 1.09] |
| MP-MOR-$G_{i1}$ | 1.13 µs [1.01, 1.31] | 1.14 µs [0.98, 1.30] | 0.94 µs [0.85, 0.96] | 2.59 µs [1.98, 3.29] | 2.47 µs [2.32, 2.95] | 0.89 µs [0.79, 0.99] | 1.24 µs [1.14, 1.45] |

**Supplementary Table 6. Median Macrostate MFPTs for Bootstrapped MSMs in units of milliseconds with 95% confidence intervals.**

### LFT-MOR-$G_{i1}$

| i →j | GTP-primed | G-ACT-1 | I1 | I2 | I3 | I4 | G-ACT-2/3 |
|---|---|---|---|---|---|---|---|
| **GTP-primed** | — | 0.002 [0.000, 0.010] | 0.125 [0.057, 1.521] | 0.130 [0.063, 1.555] | 0.521 [0.190, 4.742] | 1.702 [0.364, 9.273] | 1.930 [0.438, 10.076] |
| **G-ACT-1** | 0.049 [0.017, 1.024] | — | 0.104 [0.046, 1.353] | 0.112 [0.050, 1.386] | 0.468 [0.173, 4.718] | 1.686 [0.354, 9.231] | 1.873 [0.417, 10.033] |
| **I1** | 0.249 [0.063, 6.069] | 0.012 [0.005, 0.064] | — | 0.005 [0.002, 0.016] | 0.185 [0.043, 4.534] | 1.306 [0.176, 9.039] | 1.484 [0.195, 9.842] |
| **I2** | 0.208 [0.058, 5.975] | 0.014 [0.006, 0.052] | 0.011 [0.004, 0.029] | — | 0.239 [0.057, 4.587] | 1.424 [0.198, 9.069] | 1.557 [0.216, 9.872] |
| **I3** | 0.405 [0.161, 6.548] | 0.180 [0.056, 0.718] | 0.160 [0.048, 0.668] | 0.155 [0.046, 0.654] | — | 0.342 [0.026, 4.261] | 0.507 [0.053, 5.792] |
| **I4** | 0.484 [0.221, 6.613] | 0.258 [0.106, 0.888] | 0.241 [0.100, 0.870] | 0.236 [0.096, 0.862] | 0.032 [0.011, 0.154] | — | 0.067 [0.010, 0.718] |
| **G-ACT-2/3** | 0.489 [0.242, 6.636] | 0.273 [0.110, 1.636] | 0.257 [0.104, 1.610] | 0.248 [0.101, 1.576] | 0.055 [0.013, 0.537] | 0.003 [0.001, 0.010] | — |

### MP-MOR-$G_{i1}$

| i →j | GTP-primed | G-ACT-1 | I1 | I2 | I3 | I4 | G-ACT-2/3 |
|---|---|---|---|---|---|---|---|
| **GTP-primed** | — | 0.002 [0.001, 0.007] | 0.006 [0.003, 0.020] | 0.096 [0.035, 0.290] | 0.330 [0.107, 0.927] | 10.832 [2.282, 41.823] | 10.736 [2.288, 41.657] |
| **G-ACT-1** | 0.035 [0.011, 0.147] | — | 0.006 [0.002, 0.020] | 0.095 [0.036, 0.281] | 0.330 [0.105, 0.928] | 10.830 [2.282, 41.819] | 10.734 [2.288, 41.653] |
| **I1** | 0.053 [0.021, 0.228] | 0.007 [0.002, 0.068] | — | 0.099 [0.031, 0.291] | 0.330 [0.100, 0.930] | 10.828 [2.286, 41.785] | 10.732 [2.293, 41.620] |
| **I2** | 0.360 [0.059, 2.127] | 0.317 [0.040, 2.010] | 0.297 [0.027, 1.934] | — | 0.119 [0.023, 0.669] | 11.325 [2.213, 42.757] | 11.237 [2.219, 42.592] |
| **I3** | 1.168 [0.324, 25.291] | 1.131 [0.279, 25.183] | 1.122 [0.272, 25.102] | 0.648 [0.004, 24.818] | — | 5.919 [0.374, 37.872] | 5.983 [0.437, 37.752] |
| **I4** | 1.790 [0.426, 30.603] | 1.726 [0.420, 30.560] | 1.699 [0.414, 30.491] | 1.655 [0.364, 30.170] | 0.059 [0.016, 1.110] | — | 0.029 [0.003, 0.260] |
| **G-ACT-2/3** | 1.808 [0.452, 30.616] | 1.738 [0.397, 30.573] | 1.716 [0.373, 30.504] | 1.660 [0.343, 30.184] | 0.100 [0.029, 1.186] | 0.028 [0.001, 0.203] | — |

**Supplementary Table 7. Composition of the simulated systems.** Components of the lipid bilayer are: 1-palmitoyl-2-oleoyl-sn-glycero-3-phosphocholine (POPC), 1-palmitoyl-2-oleoyl-sn-glycero-3-phosphoethanolamine (POPE), 1-palmitoyl-2-oleoyl-sn-glycero-3-phosphoserine (POPS), palmitoyl sphingomyelin (PSM), monosialodihexosylganglioside (GM3), cholesterol, 1,2-diacyl-sn-glycero-3-phospho-1-D-myo-inositol 4,5-bisphosphate (PIP2).

**LFT-bound MOR-$G_{i1}$**

| | GTP-primed | G-ACT-1 | G-ACT-2 | G-ACT-3 |
|---|---|---|---|---|
| Simulation Box Dimensions (Å) | 146 × 146 × 188 | 146 × 146 × 183 | 137 × 137 × 218 | 137 × 137 × 214 |
| Total Number of Atoms | 413,552 | 405,693 | 423,634 | 415,920 |
| LFT-MOR-Gi1 | | | | |
| Water Molecules | 100,482 | 97,867 | 103,836 | 101,270 |
| Salt Concentration | 0.15 M | 0.15 M | 0.15 M | 0.15 M |
| Cholesterol | 192 | 192 | 192 | 192 |
| POPC | 200 | 200 | 200 | 200 |
| POPE | 200 | 200 | 200 | 200 |
| POPS | 60 | 60 | 60 | 60 |
| PIP2 | 40 | 40 | 40 | 40 |
| PSM | 60 | 60 | 60 | 60 |
| GM3 | 40 | 40 | 40 | 40 |

**MP-bound MOR-$G_{i1}$**

| | GTP-primed | G-ACT-1 | G-ACT-2 | G-ACT-3 |
|---|---|---|---|---|
| Simulation Box Dimensions (Å) | 149 × 149 × 177 | 147 × 147 × 188 | 137 × 137 × 216 | 136 × 136 × 212 |
| Total Number of Atoms | 405,199 | 416,273 | 418,898 | 406,794 |
| MP-MOR-Gi1 | | | | |
| Water Molecules | 97,679 | 101,363 | 102,260 | 98,234 |
| Salt Concentration | 0.15 M | 0.15 M | 0.15 M | 0.15 M |
| Cholesterol | 193 | 193 | 192 | 192 |
| POPC | 200 | 200 | 200 | 200 |
| POPE | 200 | 200 | 200 | 200 |
| POPS | 60 | 60 | 60 | 60 |
| PIP2 | 40 | 40 | 40 | 40 |
| PSM | 60 | 60 | 60 | 60 |
| GM3 | 40 | 40 | 40 | 40 |

**Supplementary Table 8. MD equilibration schedule.**

| | | Position Restraints | | | |
|---|---|---|---|---|---|
| Integration step, fs | Simulation time, ps | Backbone kJ/(mol*nm$^2$) | Side chain kJ/(mol*nm$^2$) | Lipid (z axis) kJ/(mol*nm$^2$) | Dihedral kJ/(mol*rad$^2$) |
| 0.5 | 250 | 4,000 | 2,000 | 1,000 | 1,000 |
| 1 | 125 | 2,000 | 1,000 | 400 | 500 |
| 1 | 125 | 2,000 | 1,000 | 200 | 500 |
| 2 | 500 | 1,000 | 500 | 200 | 200 |
| 2 | 500 | 200 | 50 | 40 | 100 |
| 2 | 500 | 50 | 0 | 0 | 0 |
| 4 | 200,000 | 50 | 0 | 0 | 0 |

**Supplementary Table 9. List of the 685 Cα–Cα distance features spanning key structural elements of MOR and Gα used for adaptive sampling.**

| MSM Contact Features | |
|---|---|
| **Residue i** | **Residue j** |
| Gα-A101$^{H.HB.03}$ | Gα-L130$^{H.HC.10}$ |
| Gα-A104$^{H.HB.06}$ | Gα-L130$^{H.HC.10}$ |
| Gα-A111$^{H.HB.13}$ | Gα-E122$^{H.HC.02}$ |
| Gα-A111$^{H.HB.13}$ | Gα-F118$^{H.hbhc.13}$ |
| Gα-A111$^{H.HB.13}$ | Gα-T120$^{H.hbhc.15}$ |
| Gα-A113$^{H.hbhc.01}$ | Gα-F118$^{H.hbhc.13}$ |
| Gα-A114$^{H.hbhc.02}$ | Gα-F118$^{H.hbhc.13}$ |
| Gα-A114$^{H.hbhc.02}$ | Gα-T120$^{H.hbhc.15}$ |
| Gα-A163$^{H.HE.13}$ | Gα-Y167$^{H.hehf.04}$ |
| Gα-A220$^{G.S4.01}$ | Gα-I343$^{G.H5.15}$ |
| Gα-A235$^{G.s4h3.09}$ | Gα-E239$^{G.s4h3.13}$ |
| Gα-A30$^{G.hns1.01}$ | MOR-S266 |
| Gα-A31$^{G.hns1.02}$ | MOR-L176$^{34x54}$ |
| Gα-A31$^{G.hns1.02}$ | MOR-L265 |
| Gα-A31$^{G.hns1.02}$ | MOR-M264 |
| Gα-A31$^{G.hns1.02}$ | MOR-S266 |
| Gα-A326$^{G.s6h5.03}$ | Gα-V332$^{G.H5.04}$ |
| Gα-A41$^{G.s1h1.02}$ | Gα-R208$^{G.H2.04}$ |
| Gα-A41$^{G.s1h1.02}$ | Gα-Q204$^{G.s3h2.03}$ |
| Gα-A41$^{G.s1h1.02}$ | Gα-G203$^{G.s3h2.02}$ |
| Gα-A59$^{G.h1ha.02}$ | Gα-R178$^{G.hfs2.02}$ |
| Gα-A59$^{G.h1ha.02}$ | Gα-H188$^{G.S2.05}$ |
| Gα-A71$^{H.HA.09}$ | Gβ-R134 |
| Gα-A71$^{H.HA.09}$ | Gβ-V135 |
| Gα-A87$^{H.HA.25}$ | Gα-I93$^{H.hahb.04}$ |
| Gα-R100$^{H.HB.02}$ | Gα-L130$^{H.HC.10}$ |
| Gα-R144$^{H.HD.11}$ | Gα-D231$^{G.s4h3.05}$ |
| Gα-R144$^{H.HD.11}$ | Gα-E238$^{G.s4h3.12}$ |
| Gα-R144$^{H.HD.11}$ | Gα-I285$^{G.HG.17}$ |
| Gα-R144$^{H.HD.11}$ | Gα-L232$^{G.s4h3.06}$ |
| Gα-R144$^{H.HD.11}$ | Gα-K277$^{G.HG.07}$ |
| Gα-R144$^{H.HD.11}$ | Gα-P282$^{G.HG.14}$ |
| Gα-R144$^{H.HD.11}$ | Gα-S281$^{G.HG.13}$ |
| Gα-R144$^{H.HD.11}$ | Gα-Y230$^{G.s4h3.04}$ |
| Gα-R144$^{H.HD.11}$ | Gα-V233$^{G.s4h3.07}$ |
| Gα-R15$^{G.HN.39}$ | Gβ-G131 |
| Gα-R161$^{H.HE.11}$ | Gα-I168$^{H.hehf.05}$ |

| | |
|---|---|
| Gα-R176$^{H.HF.06}$ | Gα-D272$^{G.HG.02}$ |
| Gα-R176$^{H.HF.06}$ | Gα-T327$^{G.s6h5.04}$ |
| Gα-R178$^{G.hfs2.02}$ | Gβ-R96 |
| Gα-R178$^{G.hfs2.02}$ | Gβ-E138 |
| Gα-R205$^{G.H2.01}$ | Gα-D237$^{G.s4h3.11}$ |
| Gα-R205$^{G.H2.01}$ | Gα-E236$^{G.s4h3.10}$ |
| Gα-R205$^{G.H2.01}$ | Gα-E245$^{G.H3.04}$ |
| Gα-R205$^{G.H2.01}$ | Gα-L234$^{G.s4h3.08}$ |
| Gα-R205$^{G.H2.01}$ | Gα-K209$^{G.H2.05}$ |
| Gα-R205$^{G.H2.01}$ | Gα-M240$^{G.s4h3.14}$ |
| Gα-R208$^{G.H2.04}$ | Gα-E236$^{G.s4h3.10}$ |
| Gα-R208$^{G.H2.04}$ | Gα-E245$^{G.H3.04}$ |
| Gα-R208$^{G.H2.04}$ | Gα-I212$^{G.H2.08}$ |
| Gα-R24$^{G.HN.48}$ | MOR-A175$^{34x53}$ |
| Gα-R24$^{G.HN.48}$ | MOR-R182$^{4x40}$ |
| Gα-R24$^{G.HN.48}$ | MOR-D177$^{34x55}$ |
| Gα-R24$^{G.HN.48}$ | MOR-L176$^{34x54}$ |
| Gα-R24$^{G.HN.48}$ | MOR-P172$^{34x50}$ |
| Gα-R24$^{G.HN.48}$ | MOR-V173$^{34x51}$ |
| Gα-R313$^{G.h4s6.04}$ | Gα-K345$^{G.H5.17}$ |
| Gα-R32$^{G.hns1.03}$ | MOR-D177$^{34x55}$ |
| Gα-R32$^{G.hns1.03}$ | MOR-L176$^{34x54}$ |
| Gα-R32$^{G.hns1.03}$ | MOR-L265 |
| Gα-R32$^{G.hns1.03}$ | MOR-M264 |
| Gα-R32$^{G.hns1.03}$ | MOR-S266 |
| Gα-R90$^{H.HA.28}$ | Gα-Y146$^{H.hdhe.01}$ |
| Gα-N141$^{H.HD.08}$ | Gα-K280$^{G.HG.12}$ |
| Gα-N149$^{H.hdhe.04}$ | Gα-R178$^{G.hfs2.02}$ |
| Gα-N311$^{G.h4s6.02}$ | Gα-E318$^{G.h4s6.12}$ |
| Gα-N347$^{G.H5.19}$ | MOR-A168$^{3x53}$ |
| Gα-N347$^{G.H5.19}$ | MOR-A175$^{34x53}$ |
| Gα-N347$^{G.H5.19}$ | MOR-R179$^{34x57}$ |
| Gα-N347$^{G.H5.19}$ | MOR-L176$^{34x54}$ |
| Gα-N347$^{G.H5.19}$ | MOR-P172$^{34x50}$ |
| Gα-N347$^{G.H5.19}$ | MOR-V169$^{3x54}$ |
| Gα-N347$^{G.H5.19}$ | MOR-V262$^{5x68}$ |
| Gα-N76$^{H.HA.14}$ | Gα-R178$^{G.hfs2.02}$ |
| Gα-D103$^{H.HB.05}$ | Gα-E122$^{H.HC.02}$ |
| Gα-D103$^{H.HB.05}$ | Gα-L130$^{H.HC.10}$ |
| Gα-D150$^{H.hdhe.05}$ | Gα-D229$^{G.s4h3.03}$ |
| Gα-D150$^{H.hdhe.05}$ | Gα-L232$^{G.s4h3.06}$ |

| | |
|---|---|
| Gα-D150$^{H.hdhe.05}$ | Gα-L273$^{G.HG.03}$ |
| Gα-D150$^{H.hdhe.05}$ | Gα-K270$^{G.s5hg.01}$ |
| Gα-D150$^{H.hdhe.05}$ | Gα-S228$^{G.s4h3.02}$ |
| Gα-D193$^{G.s2s3.02}$ | MOR-D177$^{34x55}$ |
| Gα-D193$^{G.s2s3.02}$ | MOR-V173$^{34x51}$ |
| Gα-D20$^{G.HN.44}$ | MOR-L176$^{34x54}$ |
| Gα-D231$^{G.s4h3.05}$ | Gα-I278$^{G.HG.08}$ |
| Gα-D261$^{G.h3s5.02}$ | Gα-R313$^{G.h4s6.04}$ |
| Gα-D261$^{G.h3s5.02}$ | Gα-K317$^{G.h4s6.11}$ |
| Gα-D261$^{G.h3s5.02}$ | Gα-K345$^{G.H5.17}$ |
| Gα-D261$^{G.h3s5.02}$ | Gα-T316$^{G.h4s6.10}$ |
| Gα-D261$^{G.h3s5.02}$ | MOR-R273$^{6x28}$ |
| Gα-D315$^{G.h4s6.09}$ | MOR-E270$^{6x25}$ |
| Gα-D315$^{G.h4s6.09}$ | MOR-K271$^{6x26}$ |
| Gα-D315$^{G.h4s6.09}$ | MOR-S268$^{6x23}$ |
| Gα-D328$^{G.s6h5.05}$ | Gα-Q333$^{G.H5.05}$ |
| Gα-D328$^{G.s6h5.05}$ | Gα-V332$^{G.H5.04}$ |
| Gα-D341$^{G.H5.13}$ | MOR-R258$^{5x64}$ |
| Gα-D341$^{G.H5.13}$ | MOR-R263 |
| Gα-D341$^{G.H5.13}$ | MOR-M264 |
| Gα-D341$^{G.H5.13}$ | MOR-V262$^{5x68}$ |
| Gα-D350$^{G.H5.22}$ | MOR-R179$^{34x57}$ |
| Gα-D350$^{G.H5.22}$ | MOR-R263 |
| Gα-D350$^{G.H5.22}$ | MOR-R276$^{6x31}$ |
| Gα-D350$^{G.H5.22}$ | MOR-N342$^{8x49}$ |
| Gα-D350$^{G.H5.22}$ | MOR-K269$^{6x24}$ |
| Gα-D350$^{G.H5.22}$ | MOR-T101$^{2x37}$ |
| Gα-D350$^{G.H5.22}$ | MOR-T103$^{2x39}$ |
| Gα-D350$^{G.H5.22}$ | MOR-V262$^{5x68}$ |
| Gα-C214$^{G.H2.10}$ | Gβ-M101 |
| Gα-C214$^{G.H2.10}$ | Gβ-Y59 |
| Gα-C214$^{G.H2.10}$ | Gα-V218$^{G.h2s4.04}$ |
| Gα-C325$^{G.s6h5.02}$ | Gα-N331$^{G.H5.03}$ |
| Gα-C325$^{G.s6h5.02}$ | Gα-V332$^{G.H5.04}$ |
| Gα-C351$^{G.H5.23}$ | MOR-A168$^{3x53}$ |
| Gα-C351$^{G.H5.23}$ | MOR-R165$^{3x50}$ |
| Gα-C351$^{G.H5.23}$ | MOR-R179$^{34x57}$ |
| Gα-C351$^{G.H5.23}$ | MOR-R263 |
| Gα-C351$^{G.H5.23}$ | MOR-R276$^{6x31}$ |
| Gα-C351$^{G.H5.23}$ | MOR-N342$^{8x49}$ |
| Gα-C351$^{G.H5.23}$ | MOR-D164$^{3x49}$ |

| | |
|---|---|
| Gα-C351$^{G.H5.23}$ | MOR-D340$^{8x47}$ |
| Gα-C351$^{G.H5.23}$ | MOR-L259$^{5x65}$ |
| Gα-C351$^{G.H5.23}$ | MOR-K260$^{5x66}$ |
| Gα-C351$^{G.H5.23}$ | MOR-S261$^{5x67}$ |
| Gα-C351$^{G.H5.23}$ | MOR-T103$^{2x39}$ |
| Gα-C351$^{G.H5.23}$ | MOR-V262$^{5x68}$ |
| Gα-C66$^{H.HA.04}$ | Gα-N166$^{H.hehf.03}$ |
| Gα-C66$^{H.HA.04}$ | Gα-Q171$^{H.HF.01}$ |
| Gα-C66$^{H.HA.04}$ | Gα-T170$^{H.hehf.07}$ |
| Gα-C66$^{H.HA.04}$ | Gα-Y167$^{H.hehf.04}$ |
| Gα-C66$^{H.HA.04}$ | Gα-V174$^{H.HF.04}$ |
| Gα-Q147$^{H.hdhe.02}$ | Gα-A235$^{G.s4h3.09}$ |
| Gα-Q147$^{H.hdhe.02}$ | Gα-R178$^{G.hfs2.02}$ |
| Gα-Q147$^{H.hdhe.02}$ | Gα-V233$^{G.s4h3.07}$ |
| Gα-Q164$^{H.hehf.01}$ | Gα-I168$^{H.hehf.05}$ |
| Gα-Q171$^{H.HF.01}$ | Gα-T329$^{G.H5.01}$ |
| Gα-Q172$^{H.HF.02}$ | Gα-T327$^{G.s6h5.04}$ |
| Gα-Q204$^{G.s3h2.03}$ | Gβ-N119 |
| Gα-Q204$^{G.s3h2.03}$ | Gβ-G144 |
| Gα-Q204$^{G.s3h2.03}$ | Gβ-L117 |
| Gα-Q204$^{G.s3h2.03}$ | Gβ-T143 |
| Gα-Q204$^{G.s3h2.03}$ | Gβ-Y145 |
| Gα-Q204$^{G.s3h2.03}$ | Gα-E236$^{G.s4h3.10}$ |
| Gα-Q52$^{G.H1.07}$ | Gα-N331$^{G.H5.03}$ |
| Gα-Q52$^{G.H1.07}$ | Gα-Q171$^{H.HF.01}$ |
| Gα-Q52$^{G.H1.07}$ | Gα-H57$^{G.H1.12}$ |
| Gα-Q52$^{G.H1.07}$ | Gα-I56$^{G.H1.11}$ |
| Gα-Q52$^{G.H1.07}$ | Gα-L175$^{H.HF.05}$ |
| Gα-Q52$^{G.H1.07}$ | Gα-V332$^{G.H5.04}$ |
| Gα-Q68$^{H.HA.06}$ | Gβ-R134 |
| Gα-Q68$^{H.HA.06}$ | Gβ-N132 |
| Gα-Q68$^{H.HA.06}$ | Gβ-I93 |
| Gα-Q68$^{H.HA.06}$ | Gβ-P94 |
| Gα-Q68$^{H.HA.06}$ | Gβ-Y124 |
| Gα-Q68$^{H.HA.06}$ | Gβ-V133 |
| Gα-Q79$^{H.HA.17}$ | Gβ-R137 |
| Gα-Q79$^{H.HA.17}$ | Gα-R178$^{G.hfs2.02}$ |
| Gα-E115$^{H.hbhc.03}$ | Gβ-R129 |
| Gα-E116$^{H.hbhc.04}$ | Gβ-R129 |
| Gα-E116$^{H.hbhc.04}$ | Gβ-R134 |
| Gα-E116$^{H.hbhc.04}$ | Gβ-N125 |

| | |
|---|---|
| Gα-E116$^{H.hbhc.04}$ | Gβ-T128 |
| Gα-E145$^{H.HD.12}$ | Gα-A235$^{G.s4h3.09}$ |
| Gα-E145$^{H.HD.12}$ | Gα-R242$^{G.H3.01}$ |
| Gα-E145$^{H.HD.12}$ | Gα-E238$^{G.s4h3.12}$ |
| Gα-E145$^{H.HD.12}$ | Gα-L232$^{G.s4h3.06}$ |
| Gα-E145$^{H.HD.12}$ | Gα-L234$^{G.s4h3.08}$ |
| Gα-E145$^{H.HD.12}$ | Gα-V233$^{G.s4h3.07}$ |
| Gα-E207$^{G.H2.03}$ | Gβ-D186 |
| Gα-E207$^{G.H2.03}$ | Gβ-D228 |
| Gα-E207$^{G.H2.03}$ | Gβ-C204 |
| Gα-E207$^{G.H2.03}$ | Gβ-Y145 |
| Gα-E216$^{G.h2s4.02}$ | Gα-K257$^{G.H3.16}$ |
| Gα-E216$^{G.h2s4.02}$ | Gα-F259$^{G.H3.18}$ |
| Gα-E216$^{G.h2s4.02}$ | MOR-G267 |
| Gα-E216$^{G.h2s4.02}$ | MOR-S268$^{6x23}$ |
| Gα-E236$^{G.s4h3.10}$ | Gα-M240$^{G.s4h3.14}$ |
| Gα-E239$^{G.s4h3.13}$ | Gα-I285$^{G.HG.17}$ |
| Gα-E28$^{G.HN.52}$ | MOR-V173$^{34x51}$ |
| Gα-E318$^{G.h4s6.12}$ | MOR-R263 |
| Gα-E318$^{G.h4s6.12}$ | MOR-K271$^{6x26}$ |
| Gα-E318$^{G.h4s6.12}$ | MOR-M264 |
| Gα-E33$^{G.S1.01}$ | MOR-L265 |
| Gα-E33$^{G.S1.01}$ | MOR-S266 |
| Gα-E43$^{G.s1h1.04}$ | Gα-R178$^{G.hfs2.02}$ |
| Gα-E43$^{G.s1h1.04}$ | Gα-N149$^{H.hdhe.04}$ |
| Gα-E43$^{G.s1h1.04}$ | Gα-D150$^{H.hdhe.05}$ |
| Gα-E58$^{G.h1ha.01}$ | Gα-H188$^{G.S2.05}$ |
| Gα-E58$^{G.h1ha.01}$ | Gα-L175$^{H.HF.05}$ |
| Gα-E58$^{G.h1ha.01}$ | Gα-F189$^{G.S2.06}$ |
| Gα-E58$^{G.h1ha.01}$ | Gα-T187$^{G.S2.04}$ |
| Gα-E63$^{H.HA.01}$ | Gα-I168$^{H.hehf.05}$ |
| Gα-E63$^{H.HA.01}$ | Gα-K67$^{H.HA.05}$ |
| Gα-E64$^{H.HA.02}$ | Gβ-A92 |
| Gα-E64$^{H.HA.02}$ | Gβ-P94 |
| Gα-E65$^{H.HA.03}$ | Gβ-R96 |
| Gα-E65$^{H.HA.03}$ | Gβ-L95 |
| Gα-E65$^{H.HA.03}$ | Gβ-P94 |
| Gα-E65$^{H.HA.03}$ | Gα-P169$^{H.hehf.06}$ |
| Gα-E65$^{H.HA.03}$ | Gα-Y69$^{H.HA.07}$ |
| Gα-G112$^{H.HB.14}$ | Gα-E116$^{H.hbhc.04}$ |
| Gα-G112$^{H.HB.14}$ | Gα-F118$^{H.hbhc.13}$ |

| | |
|---|---|
| Gα-G117$^{H.hbhc.12}$ | Gβ-R134 |
| Gα-G183$^{G.hfs2.07}$ | Gβ-N119 |
| Gα-G183$^{G.hfs2.07}$ | Gβ-D118 |
| Gα-G183$^{G.hfs2.07}$ | Gα-Q204$^{G.s3h2.03}$ |
| Gα-G183$^{G.hfs2.07}$ | Gα-E207$^{G.H2.03}$ |
| Gα-G183$^{G.hfs2.07}$ | Gα-G202$^{G.s3h2.01}$ |
| Gα-G183$^{G.hfs2.07}$ | Gα-G203$^{G.s3h2.02}$ |
| Gα-G183$^{G.hfs2.07}$ | Gα-K210$^{G.H2.06}$ |
| Gα-G202$^{G.s3h2.01}$ | Gα-R208$^{G.H2.04}$ |
| Gα-G202$^{G.s3h2.01}$ | Gα-E207$^{G.H2.03}$ |
| Gα-G202$^{G.s3h2.01}$ | Gα-W211$^{G.H2.07}$ |
| Gα-G203$^{G.s3h2.02}$ | Gα-R208$^{G.H2.04}$ |
| Gα-G203$^{G.s3h2.02}$ | Gα-E207$^{G.H2.03}$ |
| Gα-G203$^{G.s3h2.02}$ | Gα-W211$^{G.H2.07}$ |
| Gα-G217$^{G.h2s4.03}$ | MOR-G267 |
| Gα-G217$^{G.h2s4.03}$ | MOR-S266 |
| Gα-G217$^{G.h2s4.03}$ | MOR-S268$^{6x23}$ |
| Gα-G292$^{G.hgh4.07}$ | Gα-E298$^{G.H4.05}$ |
| Gα-G352$^{G.H5.24}$ | MOR-R263 |
| Gα-G352$^{G.H5.24}$ | MOR-R276$^{6x31}$ |
| Gα-G352$^{G.H5.24}$ | MOR-R345$^{8x52}$ |
| Gα-G352$^{G.H5.24}$ | MOR-N342$^{8x49}$ |
| Gα-G352$^{G.H5.24}$ | MOR-D272$^{6x27}$ |
| Gα-G352$^{G.H5.24}$ | MOR-D340$^{8x47}$ |
| Gα-G352$^{G.H5.24}$ | MOR-E341$^{8x48}$ |
| Gα-G352$^{G.H5.24}$ | MOR-L259$^{5x65}$ |
| Gα-G352$^{G.H5.24}$ | MOR-L339$^{7x56}$ |
| Gα-G352$^{G.H5.24}$ | MOR-K260$^{5x66}$ |
| Gα-G352$^{G.H5.24}$ | MOR-S261$^{5x67}$ |
| Gα-G40$^{G.s1h1.01}$ | Gα-G203$^{G.s3h2.02}$ |
| Gα-G40$^{G.s1h1.01}$ | Gα-W211$^{G.H2.07}$ |
| Gα-G42$^{G.s1h1.03}$ | Gα-R208$^{G.H2.04}$ |
| Gα-G42$^{G.s1h1.03}$ | Gα-Q204$^{G.s3h2.03}$ |
| Gα-G42$^{G.s1h1.03}$ | Gα-G203$^{G.s3h2.02}$ |
| Gα-G45$^{G.s1h1.06}$ | Gα-A326$^{G.s6h5.03}$ |
| Gα-G60$^{G.h1ha.03}$ | Gα-Q171$^{H.HF.01}$ |
| Gα-G60$^{G.h1ha.03}$ | Gα-E65$^{H.HA.03}$ |
| Gα-G96$^{H.hahb.07}$ | Gα-S134$^{H.HD.01}$ |
| Gα-H213$^{G.H2.09}$ | Gβ-Q75 |
| Gα-H213$^{G.H2.09}$ | Gβ-L117 |
| Gα-H213$^{G.H2.09}$ | Gβ-W99 |

| | |
|---|---|
| Gα-H213$^{G.H2.09}$ | Gα-W258$^{G.H3.17}$ |
| Gα-H322$^{G.S6.04}$ | Gα-A338$^{G.H5.10}$ |
| Gα-H322$^{G.S6.04}$ | Gα-N331$^{G.H5.03}$ |
| Gα-H322$^{G.S6.04}$ | Gα-D328$^{G.s6h5.05}$ |
| Gα-H322$^{G.S6.04}$ | Gα-K330$^{G.H5.02}$ |
| Gα-H322$^{G.S6.04}$ | Gα-F334$^{G.H5.06}$ |
| Gα-H322$^{G.S6.04}$ | Gα-V335$^{G.H5.07}$ |
| Gα-H322$^{G.S6.04}$ | Gα-V339$^{G.H5.11}$ |
| Gα-H57$^{G.H1.12}$ | Gα-Q333$^{G.H5.05}$ |
| Gα-H57$^{G.H1.12}$ | Gα-H188$^{G.S2.05}$ |
| Gα-H57$^{G.H1.12}$ | Gα-F191$^{G.S2.08}$ |
| Gα-H57$^{G.H1.12}$ | Gα-F336$^{G.H5.08}$ |
| Gα-H57$^{G.H1.12}$ | Gα-T329$^{G.H5.01}$ |
| Gα-H57$^{G.H1.12}$ | Gα-V332$^{G.H5.04}$ |
| Gα-I184$^{G.S2.01}$ | Gβ-D118 |
| Gα-I184$^{G.S2.01}$ | Gα-C214$^{G.H2.10}$ |
| Gα-I184$^{G.S2.01}$ | Gα-Q204$^{G.s3h2.03}$ |
| Gα-I184$^{G.S2.01}$ | Gα-G202$^{G.s3h2.01}$ |
| Gα-I212$^{G.H2.08}$ | Gβ-W332 |
| Gα-I222$^{G.S4.03}$ | Gα-V335$^{G.H5.07}$ |
| Gα-I222$^{G.S4.03}$ | Gα-V339$^{G.H5.11}$ |
| Gα-I265$^{G.S5.03}$ | Gα-A338$^{G.H5.10}$ |
| Gα-I265$^{G.S5.03}$ | Gα-I343$^{G.H5.15}$ |
| Gα-I265$^{G.S5.03}$ | Gα-F334$^{G.H5.06}$ |
| Gα-I265$^{G.S5.03}$ | Gα-V335$^{G.H5.07}$ |
| Gα-I265$^{G.S5.03}$ | Gα-V339$^{G.H5.11}$ |
| Gα-I319$^{G.S6.01}$ | MOR-R263 |
| Gα-I343$^{G.H5.15}$ | MOR-L176$^{34x54}$ |
| Gα-I343$^{G.H5.15}$ | MOR-P172$^{34x50}$ |
| Gα-I343$^{G.H5.15}$ | MOR-V173$^{34x51}$ |
| Gα-I344$^{G.H5.16}$ | MOR-A168$^{3x53}$ |
| Gα-I344$^{G.H5.16}$ | MOR-R258$^{5x64}$ |
| Gα-I344$^{G.H5.16}$ | MOR-L259$^{5x65}$ |
| Gα-I344$^{G.H5.16}$ | MOR-P172$^{34x50}$ |
| Gα-I344$^{G.H5.16}$ | MOR-V169$^{3x54}$ |
| Gα-I344$^{G.H5.16}$ | MOR-V262$^{5x68}$ |
| Gα-I49$^{G.H1.04}$ | Gα-A326$^{G.s6h5.03}$ |
| Gα-I49$^{G.H1.04}$ | Gα-V332$^{G.H5.04}$ |
| Gα-I55$^{G.H1.10}$ | Gα-A59$^{G.h1ha.02}$ |
| Gα-I55$^{G.H1.10}$ | Gα-Q171$^{H.HF.01}$ |
| Gα-I55$^{G.H1.10}$ | Gα-G60$^{G.h1ha.03}$ |

| | |
|---|---|
| Gα-I55$^{G.H1.10}$ | Gα-L175$^{H.HF.05}$ |
| Gα-I55$^{G.H1.10}$ | Gα-F189$^{G.S2.06}$ |
| Gα-I55$^{G.H1.10}$ | Gα-T190$^{G.S2.07}$ |
| Gα-I55$^{G.H1.10}$ | Gα-T329$^{G.H5.01}$ |
| Gα-I55$^{G.H1.10}$ | Gα-Y61$^{G.h1ha.04}$ |
| Gα-I56$^{G.H1.11}$ | Gα-Q333$^{G.H5.05}$ |
| Gα-I56$^{G.H1.11}$ | Gα-H188$^{G.S2.05}$ |
| Gα-I56$^{G.H1.11}$ | Gα-F189$^{G.S2.06}$ |
| Gα-I56$^{G.H1.11}$ | Gα-T187$^{G.S2.04}$ |
| Gα-I56$^{G.H1.11}$ | Gα-T329$^{G.H5.01}$ |
| Gα-I56$^{G.H1.11}$ | Gα-V332$^{G.H5.04}$ |
| Gα-I78$^{H.HA.16}$ | Gα-E116$^{H.hbhc.04}$ |
| Gα-I78$^{H.HA.16}$ | Gα-I127$^{H.HC.07}$ |
| Gα-I81$^{H.HA.19}$ | Gα-L123$^{H.HC.03}$ |
| Gα-I81$^{H.HA.19}$ | Gα-W131$^{H.HC.11}$ |
| Gα-I81$^{H.HA.19}$ | Gα-V126$^{H.HC.06}$ |
| Gα-I81$^{H.HA.19}$ | Gα-V136$^{H.HD.03}$ |
| Gα-I84$^{H.HA.22}$ | Gα-C139$^{H.HD.06}$ |
| Gα-I84$^{H.HA.22}$ | Gα-L130$^{H.HC.10}$ |
| Gα-I84$^{H.HA.22}$ | Gα-F108$^{H.HB.10}$ |
| Gα-I84$^{H.HA.22}$ | Gα-F140$^{H.HD.07}$ |
| Gα-I84$^{H.HA.22}$ | Gα-V126$^{H.HC.06}$ |
| Gα-I85$^{H.HA.23}$ | Gα-V136$^{H.HD.03}$ |
| Gα-L107$^{H.HB.09}$ | Gα-E122$^{H.HC.02}$ |
| Gα-L107$^{H.HB.09}$ | Gα-I127$^{H.HC.07}$ |
| Gα-L110$^{H.HB.12}$ | Gα-A114$^{H.hbhc.02}$ |
| Gα-L110$^{H.HB.12}$ | Gα-L123$^{H.HC.03}$ |
| Gα-L110$^{H.HB.12}$ | Gα-T120$^{H.hbhc.15}$ |
| Gα-L130$^{H.HC.10}$ | Gα-L156$^{H.HE.06}$ |
| Gα-L148$^{H.hdhe.03}$ | Gα-R178$^{G.hfs2.02}$ |
| Gα-L175$^{H.HF.05}$ | Gβ-R96 |
| Gα-L175$^{H.HF.05}$ | Gα-T327$^{G.s6h5.04}$ |
| Gα-L194$^{G.S3.01}$ | Gα-I344$^{G.H5.16}$ |
| Gα-L194$^{G.S3.01}$ | Gα-F336$^{G.H5.08}$ |
| Gα-L194$^{G.S3.01}$ | Gα-V339$^{G.H5.11}$ |
| Gα-L194$^{G.S3.01}$ | MOR-L176$^{34x54}$ |
| Gα-L194$^{G.S3.01}$ | MOR-L265 |
| Gα-L194$^{G.S3.01}$ | MOR-V173$^{34x51}$ |
| Gα-L232$^{G.s4h3.06}$ | Gα-N241$^{G.s4h3.15}$ |
| Gα-L234$^{G.s4h3.08}$ | Gα-E239$^{G.s4h3.13}$ |
| Gα-L348$^{G.H5.20}$ | Gα-G352$^{G.H5.24}$ |

| | |
|---|---|
| Gα-L348$^{G.H5.20}$ | Gα-L353$^{G.H5.25}$ |
| Gα-L348$^{G.H5.20}$ | Gα-F354$^{G.H5.26}$ |
| Gα-L348$^{G.H5.20}$ | MOR-I278$^{6x33}$ |
| Gα-L348$^{G.H5.20}$ | MOR-L259$^{5x65}$ |
| Gα-L348$^{G.H5.20}$ | MOR-L265 |
| Gα-L348$^{G.H5.20}$ | MOR-M264 |
| Gα-L348$^{G.H5.20}$ | MOR-V169$^{3x54}$ |
| Gα-L348$^{G.H5.20}$ | MOR-V262$^{5x68}$ |
| Gα-L353$^{G.H5.25}$ | MOR-R165$^{3x50}$ |
| Gα-L353$^{G.H5.25}$ | MOR-R258$^{5x64}$ |
| Gα-L353$^{G.H5.25}$ | MOR-R263 |
| Gα-L353$^{G.H5.25}$ | MOR-R276$^{6x31}$ |
| Gα-L353$^{G.H5.25}$ | MOR-R277$^{6x32}$ |
| Gα-L353$^{G.H5.25}$ | MOR-N274$^{6x29}$ |
| Gα-L353$^{G.H5.25}$ | MOR-D272$^{6x27}$ |
| Gα-L353$^{G.H5.25}$ | MOR-D340$^{8x47}$ |
| Gα-L353$^{G.H5.25}$ | MOR-E341$^{8x48}$ |
| Gα-L353$^{G.H5.25}$ | MOR-I256$^{5x62}$ |
| Gα-L353$^{G.H5.25}$ | MOR-I278$^{6x33}$ |
| Gα-L353$^{G.H5.25}$ | MOR-L259$^{5x65}$ |
| Gα-L353$^{G.H5.25}$ | MOR-K260$^{5x66}$ |
| Gα-L353$^{G.H5.25}$ | MOR-M255$^{5x61}$ |
| Gα-L353$^{G.H5.25}$ | MOR-M281$^{6x36}$ |
| Gα-L353$^{G.H5.25}$ | MOR-S261$^{5x67}$ |
| Gα-L353$^{G.H5.25}$ | MOR-V262$^{5x68}$ |
| Gα-L36$^{G.S1.04}$ | Gα-I343$^{G.H5.15}$ |
| Gα-L36$^{G.S1.04}$ | Gα-V335$^{G.H5.07}$ |
| Gα-L37$^{G.S1.05}$ | Gα-C214$^{G.H2.10}$ |
| Gα-L38$^{G.S1.06}$ | Gα-G202$^{G.s3h2.01}$ |
| Gα-L39$^{G.S1.07}$ | Gα-G202$^{G.s3h2.01}$ |
| Gα-L39$^{G.S1.07}$ | Gα-W211$^{G.H2.07}$ |
| Gα-L91$^{H.HA.29}$ | Gα-E145$^{H.HD.12}$ |
| Gα-K17$^{G.HN.41}$ | MOR-D177$^{34x55}$ |
| Gα-K17$^{G.HN.41}$ | MOR-L176$^{34x54}$ |
| Gα-K180$^{G.hfs2.04}$ | Gβ-I120 |
| Gα-K180$^{G.hfs2.04}$ | Gα-Q204$^{G.s3h2.03}$ |
| Gα-K192$^{G.s2s3.01}$ | Gα-D341$^{G.H5.13}$ |
| Gα-K192$^{G.s2s3.01}$ | Gα-Q333$^{G.H5.05}$ |
| Gα-K192$^{G.s2s3.01}$ | Gα-I344$^{G.H5.16}$ |
| Gα-K192$^{G.s2s3.01}$ | Gα-T340$^{G.H5.12}$ |
| Gα-K192$^{G.s2s3.01}$ | MOR-V173$^{34x51}$ |

| | |
|---|---|
| Gα-K209$^{G.H2.05}$ | Gβ-D246 |
| Gα-K210$^{G.H2.06}$ | Gβ-N230 |
| Gα-K210$^{G.H2.06}$ | Gβ-D186 |
| Gα-K210$^{G.H2.06}$ | Gβ-D228 |
| Gα-K210$^{G.H2.06}$ | Gβ-D246 |
| Gα-K210$^{G.H2.06}$ | Gβ-C204 |
| Gα-K210$^{G.H2.06}$ | Gβ-L117 |
| Gα-K210$^{G.H2.06}$ | Gβ-M188 |
| Gα-K271$^{G.HG.01}$ | Gα-N331$^{G.H5.03}$ |
| Gα-K314$^{G.h4s6.08}$ | Gα-E318$^{G.h4s6.12}$ |
| Gα-K314$^{G.h4s6.08}$ | MOR-K271$^{6x26}$ |
| Gα-K317$^{G.h4s6.11}$ | Gα-K345$^{G.H5.17}$ |
| Gα-K345$^{G.H5.17}$ | Gα-F354$^{G.H5.26}$ |
| Gα-K345$^{G.H5.17}$ | MOR-M264 |
| Gα-K349$^{G.H5.21}$ | Gα-L353$^{G.H5.25}$ |
| Gα-K349$^{G.H5.21}$ | Gα-F354$^{G.H5.26}$ |
| Gα-K349$^{G.H5.21}$ | MOR-N342$^{8x49}$ |
| Gα-K35$^{G.S1.03}$ | Gβ-W99 |
| Gα-K35$^{G.S1.03}$ | MOR-S266 |
| Gα-K46$^{G.H1.01}$ | Gα-G202$^{G.s3h2.01}$ |
| Gα-K46$^{G.H1.01}$ | Gα-G203$^{G.s3h2.02}$ |
| Gα-K46$^{G.H1.01}$ | Gα-V201$^{G.S3.08}$ |
| Gα-K51$^{G.H1.06}$ | Gα-L175$^{H.HF.05}$ |
| Gα-K51$^{G.H1.06}$ | Gα-T177$^{G.hfs2.01}$ |
| Gα-K51$^{G.H1.06}$ | Gα-Y61$^{G.h1ha.04}$ |
| Gα-K51$^{G.H1.06}$ | Gα-Y69$^{H.HA.07}$ |
| Gα-K51$^{G.H1.06}$ | Gα-V174$^{H.HF.04}$ |
| Gα-K51$^{G.H1.06}$ | Gα-V179$^{G.hfs2.03}$ |
| Gα-K54$^{G.H1.09}$ | Gα-A59$^{G.h1ha.02}$ |
| Gα-K54$^{G.H1.09}$ | Gα-E58$^{G.h1ha.01}$ |
| Gα-K54$^{G.H1.09}$ | Gα-E65$^{H.HA.03}$ |
| Gα-K54$^{G.H1.09}$ | Gα-G60$^{G.h1ha.03}$ |
| Gα-K54$^{G.H1.09}$ | Gα-F191$^{G.S2.08}$ |
| Gα-K54$^{G.H1.09}$ | Gα-T187$^{G.S2.04}$ |
| Gα-K54$^{G.H1.09}$ | Gα-T190$^{G.S2.07}$ |
| Gα-K54$^{G.H1.09}$ | Gα-Y61$^{G.h1ha.04}$ |
| Gα-K54$^{G.H1.09}$ | Gα-V332$^{G.H5.04}$ |
| Gα-K67$^{H.HA.05}$ | Gβ-R134 |
| Gα-K67$^{H.HA.05}$ | Gβ-N132 |
| Gα-K67$^{H.HA.05}$ | Gβ-E130 |
| Gα-K67$^{H.HA.05}$ | Gβ-V133 |

| | |
|---|---|
| Gα-K67$^{H.HA.05}$ | Gα-A71$^{H.HA.09}$ |
| Gα-K67$^{H.HA.05}$ | Gα-I168$^{H.hehf.05}$ |
| Gα-K67$^{H.HA.05}$ | Gα-P169$^{H.hehf.06}$ |
| Gα-K70$^{H.HA.08}$ | Gβ-R134 |
| Gα-K70$^{H.HA.08}$ | Gα-G117$^{H.hbhc.12}$ |
| Gα-K70$^{H.HA.08}$ | Gα-P165$^{H.hehf.02}$ |
| Gα-M53$^{G.H1.08}$ | Gα-N331$^{G.H5.03}$ |
| Gα-M53$^{G.H1.08}$ | Gα-E58$^{G.h1ha.01}$ |
| Gα-M53$^{G.H1.08}$ | Gα-H57$^{G.H1.12}$ |
| Gα-M53$^{G.H1.08}$ | Gα-F336$^{G.H5.08}$ |
| Gα-M53$^{G.H1.08}$ | Gα-V335$^{G.H5.07}$ |
| Gα-M88$^{H.HA.26}$ | Gα-A104$^{H.HB.06}$ |
| Gα-M88$^{H.HA.26}$ | Gα-D133$^{H.hchd.01}$ |
| Gα-M88$^{H.HA.26}$ | Gα-C139$^{H.HD.06}$ |
| Gα-M88$^{H.HA.26}$ | Gα-F108$^{H.HB.10}$ |
| Gα-M88$^{H.HA.26}$ | Gα-V136$^{H.HD.03}$ |
| Gα-F108$^{H.HB.10}$ | Gα-E122$^{H.HC.02}$ |
| Gα-F108$^{H.HB.10}$ | Gα-G112$^{H.HB.14}$ |
| Gα-F189$^{G.S2.06}$ | Gα-V332$^{G.H5.04}$ |
| Gα-F191$^{G.S2.08}$ | Gα-D337$^{G.H5.09}$ |
| Gα-F191$^{G.S2.08}$ | Gα-T340$^{G.H5.12}$ |
| Gα-F191$^{G.S2.08}$ | Gα-V332$^{G.H5.04}$ |
| Gα-F196$^{G.S3.03}$ | Gα-T340$^{G.H5.12}$ |
| Gα-F196$^{G.S3.03}$ | Gα-V332$^{G.H5.04}$ |
| Gα-F196$^{G.S3.03}$ | Gα-V335$^{G.H5.07}$ |
| Gα-F196$^{G.S3.03}$ | Gα-V339$^{G.H5.11}$ |
| Gα-F199$^{G.S3.06}$ | Gα-C214$^{G.H2.10}$ |
| Gα-F215$^{G.h2s4.01}$ | Gβ-L117 |
| Gα-F215$^{G.h2s4.01}$ | Gβ-W99 |
| Gα-F267$^{G.S5.05}$ | Gα-N331$^{G.H5.03}$ |
| Gα-F267$^{G.S5.05}$ | Gα-F336$^{G.H5.08}$ |
| Gα-F267$^{G.S5.05}$ | Gα-V339$^{G.H5.11}$ |
| Gα-F323$^{G.S6.05}$ | Gα-D328$^{G.s6h5.05}$ |
| Gα-F323$^{G.S6.05}$ | Gα-V335$^{G.H5.07}$ |
| Gα-F336$^{G.H5.08}$ | MOR-V173$^{34x51}$ |
| Gα-F354$^{G.H5.26}$ | MOR-R276$^{6x31}$ |
| Gα-F354$^{G.H5.26}$ | MOR-R277$^{6x32}$ |
| Gα-F354$^{G.H5.26}$ | MOR-R280$^{6x35}$ |
| Gα-F354$^{G.H5.26}$ | MOR-R345$^{8x52}$ |
| Gα-F354$^{G.H5.26}$ | MOR-N274$^{6x29}$ |
| Gα-F354$^{G.H5.26}$ | MOR-E270$^{6x25}$ |

| | |
|---|---|
| Gα-F354$^{G.H5.26}$ | MOR-E341$^{8x48}$ |
| Gα-F354$^{G.H5.26}$ | MOR-I278$^{6x33}$ |
| Gα-F354$^{G.H5.26}$ | MOR-L259$^{5x65}$ |
| Gα-F354$^{G.H5.26}$ | MOR-L265 |
| Gα-F354$^{G.H5.26}$ | MOR-K271$^{6x26}$ |
| Gα-F354$^{G.H5.26}$ | MOR-M264 |
| Gα-F354$^{G.H5.26}$ | MOR-T279$^{6x34}$ |
| Gα-S143$^{H.HD.10}$ | Gα-L232$^{G.s4h3.06}$ |
| Gα-S206$^{G.H2.02}$ | Gβ-D163 |
| Gα-S206$^{G.H2.02}$ | Gβ-D186 |
| Gα-S206$^{G.H2.02}$ | Gβ-G162 |
| Gα-S206$^{G.H2.02}$ | Gβ-Y145 |
| Gα-S206$^{G.H2.02}$ | Gα-W211$^{G.H2.07}$ |
| Gα-S263$^{G.S5.01}$ | Gα-N346$^{G.H5.18}$ |
| Gα-S263$^{G.S5.01}$ | Gα-E318$^{G.h4s6.12}$ |
| Gα-S263$^{G.S5.01}$ | Gα-K317$^{G.h4s6.11}$ |
| Gα-S263$^{G.S5.01}$ | Gα-V342$^{G.H5.14}$ |
| Gα-S263$^{G.S5.01}$ | MOR-M264 |
| Gα-S47$^{G.H1.02}$ | Gα-D200$^{G.S3.07}$ |
| Gα-S47$^{G.H1.02}$ | Gα-T181$^{G.hfs2.05}$ |
| Gα-S47$^{G.H1.02}$ | Gα-V179$^{G.hfs2.03}$ |
| Gα-S62$^{G.h1ha.05}$ | Gα-C66$^{H.HA.04}$ |
| Gα-S62$^{G.h1ha.05}$ | Gα-Q68$^{H.HA.06}$ |
| Gα-S62$^{G.h1ha.05}$ | Gα-Y69$^{H.HA.07}$ |
| Gα-S80$^{H.HA.18}$ | Gα-L130$^{H.HC.10}$ |
| Gα-T177$^{G.hfs2.01}$ | Gβ-R96 |
| Gα-T177$^{G.hfs2.01}$ | Gβ-E138 |
| Gα-T181$^{G.hfs2.05}$ | Gβ-N119 |
| Gα-T181$^{G.hfs2.05}$ | Gβ-D118 |
| Gα-T181$^{G.hfs2.05}$ | Gβ-I120 |
| Gα-T181$^{G.hfs2.05}$ | Gα-R205$^{G.H2.01}$ |
| Gα-T181$^{G.hfs2.05}$ | Gα-D200$^{G.S3.07}$ |
| Gα-T181$^{G.hfs2.05}$ | Gα-Q204$^{G.s3h2.03}$ |
| Gα-T181$^{G.hfs2.05}$ | Gα-E207$^{G.H2.03}$ |
| Gα-T181$^{G.hfs2.05}$ | Gα-G202$^{G.s3h2.01}$ |
| Gα-T181$^{G.hfs2.05}$ | Gα-G203$^{G.s3h2.02}$ |
| Gα-T182$^{G.hfs2.06}$ | Gβ-N119 |
| Gα-T182$^{G.hfs2.06}$ | Gβ-G144 |
| Gα-T182$^{G.hfs2.06}$ | Gβ-T143 |
| Gα-T182$^{G.hfs2.06}$ | Gα-R205$^{G.H2.01}$ |
| Gα-T182$^{G.hfs2.06}$ | Gα-E207$^{G.H2.03}$ |

| | |
|---|---|
| Gα-T182$^{G.hfs2.06}$ | Gα-G202$^{G.s3h2.01}$ |
| Gα-T182$^{G.hfs2.06}$ | Gα-K210$^{G.H2.06}$ |
| Gα-T219$^{G.h2s4.05}$ | Gα-N347$^{G.H5.19}$ |
| Gα-T219$^{G.h2s4.05}$ | Gα-V339$^{G.H5.11}$ |
| Gα-T219$^{G.h2s4.05}$ | Gα-V342$^{G.H5.14}$ |
| Gα-T219$^{G.h2s4.05}$ | MOR-L265 |
| Gα-T219$^{G.h2s4.05}$ | MOR-S266 |
| Gα-T284$^{G.HG.16}$ | Gα-N294$^{G.H4.01}$ |
| Gα-T316$^{G.h4s6.10}$ | Gα-K345$^{G.H5.17}$ |
| Gα-T321$^{G.S6.03}$ | Gα-F334$^{G.H5.06}$ |
| Gα-T324$^{G.s6h5.01}$ | Gα-D328$^{G.s6h5.05}$ |
| Gα-T324$^{G.s6h5.01}$ | Gα-V332$^{G.H5.04}$ |
| Gα-T324$^{G.s6h5.01}$ | Gα-V335$^{G.H5.07}$ |
| Gα-T329$^{G.H5.01}$ | Gα-Q333$^{G.H5.05}$ |
| Gα-T340$^{G.H5.12}$ | MOR-R258$^{5x64}$ |
| Gα-T340$^{G.H5.12}$ | MOR-P172$^{34x50}$ |
| Gα-T340$^{G.H5.12}$ | MOR-V173$^{34x51}$ |
| Gα-T48$^{G.H1.03}$ | Gα-L175$^{H.HF.05}$ |
| Gα-T48$^{G.H1.03}$ | Gα-T327$^{G.s6h5.04}$ |
| Gα-T77$^{H.HA.15}$ | Gα-L123$^{H.HC.03}$ |
| Gα-W211$^{G.H2.07}$ | Gβ-L117 |
| Gα-W211$^{G.H2.07}$ | Gβ-M101 |
| Gα-W211$^{G.H2.07}$ | Gβ-Y145 |
| Gα-W211$^{G.H2.07}$ | Gα-I253$^{G.H3.12}$ |
| Gα-Y155$^{H.HE.05}$ | Gα-R178$^{G.hfs2.02}$ |
| Gα-Y320$^{G.S6.02}$ | Gα-A338$^{G.H5.10}$ |
| Gα-Y320$^{G.S6.02}$ | Gα-N346$^{G.H5.18}$ |
| Gα-Y320$^{G.S6.02}$ | Gα-F334$^{G.H5.06}$ |
| Gα-Y320$^{G.S6.02}$ | Gα-V342$^{G.H5.14}$ |
| Gα-Y320$^{G.S6.02}$ | MOR-R263 |
| Gα-Y320$^{G.S6.02}$ | MOR-M264 |
| Gα-Y61$^{G.h1ha.04}$ | Gβ-R96 |
| Gα-Y61$^{G.h1ha.04}$ | Gα-C66$^{H.HA.04}$ |
| Gα-Y61$^{G.h1ha.04}$ | Gα-E65$^{H.HA.03}$ |
| Gα-Y69$^{H.HA.07}$ | Gβ-R134 |
| Gα-Y69$^{H.HA.07}$ | Gβ-R96 |
| Gα-Y69$^{H.HA.07}$ | Gβ-E138 |
| Gα-Y69$^{H.HA.07}$ | Gβ-V135 |
| Gα-V109$^{H.HB.11}$ | Gα-A113$^{H.hbhc.01}$ |
| Gα-V174$^{H.HF.04}$ | Gβ-R96 |
| Gα-V174$^{H.HF.04}$ | Gα-R178$^{G.hfs2.02}$ |

| | |
|---|---|
| Gα-V179$^{G.hfs2.03}$ | Gβ-A140 |
| Gα-V179$^{G.hfs2.03}$ | Gβ-R96 |
| Gα-V179$^{G.hfs2.03}$ | Gβ-E138 |
| Gα-V179$^{G.hfs2.03}$ | Gβ-I120 |
| Gα-V201$^{G.S3.08}$ | Gα-C214$^{G.H2.10}$ |
| Gα-V218$^{G.h2s4.04}$ | MOR-S266 |
| Gα-V233$^{G.s4h3.07}$ | Gα-D237$^{G.s4h3.11}$ |
| Gα-V233$^{G.s4h3.07}$ | Gα-E238$^{G.s4h3.12}$ |
| Gα-V34$^{G.S1.02}$ | Gα-V339$^{G.H5.11}$ |
| Gα-V34$^{G.S1.02}$ | MOR-L265 |
| Gα-V34$^{G.S1.02}$ | MOR-S266 |
| Gα-V342$^{G.H5.14}$ | MOR-M264 |
| Gα-V50$^{G.H1.05}$ | Gα-K54$^{G.H1.09}$ |
| Gα-V72$^{H.HA.10}$ | Gβ-S136 |
| Gα-V72$^{H.HA.10}$ | Gβ-V135 |
| Gα-V72$^{H.HA.10}$ | Gα-R178$^{G.hfs2.02}$ |
| Gα-V72$^{H.HA.10}$ | Gα-V179$^{G.hfs2.03}$ |
| MOR-A102$^{2x38}$ | MOR-A184$^{4x42}$ |
| MOR-A102$^{2x38}$ | MOR-P181$^{4x39}$ |
| MOR-A102$^{2x38}$ | MOR-T180$^{4x38}$ |
| MOR-A168$^{3x53}$ | MOR-L275$^{6x30}$ |
| MOR-A337$^{7x54}$ | MOR-E341$^{8x48}$ |
| MOR-A337$^{7x54}$ | MOR-K344$^{8x51}$ |
| MOR-A337$^{7x54}$ | MOR-F347$^{8x54}$ |
| MOR-R165$^{3x50}$ | MOR-R276$^{6x31}$ |
| MOR-R165$^{3x50}$ | MOR-N274$^{6x29}$ |
| MOR-R165$^{3x50}$ | MOR-D340$^{8x47}$ |
| MOR-R165$^{3x50}$ | MOR-L275$^{6x30}$ |
| MOR-R165$^{3x50}$ | MOR-M281$^{6x36}$ |
| MOR-R165$^{3x50}$ | MOR-T279$^{6x34}$ |
| MOR-R165$^{3x50}$ | MOR-Y336$^{7x53}$ |
| MOR-R258$^{5x64}$ | MOR-V262$^{5x68}$ |
| MOR-R263 | MOR-R273$^{6x28}$ |
| MOR-R263 | MOR-R276$^{6x31}$ |
| MOR-R263 | MOR-G267 |
| MOR-R263 | MOR-K269$^{6x24}$ |
| MOR-R263 | MOR-K271$^{6x26}$ |
| MOR-R277$^{6x32}$ | MOR-A337$^{7x54}$ |
| MOR-R277$^{6x32}$ | MOR-N342$^{8x49}$ |
| MOR-R277$^{6x32}$ | MOR-D340$^{8x47}$ |
| MOR-R277$^{6x32}$ | MOR-L339$^{7x56}$ |

| | |
|---|---|
| MOR-R277$^{6x32}$ | MOR-F338$^{7x55}$ |
| MOR-R280$^{6x35}$ | MOR-D340$^{8x47}$ |
| MOR-R280$^{6x35}$ | MOR-L339$^{7x56}$ |
| MOR-R95$^{1x59}$ | MOR-F350$^{8x57}$ |
| MOR-N332$^{7x49}$ | MOR-A337$^{7x54}$ |
| MOR-N342$^{8x49}$ | MOR-F347$^{8x54}$ |
| MOR-G267 | MOR-R273$^{6x28}$ |
| MOR-G267 | MOR-D272$^{6x27}$ |
| MOR-G267 | MOR-K271$^{6x26}$ |
| MOR-H171$^{3x56}$ | MOR-R179$^{34x57}$ |
| MOR-I107$^{2x43}$ | MOR-A337$^{7x54}$ |
| MOR-I107$^{2x43}$ | MOR-D340$^{8x47}$ |
| MOR-I107$^{2x43}$ | MOR-M281$^{6x36}$ |
| MOR-I107$^{2x43}$ | MOR-Y336$^{7x53}$ |
| MOR-I155$^{3x40}$ | MOR-V245$^{5x51}$ |
| MOR-I167$^{3x52}$ | MOR-A175$^{34x53}$ |
| MOR-I248$^{5x54}$ | MOR-F289$^{6x44}$ |
| MOR-I256$^{5x62}$ | MOR-I278$^{6x33}$ |
| MOR-I256$^{5x62}$ | MOR-L283$^{6x38}$ |
| MOR-I256$^{5x62}$ | MOR-T279$^{6x34}$ |
| MOR-I256$^{5x62}$ | MOR-V282$^{6x37}$ |
| MOR-I93$^{1x57}$ | MOR-K100$^{12x51}$ |
| MOR-I93$^{1x57}$ | MOR-K98$^{12x49}$ |
| MOR-I93$^{1x57}$ | MOR-T97$^{12x48}$ |
| MOR-L110$^{2x46}$ | MOR-P333$^{7x50}$ |
| MOR-L110$^{2x46}$ | MOR-V285$^{6x40}$ |
| MOR-L158$^{3x43}$ | MOR-N332$^{7x49}$ |
| MOR-L158$^{3x43}$ | MOR-Y336$^{7x53}$ |
| MOR-L158$^{3x43}$ | MOR-V286$^{6x41}$ |
| MOR-L176$^{34x54}$ | MOR-N183$^{4x41}$ |
| MOR-L176$^{34x54}$ | MOR-T180$^{4x38}$ |
| MOR-L259$^{5x65}$ | MOR-R276$^{6x31}$ |
| MOR-L259$^{5x65}$ | MOR-I278$^{6x33}$ |
| MOR-L259$^{5x65}$ | MOR-L265 |
| MOR-L265 | MOR-N274$^{6x29}$ |
| MOR-L265 | MOR-D272$^{6x27}$ |
| MOR-L265 | MOR-I278$^{6x33}$ |
| MOR-L265 | MOR-L275$^{6x30}$ |
| MOR-L265 | MOR-K271$^{6x26}$ |
| MOR-L339$^{7x56}$ | MOR-K344$^{8x51}$ |
| MOR-K100$^{12x51}$ | MOR-P181$^{4x39}$ |

| | |
|---|---|
| MOR-K174$^{34x52}$ | MOR-R179$^{34x57}$ |
| MOR-K174$^{34x52}$ | MOR-N183$^{4x41}$ |
| MOR-K174$^{34x52}$ | MOR-T180$^{4x38}$ |
| MOR-K260$^{5x66}$ | MOR-L265 |
| MOR-K260$^{5x66}$ | MOR-L275$^{6x30}$ |
| MOR-K260$^{5x66}$ | MOR-S266 |
| MOR-K260$^{5x66}$ | MOR-T279$^{6x34}$ |
| MOR-K98$^{12x49}$ | MOR-R345$^{8x52}$ |
| MOR-K98$^{12x49}$ | MOR-N104$^{2x40}$ |
| MOR-K98$^{12x49}$ | MOR-N342$^{8x49}$ |
| MOR-K98$^{12x49}$ | MOR-E341$^{8x48}$ |
| MOR-K98$^{12x49}$ | MOR-K344$^{8x51}$ |
| MOR-K98$^{12x49}$ | MOR-F343$^{8x50}$ |
| MOR-M161$^{3x46}$ | MOR-Y336$^{7x53}$ |
| MOR-M161$^{3x46}$ | MOR-V282$^{6x37}$ |
| MOR-M255$^{5x61}$ | MOR-I278$^{6x33}$ |
| MOR-M255$^{5x61}$ | MOR-V282$^{6x37}$ |
| MOR-M264 | MOR-K269$^{6x24}$ |
| MOR-M264 | MOR-K271$^{6x26}$ |
| MOR-M281$^{6x36}$ | MOR-D340$^{8x47}$ |
| MOR-M281$^{6x36}$ | MOR-F343$^{8x50}$ |
| MOR-M99$^{12x50}$ | MOR-N342$^{8x49}$ |
| MOR-M99$^{12x50}$ | MOR-D340$^{8x47}$ |
| MOR-M99$^{12x50}$ | MOR-I105$^{2x41}$ |
| MOR-M99$^{12x50}$ | MOR-K185$^{4x43}$ |
| MOR-M99$^{12x50}$ | MOR-F108$^{2x44}$ |
| MOR-M99$^{12x50}$ | MOR-F343$^{8x50}$ |
| MOR-F178$^{34x56}$ | MOR-A184$^{4x42}$ |
| MOR-F338$^{7x55}$ | MOR-K344$^{8x51}$ |
| MOR-F338$^{7x55}$ | MOR-F343$^{8x50}$ |
| MOR-F343$^{8x50}$ | MOR-R348$^{8x55}$ |
| MOR-P172$^{34x50}$ | s |
| MOR-P244$^{5x50}$ | MOR-F289$^{6x44}$ |
| MOR-P333$^{7x50}$ | MOR-F338$^{7x55}$ |
| MOR-P333$^{7x50}$ | MOR-F347$^{8x54}$ |
| MOR-S154$^{3x39}$ | MOR-F289$^{6x44}$ |
| MOR-S162$^{3x47}$ | MOR-V282$^{6x37}$ |
| MOR-S162$^{3x47}$ | MOR-V286$^{6x41}$ |
| MOR-S266 | MOR-D272$^{6x27}$ |
| MOR-S266 | MOR-K271$^{6x26}$ |
| MOR-S268$^{6x23}$ | MOR-D272$^{6x27}$ |

| | |
|---|---|
| MOR-T101$^{2x37}$ | MOR-N342$^{8x49}$ |
| MOR-T101$^{2x37}$ | MOR-P181$^{4x39}$ |
| MOR-T103$^{2x39}$ | MOR-R179$^{34x57}$ |
| MOR-T103$^{2x39}$ | MOR-I278$^{6x33}$ |
| MOR-T103$^{2x39}$ | MOR-M281$^{6x36}$ |
| MOR-T103$^{2x39}$ | MOR-V282$^{6x37}$ |
| MOR-T249$^{5x55}$ | MOR-V286$^{6x41}$ |
| MOR-T97$^{12x48}$ | MOR-N104$^{2x40}$ |
| MOR-T97$^{12x48}$ | MOR-N342$^{8x49}$ |
| MOR-T97$^{12x48}$ | MOR-F343$^{8x50}$ |
| MOR-T97$^{12x48}$ | MOR-F347$^{8x54}$ |
| MOR-Y252$^{5x58}$ | MOR-R280$^{6x35}$ |
| MOR-Y252$^{5x58}$ | MOR-Y336$^{7x53}$ |
| MOR-Y252$^{5x58}$ | MOR-V285$^{6x40}$ |
| MOR-Y336$^{7x53}$ | MOR-F343$^{8x50}$ |
| MOR-Y96$^{1x60}$ | MOR-C351$^{8x58}$ |
| MOR-Y96$^{1x60}$ | MOR-K100$^{12x51}$ |
| MOR-Y96$^{1x60}$ | MOR-F347$^{8x54}$ |
| MOR-Y96$^{1x60}$ | MOR-F350$^{8x57}$ |
| MOR-V163$^{3x48}$ | MOR-F178$^{34x56}$ |
| MOR-V169$^{3x54}$ | MOR-L275$^{6x30}$ |
| MOR-V169$^{3x54}$ | MOR-T279$^{6x34}$ |
| MOR-V262$^{5x68}$ | MOR-S266 |
| MOR-V284$^{6x39}$ | MOR-L339$^{7x56}$ |
| MOR-V285$^{6x40}$ | MOR-N332$^{7x49}$ |
| MOR-V285$^{6x40}$ | MOR-Y336$^{7x53}$ |
| MOR-V334$^{7x51}$ | MOR-L339$^{7x56}$ |
| MOR-V89$^{1x53}$ | MOR-A337$^{7x54}$ |
| MOR-V89$^{1x53}$ | MOR-F338$^{7x55}$ |
| MOR-V89$^{1x53}$ | MOR-F343$^{8x50}$ |
| MOR-V89$^{1x53}$ | MOR-F347$^{8x54}$ |
| MOR-V92$^{1x56}$ | MOR-C346$^{8x53}$ |
| MOR-V92$^{1x56}$ | MOR-C351$^{8x58}$ |
